# An antifungal with a novel mechanism of action discovered via resistance gene-guided genome mining

**DOI:** 10.1101/2025.10.09.681488

**Authors:** Bruno Perlatti, Sandeep Vellanki, Yalong Zhang, Yi-Ming Chiang, Yingxia Hu, Mengdi Yuan, Kyle Dunbar, Abigail Fine, Michelle F Grau, Sheena Li, Timothy O’Donnell, Rajani Shenoy, Hongtao Li, Hui Shi, Xia Xu, Zeyu Chen, Tara Arvedson, Yi Tang, Robert A Cramer, Victor Cee, Colin JB Harvey

## Abstract

Invasive fungal infections claim over two million lives annually, a problem exacerbated by rising resistance to current antifungal treatments and an increasing population of immunocompromised individuals. Despite this, antifungal drug development has stagnated, with few novel agents and fewer novel targets explored in recent decades. Here, we validate acetolactate synthase (ALS), an enzyme critical for branched-chain amino acid biosynthesis and absent in humans, as a promising target for new therapeutics. Using resistance gene-guided genome mining, we discovered a biosynthetic gene cluster in *Aspergillus terreus* encoding HB-35018 (**1)**, a novel spiro-*cis*-decalin tetramic acid that potently inhibits ALS. Biochemical and antifungal assays demonstrate that **1** surpasses existing ALS inhibitors in efficacy against *Aspergillus fumigatus* and other pathogenic fungi. Structural studies via cryo-electron microscopy reveal a unique covalent binding interaction between compound **1** and ALS, distinct from known inhibitors and finally, we demonstrate that ALS is essential for the virulence in a mouse model of invasive aspergillosis. These findings position ALS as a promising target for antifungal development and demonstrate the potential of resistance gene-guided genome mining for antifungal discovery.

## Introduction

The need for novel antifungal therapies is urgent. Fungal infections cause an estimated 2.5 million deaths annually^1^, a number that is expected to rise as the number of immunocompromised patients increases and a warming climate extends the reach of pathogenic fungi^2,3^. At least 1.3 million of these deaths are directly attributable to invasive aspergillosis, leading to *Aspergillus fumigatus*, the fungus primarily responsible for this disease, being included in the “critical priority group” of the “WHO fungal priority pathogens list to guide research, development and public health action”^4^. Current antifungal drugs—including the azoles, echinocandins, and polyenes—are decades old, with the efficacy of azoles and echinocandins diminishing as resistance spreads and the toxicity liabilities of the polyenes well known.^5^ Despite the severe impact of fungal infections and the demonstrated need for alternative therapies, development of new antifungals, particularly those with novel mechanisms of action, has been limited in recent years^6,7^.

Natural products have long played a pivotal role in the discovery of antifungal therapies^8^, and resistance-gene guided genome mining has emerged as a promising method for targeted discovery of natural products with specific activities. In fungi, the phenomenon of biosynthetic gene clusters (BGCs) encoding resistance genes homologous to their product’s target has been appreciated for a range of targets, including inhibitors of glycolysis^9^, the proteasome^10^, ergosterol biosynthesis^11–14^, and cyclin dependent kinases^15,16^. Owing to the breadth of biology addressable by this approach, it presents a significant opportunity for the discovery of antifungal agents with novel mechanisms of action.

The ideal antifungal target is essential for the pathogen’s survival but absent in humans, reducing the risk of host toxicity. This constraint is crucial when treating the immunocompromised individuals that most commonly suffer from fungal infections^17^. Given the relatively recent evolutionary divergence of humans and fungi, potential targets meeting this criterion are rare. The biosynthesis of essential amino acids, including branched-chain amino acids (BCAAs, **Figure 1A**), represents a set of such targets. Acetolactate synthase (ALS), the first dedicated enzyme in BCAA biosynthesis has been targeted extensively for agricultural applications with >50 inhibitors developed as commercial herbicides, including the widely used bensulfuron methyl (**2**), penoxsulam (**3**), bispyribac (**4**), and chlormuron ethyl (**5, Figure 1B**).^18^ Despite this effort, to date, the potential for ALS inhibitors as human antifungals remains largely untapped.^19,20^

**Figure 1:**
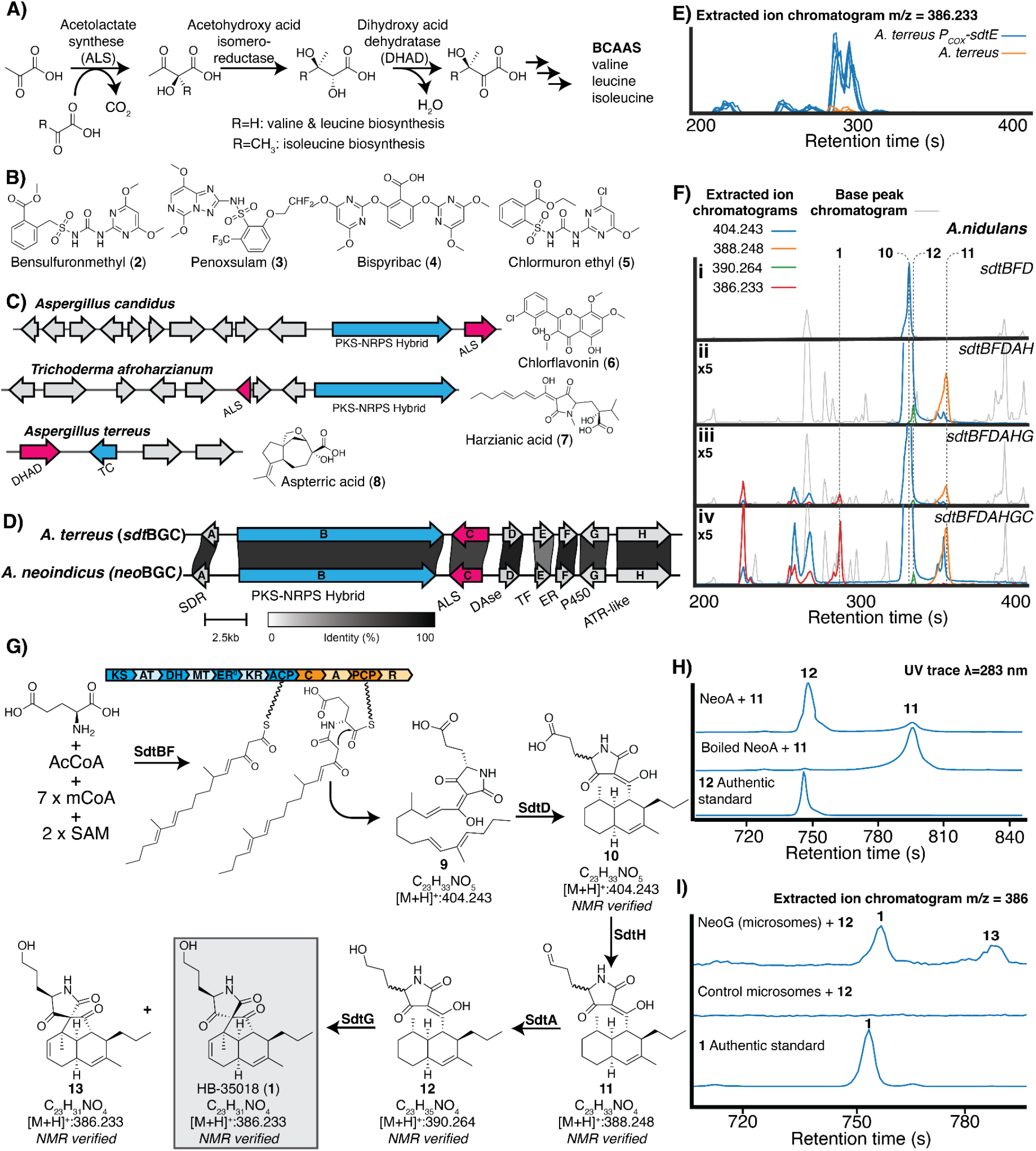
Discovery and biosynthetic characterization of 1. **A)** Overview of the biosynthesis of branched chain amino acids (BCAAs) **B)** Structure of select known inhibitors of ALS.**C)** BGCs that produce BCAA inhibitors **6**-**8** containing putative resistance genes homologous to genes involved in BCAA biosynthesis. **D)** *sdt*BGC from the genome of *Aspergillus terreus* and a related cluster from the genome of *Aspergillus neoindicus*. All proteins share >60% amino acid identity. Abbreviations: short chain reductase reductase (SDR), polyketide synthase/non-ribosomal synthetase hybrid (PKS/NRPS hybrid), Diels-Alderase (DAse), transcription factor (TF), enoyl-reductase (ER), cytochrome P450 (P450), adenylation-thiolation-reductase-like (ATR-like) **E)** Extracted ion chromatograms for *m/z* = 386.233 in wild-type *A. terreus* and *A. terreus* overexpressing the transcription for *sdtE* (*A. terreus* P_COX_-*sdtE*). n=4 clones per strain. **F)** Base peak chromatograms and extracted ion chromatograms of cultures of *A. nidulans* strains expressing subsets of genes from the *sdt*BGC **G)** Proposed biosynthesis of **1**. Key products and intermediates shown. Domain abbreviations: ketosynthase (KS), acyltransferase (AT), dehydratase (DH), methyltransferase (MT), ketoreductase (KR), acyl carrier protein (ACP), condensation (C), adenylation (A), peptidyl carrier protein (PCP), reductase (R). See **Figure S1** for a more detailed scheme inclusive of all identified shunt and side products. **H**) Conversion of **11** to **12** by recombinant NeoA reductase. **I**) Conversion of **12** to **1** by microsomes from *S. cerevisiae* strains expressing cytochrome P450 NeoG.

In this study by leveraging resistance gene-guided genome mining we discover the *sdt*BGC, a BGC that encodes within it a resistance gene homologous to ALS. Using heterologous expression coupled with in situ BGC activation we identify HB-35018 (**1**), as the product of the *sdt*BGC and a novel ALS inhibitor. Through detailed biosynthetic characterization of this unique spiro-*cis*-decalin tetramic acid, we uncovered the activities of several notable enzymes. These include the first lipocalin-type Diels-Alderase mediating *cis*-decalin formation *via* a normal electron demand intramolecular Diels-Alder reaction and a cytochrome P450 responsible for establishing the highly strained spiro center. We further demonstrate that **1** is a strong inhibitor of recombinant ALS from multiple fungal species and employ structural analyses using intact protein mass spectrometry and cryo-electron microscopy to reveal **1** to be a covalent inhibitor of ALS, a binding mode distinct from previously known inhibitors. The activity observed in biochemical assays translates to whole fungal cells with **1** demonstrating potent antifungal activity against a panel of *A. fumigatus* isolates and several other fungal pathogens. Finally, using a mouse model of invasive aspergillosis, we validate ALS as essential to the pathogenicity of *A. fumigatus*. These findings both validate ALS as a promising antifungal target and highlight the power of genome mining to uncover innovative chemical scaffolds with potential to be developed as new antifungals.

## Results

### Genome mining and biosynthesis of 1

Resistance gene-guided genome mining has emerged as a powerful approach for identifying inhibitors of the essential fungal BCAA biosynthesis pathway. ALS performs the first dedicated step in this pathway and is the sole target to date with two distinct characterized BGC families—those of chlorflavonin (**6**)^21^ and harzianic acid (**7**)^22^—containing resistance genes encoding homologous proteins discovered in fungi. Additionally, aspterric acid (**8**) has been identified as an inhibitor of dihydroxy acid dehydratase (DHAD), the third protein in the BCAA pathway (**Figure 1C**)^23^. Guided by these precedents, we initiated our discovery of novel BCAA inhibitors by mining our database of annotated fungal genomes for BGCs containing homologs of *ILV2*, the gene for ALS in *Saccharomyces cerevisiae*. This search yielded the *sdt*BGC in *Aspergillus terreus*, along with a closely related BGC in *Aspergillus neoindicus* (*neo*BGC, **Figure 1D**).

To determine the product of *sdt*BGC, we used two complementary approaches: in situ transcription factor activation and heterologous expression with complete refactoring in *A. nidulans*. In the first approach, *A. terreus* was transformed with a plasmid encoding *sdtE*, the gene for a transcription factor encoded within the BGC, under a constitutive promoter (P_COX_). Overexpression of such BGC-specific transcription factors has been shown to lead to significant upregulation of BGC expression and increased metabolite production^24^. Consistent with this precedent, LCMS analysis of *sdtE* overexpression strains led to the identification of multiple features that were upregulated as compared to the wild-type strain, including a prominent feature with *m/z* = 386.233 (**Figure 1E**). In parallel, the *sdt*BGC was refactored for heterologous expression in *A. nidulans*, resulting in the observation of multiple differential features, including the *m/z* = 386.233 feature identified through in situ BGC activation (**Figure 1Fiv**). Upon isolation, the metabolite responsible for this feature was determined to be HB-35018 (**1, Table S5, Figures S11**-**S16**), a unique spiro-*cis*-decalin containing tetramic acid.

Given the unique structure of **1**, we sought to understand key steps in its biosynthesis using a combination of heterologous expression strains containing subsets of genes from *sdtBGC* and in vitro enzyme characterization. Coexpression of *sdtB* and *sdtF* in *A. nidulans* demonstrated that, similar to other fungal tetramic acids^25^, biosynthesis is **1** is initiated by the production of an octaketide-glutamic acid hybrid through the activity of polyketide synthase/non-ribosomal synthetase (PKS/NRPS) hybrid SdtB and in trans enoyl reductase (ER) SdtF. Release of this hybrid from SdtB via a Dieckmann cyclization catalysed by the terminal reductase domain yielded compound **9** (**Figure 1G, Figure S1Ci**). Addition of *sdtD*, a gene encoding a lipocalin-type Diels-Alderase (DAase), to *A. nidulans sdtBF* yielded *cis*-decalin **10** (**Table S6, Figures S17**-**S22**) as the dominant product (**Figure 1Fi, Figure S1Civ, Figure 1G**). The stereospecific formation of *cis*-decalins is not commonly observed as they are thermodynamically disfavored relative to the *trans-*isomer. Characterized examples of *cis*-decalin formation in fungi include catalysis by an S-adenosylmethionine (SAM)-dependent DAase in fischerin biosynthesis^26^ and a two-step mechanism involving an oxidation for electron-demand inversion in the biosynthesis of varicidins A and B^27^. To our knowledge, SdtD represents the first lipocalin-type DAase known to catalyze *cis*-decalin formation directly via a normal electron demand intramolecular Diels-Alder reaction. Intriguingly, CghA, the DAase from the BGC for *trans*-decalin Sch210971^28^ has been successfully engineered to favor formation of the *cis*-product through the installation of three mutations, one of which, A242S, is present in SdtD (**Figure S2**), perhaps explaining this inverted stereospecificity^29^.

Subsequent to formation of the *cis*-decalin, addition of putative ATR (adenylation-thiolation-reduction) gene *sdtH* and the *sdtA* short-chain dehydrogenase/reductase (SDR) gene led to production of aldehyde **11** (**Figure 1G, Figure 1Fii, Table S7, Figures S23-S28**) and alcohol **12** (**Figure 1G, Figure 1Fii, Table S8, Figures S29-S34**). That SdtA alone leads to conversion of **11** to **12** was confirmed by incubation of **11** with recombinant NeoA, a close homolog of SdtA from the *neo*BGC (**Figure 1H**). The transformations mediated by SdtH and SdtA were specific for decalin-containing substrates, with the addition of either *sdtH* or *sdtA* to strains lacking *sdtD* showing no impact on the metabolomic profile (**Figure S1ii**,**Figure S1iii**). The final key transformation, installation of the spiro center at C2’, was catalyzed by the cytochrome P450 (P450) SdtG. Addition of *sdtG* to the heterologous expression system produced a complex mixture, as it could catalyse spiro center formation from both the C7’ acid and C7’ alcohol regardless of C4’ stereochemistry, resulting in a variety of shunt products (**Figure S1vi, 14-19, Tables S10-S14, Figures S41-S70**). To confirm SdtG alone catalyzed spiro center formation, we incubated **12** with microsomes from *S. cerevisiae* expressing *neoG*, an *sdtG* homolog from the *neo*BGC (**Figure 1D**). This reaction yielded conversion of **12** to spiro compounds **1** and its C2’ diastereomer **13** (**Figure 1I, Table S9, Figures S35-S40**). X-ray crystallography confirmed the absolute stereochemistry of **1** (**Figure S3**).

While the presence of a resistance gene homologous to *ILV2* within the *sdt*BGC suggested that ALS is the target of **1**, we undertook a genetic and biochemical approach to confirm the target. Initially we performed a forward genetic screen in which a mutagenized library of *S. cerevisiae* was exposed to **1** at concentrations ranging from 5 µM to 80 µM. This treatment resulted in the development of resistant clones. We performed whole genome sequencing on 24 clones and 23 had mutations in *ILV2*, the *S. cerevisiae* gene for ALS (*Sc*ALS), suggesting that ALS is the target of **1**. Resistance-conferring mutations localized near the substrate entry tunnel of the ALS homodimer, proximal to the established binding sites of known ALS inhibitors **2** and **5** (**Figure 2A**). Mutations in Pro192 were present in 75% of resistant clones (**Figure 2A, Figure S4, Table S1**), the same residue whose mutation imparted resistance to the triazolopyrimidine-sulfonamides, a class of ALS inhibitors previously explored as antifungals^20^. To further confirm that ALS is the target of **1**, we evaluated its ability to inhibit ALS-mediated formation of acetolactate from pyruvate using recombinant *Sc*ALS. We observed potent inhibition, in line with known ALS inhibitors **2-5**. However, while **2-5** were significantly less potent against P192S mutant *Sc*ALS compared to wild-type *Sc*ALS, with potencies diminished by at least 100-fold in all cases, there was only a four-fold shift in potency for **1** (**Figure 2B**). **1** is also significantly more potent against *Sc*ALS than its diastereomer **13**, further supporting the assignment of **1** as the final product of the *sdt*BGC (**Figure S5**).

**Figure 2:**
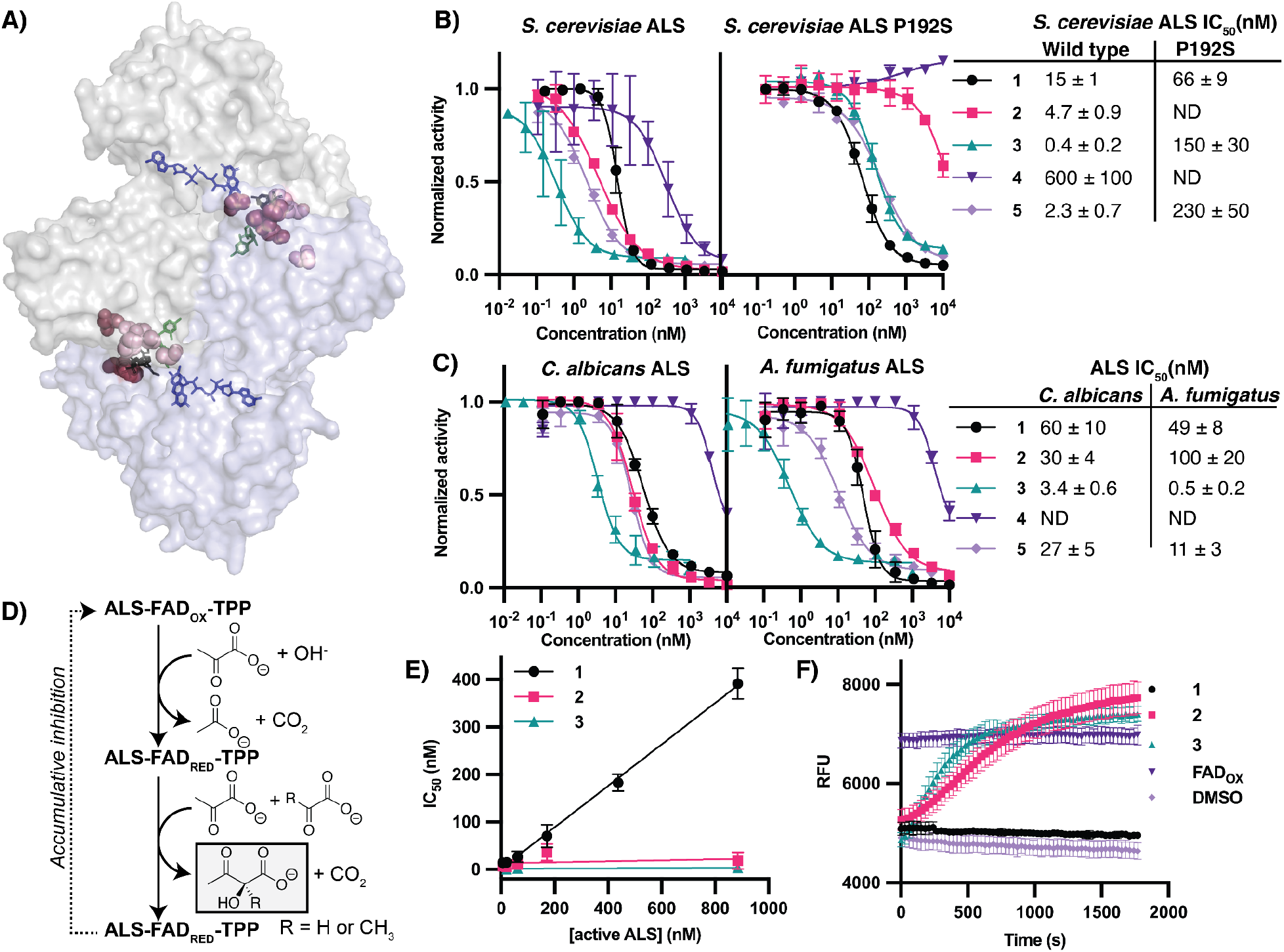
Confirmation that ALS is the target of 1. **A)** Mutations in ALS observed in *S. cerevisiae* clones resistant to **1**. Resistance mutations (shown in pink) are mapped onto the crystal structure of **2** (black) bound to *S. cerevisiae* ALS (PDB:5FEM^30^,α-chain in gray, β-chain in blue). Essential thiamine pyrophosphate (TPP,green) and FAD (blue) cofactors shown. **B)** Inhibition of wildtype (WT) and P192S mutants of ALS from *S. cerevisiae* by **1**-**3. C)** Biochemical inhibition by **1**-**5** of ALS from *S. cerevisiae, Candida albicans* and *A. fumigatus*. **D)** Reaction catalyzed by ALS with the oxidation state of FAD at each step indicated (FAD_OX_ = Oxidized, FAD_RED_ = reduced). **E)** IC_50_ vs concentration of active *S. cerevisiae* ALS protein suggests tight-binding kinetics for the binding of **1** and not for **2** or **3 F)** FAD cofactor reoxidation upon incubation of *S. cerevisiae* ALS with **1-3**. Oxidized FAD (FAD_OX_) included as a positive control.

To assess **1** as an antifungal, recombinant ALS from the key pathogenic fungal species *A. fumigatus* (*Af*ALS) and *C. albicans* (*Ca*ALS) was prepared. In both species, **1** demonstrated activity comparable to that observed against *Sc*ALS and of known ALS inhibitors **2**-**5** (**Figure 2C**).

Multiple ALS inhibitors, including both **2** and **3**, have been shown to function through accumulative inhibition, a mechanism by which inhibitors function not simply by occluding the substrate binding tunnel, but by trapping reactive oxygen species in the active site, leading to sustained inactivation of the enzyme through oxidation of the flavin adenine dinucleotide (FAD) cofactor. FAD in its reduced form is essential to ALS function, so this cofactor oxidation allows the effect of accumulative inhibitors to persist after dissociation and imparts super-stoichiometric activity (**Figure 2D**).^30,31^ Kinetic characterization indicates **1** does not function via this mechanism as standard tight-binding kinetics with IC_50_ equal to half the concentration of active ALS is observed (**Figure 2E**). Furthermore, we see no indication of FAD reoxidation upon treatment with **1** in contrast to known accumulative inhibitors **2** and **3** (**Figure 2F**).

We next characterized the structural basis for binding of **1** to ALS. Given the strained structure and the observed tight binding kinetics, we hypothesized that **1** may be binding covalently. Analysis of ALS from both *Af*ALS and *Sc*ALS by intact protein mass spectrometry yielded traces with multiple proteoforms for both proteins. Similar analysis after incubation of both proteins with saturating concentrations of **1** (10 μM) for 30 minutes produced mass shifts consistent with covalent addition of **1** (**Figure 3A**) supporting this hypothesis. Limited proteolysis followed by peptide mapping localized the covalent attachment to the _239_SGRPGPVLVDLPKDVTAAIL_258_ peptide of *Sc*ALS with Lys251 and Asp252 (equivalent to Lys310 and Asp311 in *Af*ALS) serving as the most likely attachment sites (**Figure S6**). The ability to detect modified peptide fragments following proteolysis suggests that **1** had irreversibly modified the ALS protein.

**Figure 3:**
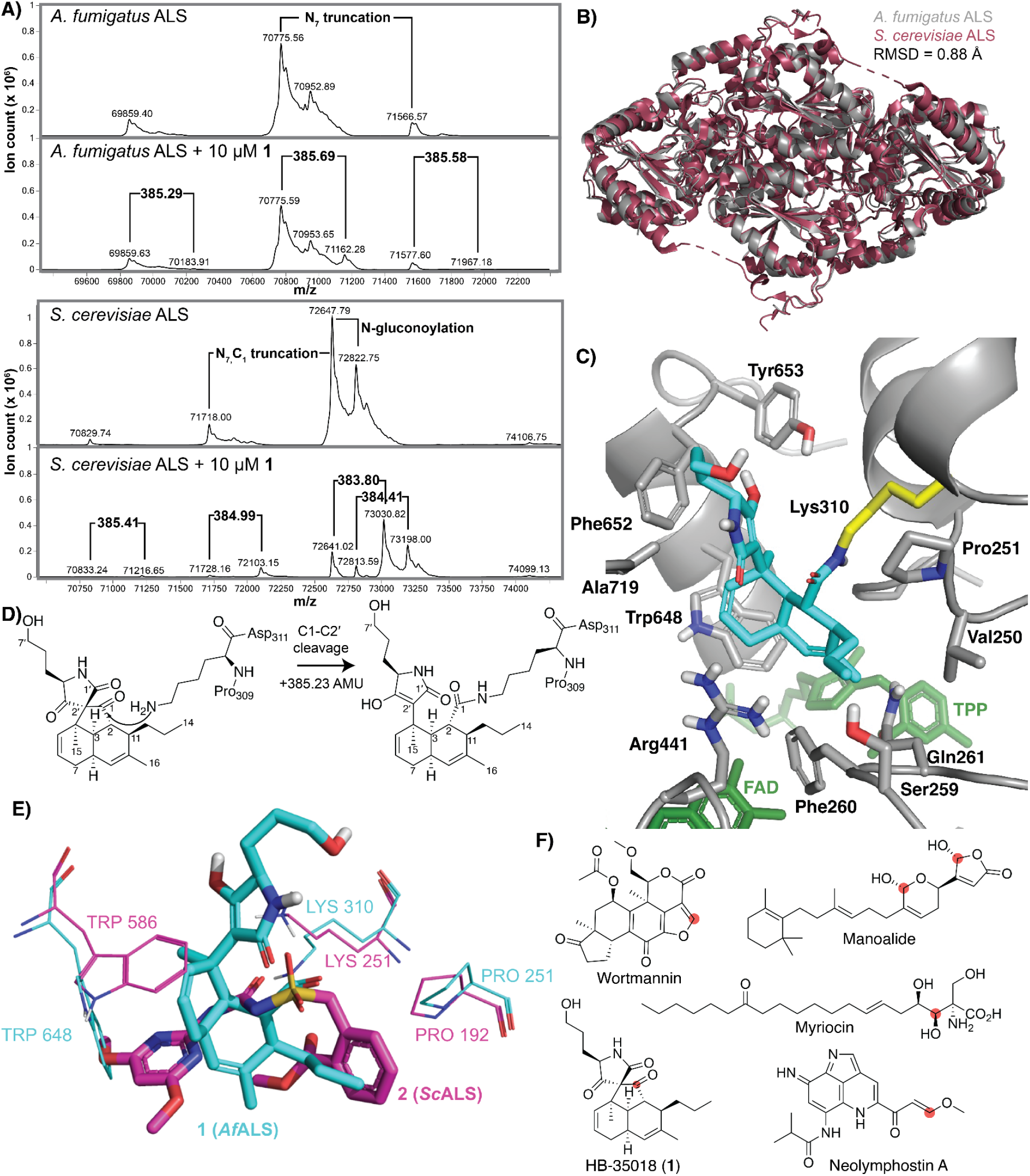
1 is a covalent inhibitor of ALS. **A)** Intact mass-spectrometry traces of *Af*ALS and *Sc*ALS both alone and treated with 10 μM **1**. Observed proteoforms are annotated in the untreated trace and *m/z* shifts in treated samples are highlighted. **B)** Cryo-EM structure of apo-*Af*ALS overlaid with *Sc*ALS (PDB 5FEM). **C)** Cryo-EM structure of *A. fumigatus* ALS with **1** (aqua) covalently bound to Lys310 (yellow) blocking the substrate channel (TPP is the cofactor thiamine pyrophosphate and FAD is the cofactor flavin adenine dinucleotide). **D)** Model for covalent addition of the sidechain of Lys_310_ into the C1 ketone of **1** with subsequent C1-C2’ bond fragmentation. **E)** Overlay of **1** (aqua) bound to *A. fumigatus* ALS with sulfonylurea herbicide bensulfuronmethyl **2** (pink) bound to *S. cerevisiae* ALS (PDB 5FEM) ^30^. **F)** Examples of natural products that covalently modify lysine residues of target proteins. Electrophilic attachment points are highlighted in red.

Structures of multiple plant and bacterial ALS proteins exist, but fungal structures are limited to those of *S. cerevisiae* and *C. albicans*^*32,33*^. To more fully understand the interaction of **1** with ALS from a clinically relevant fungus, as well as facilitate optimization of this scaffold and development of future antifungals, we determined the structure of *Af*ALS. Using cryo-electron microscopy, we solved the structures of both the apo-ALS homodimer at a resolution of 2.76 Å and the **1**-bound form with a resolution of 2.36 Å (**Figures 3B and 3C, Figure S10, Table S2**). Electron density related to compound **1** was observed in the narrow substrate channel at the homodimer interface, which is consistent with the established binding site for multiple herbicide ALS inhibitors, including **3**-**5**. In agreement with our proteomic data, the electron density was continuous with the side chain of Lys310, consistent with covalent modification of the protein and inconsistent with the spiro center of **1** being intact (as would be expected for non-covalent binding) (**Figure 3C, Figure S7A**). The structure achieving the best fit to the observed density contained an amide between C1 and the ε-amine of Lys310 with fragmentation of the C1-C2’ bond. This could arise through the reversible reaction of Lys310 with the C1 ketone, followed by irreversible fragmentation of the C1-C2’ bond in a retro-Dieckmann-like reaction (**Figure 3D**).

The proposed fragmentation is similar to the amine-induced fragmentation of tricarbonyl C-acyl β-ketoesters first described by C. Kitsiou et al.^34^ The driving force for this fragmentation is likely the relief of ring strain in the spiro center. In the covalently-bound, post-fragmentation state, the decalin portion of the ligand is within 4.5 Å of hydrophobic amino acid sidechains of Trp648, Phe260, Pro251, and Val250, as well as the π-faces of polar sidechains of Arg441 and Gln261, and the oxygen of the Ser259 sidechain. The tetramic acid portion of the ligand, modeled for convenience as the enol tautomer, is within 4.5 Å of Tyr653, Phe652, and Ala719 (**Figure 3C**). Compared to the structure of apo *Af*ALS, binding of **1** triggered major conformational changes in the side chains of Lys310, Arg441, Trp648, and Phe652 (**Figure S7B**). The side chain of Trp649 changed orientation due to spatial clash with the decalin ring of **1**, the aromatic ring of Phe652 shifted closer to interact with **1** via π-stacking interactions, while the side chain of Arg441 also changed orientation to interact with **1** through electronic interactions. Due to the proximity of the Tyr653 oxygen and the C3’ oxygen (O-O distance 4.1 Å), it is possible that the OH of Tyr653 assists in C1-C2’ fragmentation by providing a hydrogen-bond to the C3’ ketone, activating it to accept the electrons from the C1-C2’ bond. The ligand-protein interactions of inhibitor **1** and *Af*ALS are largely consistent with the *S. cerevisiae* forward genetics experiments, with 5/6 mutated residues observed within 5 Å of ligand atoms in the *Af*ALS structure (**Supporting Table S1**). Comparison of the binding of **1** and **2** to *Af*ALS provides a potential justification for the differential resistance to P192 mutants as P192 (P251 in *Af*ALS) contacts a flexible propyl group in **1**, likely resulting in a weaker interaction, while P192 contacts a much more rigid benzyl group in **2**, likely resulting in a stronger interaction.

While **1** binds the same region as synthetic herbicides, the binding mode is distinct as shown in the overlay in **Figure 3E**. In the **1**-AfALS (aqua) and **2**-ScALS (pink) structures the sidechains of Trp648 (*Af* numbering) and Trp586 (*Sc* numbering) are observed in different conformations and there is little overlap between the atoms of **1** and **2**.

The side-chain thiol of cysteine is by far the most frequent point of attachment to proteins by covalent inhibitors, but lysine represents an attractive target, partially due to lysine’s significantly higher abundance in the proteome (5.8% lysine vs. 1.9% cysteine).^35^ Despite this promise, design of lysine targeting covalent inhibitors is often challenging due to the high pKa of the Lys ε-amino group.^36^ Several natural products are among those compounds known to covalently modify lysine, including manoalide, which irreversibly inhibits phospholipase A2,^37^ wortmannin and neolymphostin A, both of which target phosphoinositide 3-kinases (PI3K),^38,39^ and myriocin, which covalently binds to serine palmitoyltransferase (**Figure 3F**).^40^ The mechanism of **1** is unique among these examples with strain release via C-C bond fragmentation serving to irreversibly trap a lysine sidechain in an amide linkage.

With the mechanism of ALS inhibition by **1** established and understood, we next tested the translation of this biochemical potency to antifungal activity. While the potency of **1** was approximately equivalent to that of **2**-**5** in biochemical screens, **1** inhibited *A. fumigatus* growth similarly to voriconazole (IC_50_=0.7 µM and 0.5 µM, respectively) and was more potent than the all the known ALS inhibitors (IC_50_= 200 µM (**2**), 25 µM (**3**), ND (**4**), 90 µM (**5**)). I (**Figure 4A**). The antifungal activity of **1** extends beyond lab strains and persists in clinically relevant isolates of *A. fumigatus* from the antimicrobial resistance cell bank from the Center for disease control (CDC)^41^ including multiple isolates resistant to azoles, the standard of care for aspergillosis^42^ (**Figure 4B**). **1** also inhibits growth of a collection of rarer pathogenic fungi including *Aspergillus niger, Rhizopus oryzae, Mucor circinelloides, Scedosporium apiospermum*, and *Scedosporium prolificans* (**Figure 4B**).

**Figure 4:**
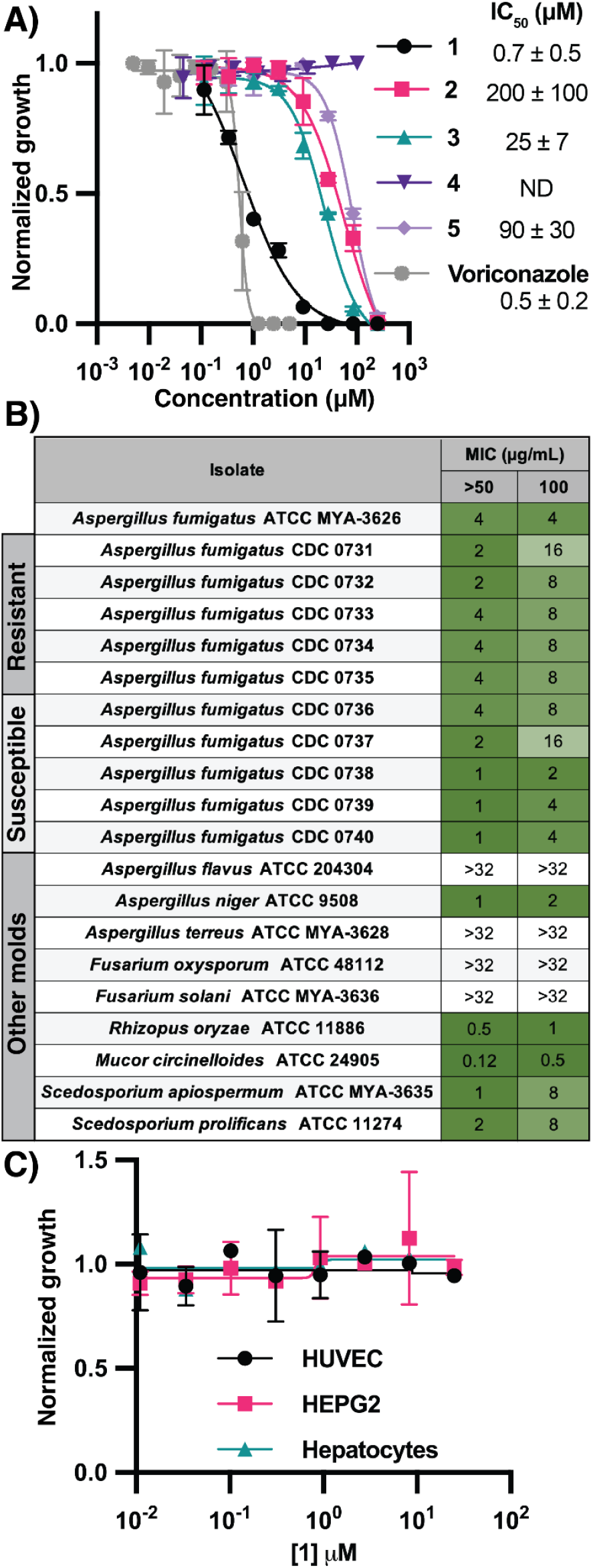
1 is a potent inhibitor of the pathogenic fungi. **A)** Inhibition of *A. fumigatus* growth by ALS inhibitors **1**-**5. B)** Minimum inhibitory concentrations (MICs) for >50 % and 100% growth inhibition of a panel of fungi by **1** at 24 hrs. Isolates of *A. fumigatus* are labeled as either ‘resistant’ depending on azole susceptibility per the CDC ARIsolate bank. **C)** Cytotoxicity of **1** against primary human cells HUVEC (human umbilical vein endothelial cells) and hepatocytes and the liver cancer cell line HEPG2..

Consistent with its observed antifungal activity arising from specific inhibition of ALS, a target with no close homolog in humans, **1** shows no cytotoxicity when tested against a panel of primary human cells and a cell line at concentrations up to 100 μM, suggesting that off-target toxicity is unlikely to be a liability (**Figure 4C**).

Essential amino acids, including BCAAs, are so named because humans must acquire them through their diet. This is in contrast to fungi and bacteria, which can synthesize these nutrients *de novo*. This distinction makes proteins involved in BCAA biosynthesis attractive targets for antifungal development. However, a potential limitation of targeting these pathways is the ability of pathogenic organisms to scavenge BCAAs from their environment, rendering these targets conditionally essential. Despite this concern, ALS has been shown to be essential for the pathogenicity of both *Cryptococcus neoformans*^*43*^ and *Candida albicans*^*44*^ in mouse models of infection. These findings suggest that host-derived BCAA scavenging is insufficient to compensate for ALS loss from these species in the context of an infection. We aimed to determine whether the same holds true for invasive aspergillosis caused by *A. fumigatus*.

To investigate this, an *A. fumigatus* AF293 strain lacking the *ilv2* gene was constructed. In vitro growth assays revealed that this *ilv2Δ* strain required exceptionally high levels of BCAAs in the medium to proliferate (**Figure S8A-C**), indicating limited BCAA import capacity. In vivo infection studies in a mouse model of invasive aspergillosis showed a significant increase in the survival of mice infected with an *ilv2Δ* strain (**Figure 5A**), demonstrating reduced virulence.

**Figure 5:**
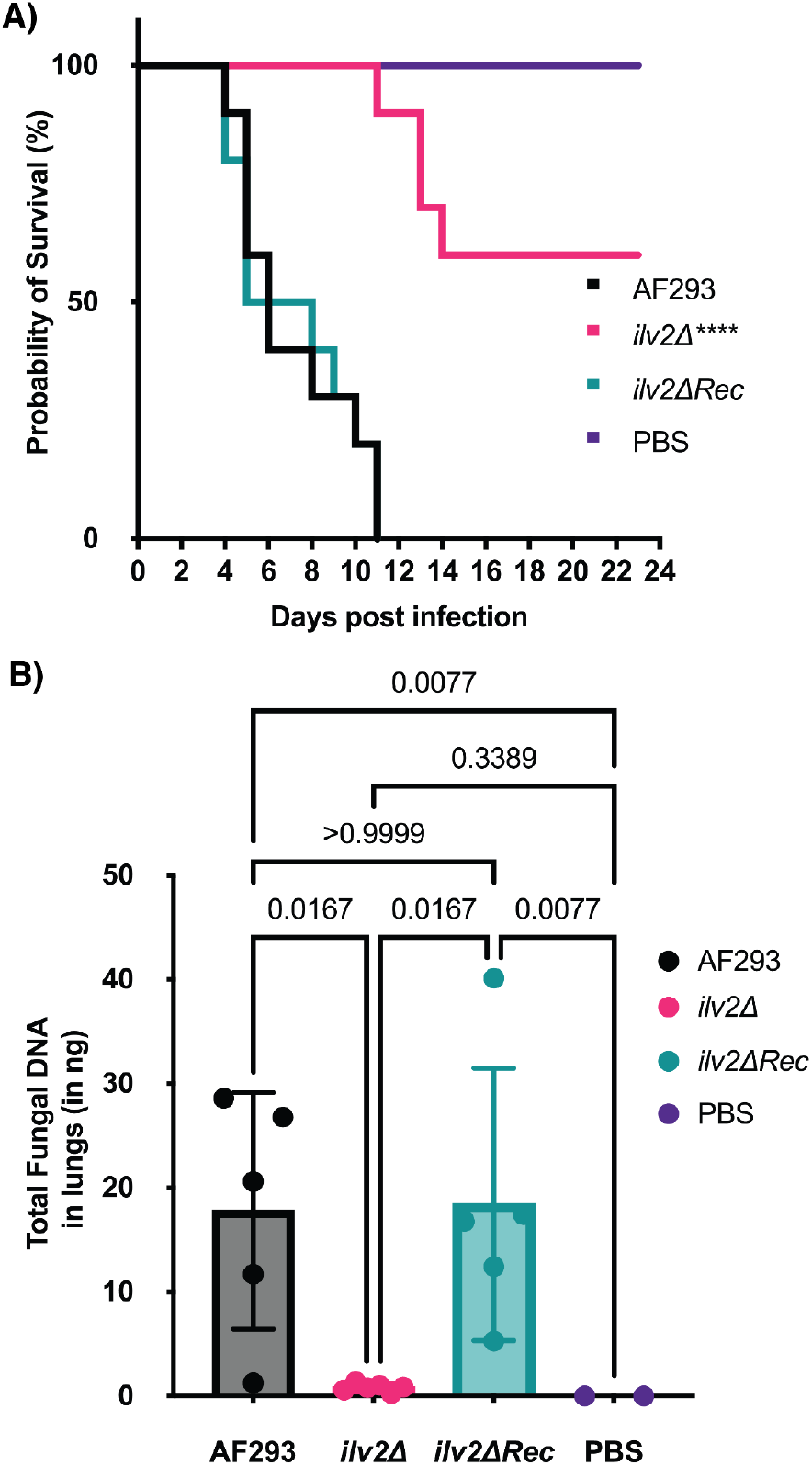
ALS is essential for pathogenicity in invasive aspergillosis. **A)** Survival curves for a triamcinolone mouse model of invasive aspergillosis using *Aspergillus fumigatus* AF293 (wild-type), a mutant of AF293 with *ilv2* (the gene for ALS) deleted (*ilv2Δ*), and the deletion mutant with *ilv2* expression reconstituted at *aft4* safe haven site (*ilv2ΔRec*). Log-rank (Mantel-Cox) test was significant (****P<0.0001; n=10 for infected groups; PBS n=6). **B)** Total fungal DNA quantified in the lungs of mice in each group. Kruskal-Wallis non-parametric test was significant (****P=0.0002; AF293/*ilv2*ΔRec: n=5; *ilv2*Δ n= 6; PBS n=2). Dunn’s post-hoc test was used for pair-wise comparisons. The p-values are indicated on the graph.

Furthermore, restoration of *ilv2* expression in the mutant background fully recapitulated the pathogenicity of the wild-type strain (*ilv2ΔRec*, **Figure 5A**). The inability of the *ilv2Δ* strain to sufficiently grow in the respiratory tract at day 3 post-infection was supported by a 20-fold reduction in fungal DNA levels in the lungs of infected mice, suggesting significantly reduced fungal burden as compared to those infected with wild-type AF293 or *ilv2ΔRec* strains (**Figure 5B**).

These results demonstrate that ALS is essential for the pathogenicity of *A. fumigatus* in the context of invasive aspergillosis and establish it as a promising therapeutic target for antifungal treatment. Next we assessed the potential of **1** for in vivo efficacy through detailed characterization of its physicochemical and pharmacokinetic properties (**Table S3**). We found that **1** is unstable in multiple matrices, including pH 7.4 PBS buffer (t_1/2_ = 141 min), mouse plasma (t_1/2_ = 88 min), and mouse liver microsomes (t_1/2_ = 3 min). It is likely that the instability in these matrices is due to fragmentation chemistry similar to that described between **1** and ALS (**Figure 3D**), with water (or another biorelevant nucleophile) reacting with the C1 ketone followed by fragmentation of the C1-C2’ bond. From these data, we reasoned that even though **1** is a potent, irreversible covalent ALS inhibitor, it would be unlikely to achieve the plasma concentrations needed for 100% inhibition of pathogenic *A. fumigatus* strains at feasible dose levels. Therefore, we believe **1** will require further optimization to achieve success in in vivo efficacy studies.

## Discussion

The rising global incidence of invasive fungal infections, now responsible for over 2 million deaths annually, underscores an urgent need for new antifungal treatments. Current therapies face increasing resistance and limited efficacy, driving a critical need for novel antifungal agents with new mechanisms of action. Here, we utilize resistance gene-guided genome mining to discover such an agent in HB-35018 (**1**), an inhibitor of BCAA biosynthesis with potent in vitro activity across a breadth of pathogenic fungi. This antifungal activity is in stark contrast to what we observe from known ALS inhibitors which each show potent biochemical activity, but are very poor antifungals. It is possible that, for these herbicides, their anionic nature at cellular pH is a liability. In fungi, ALS is localized to the mitochondria, meaning that ALS inhibitors must cross the cell wall, cell membrane, and mitochondrial membrane to exert activity.^45^ The passive permeability of anionic compounds is poor and they are likely to become substrates for the organic anion transporter (OAT) family of efflux pumps, providing a possible explanation for the lack of antifungal activity among most known ALS inhibitors.^46^

Intriguingly, the *sdt*BGC, encoding **1**, represents the third BGC—along with those of chlorflavonin (**6**) and harzianic acid (**7**)—to contain an ALS-homologous putative resistance gene that produces a verified ALS inhibitor. Furthermore, the *sdt*BGC was identified in the genome of *A. terreus*, which also harbors the BGC for aspterric acid (**8**). The latter contains a homolog DHAD, the molecular target of **8** and a key enzyme downstream of ALS in the BCAA pathway. The evolution of three distinct BGC families with ALS-homologous resistance genes, coupled with the occurrence of two BGCs with resistance genes from the same pathway in a single fungal genome, demonstrates a convergence on BCAA biosynthesis as a significant target for antifungal activity.

While this convergence on BCAA biosynthesis suggests the promise of this pathway as an antifungal target in the ecologies in which these fungi evolved, here we demonstrate that this promise translates into a therapeutic context by showing that ALS is essential to pathogenicity in a mouse model of invasive aspergillosis. Whether or not this essentiality translates to biosynthesis of other essential amino acids remains to be seen, but were this to be the case, it would unlock another large set of promising antifungal targets.

Finally, in establishing that **1** acts through a covalent modification of a lysine residue, we report the cryo-EM structure of the ALS of *A. fumigatus*. While the inherent instability of **1** may limit its direct use in in vivo studies, this structure provides a clear path forward for optimizing the scaffold into a viable therapeutic. The detailed structural insights will not only enable the rational design of more stable analogs but will also facilitate future efforts toward the discovery and development of novel antifungals targeting this protein. While this manuscript was in the final stages of preparation, another study was published describing the biosynthesis of **1** and demonstrating that it has modest herbicidal activity, demonstrating another path forward for the development of **1**.^47^

Overall, this study highlights the power of genome mining as an effective strategy for the discovery of unique natural product scaffolds with therapeutic potential and by demonstrating the repeated evolutionary targeting of BCAA biosynthesis by distinct fungal BGCs, reinforces the relevance of this pathway as a target in further antifungal drug development efforts.

## Support information

Supporting Methods, Supporting Tables S1-S14, and Supporting Figures S1-S70 are available in the Supporting Information.

## Data Availability

The small molecule X-ray crystal structure of **1** is deposited in the Cambridge Crystallographic Data Centre with deposition number CCDC 2491792

The model of the apo and **1**-bound *Af*ALS have been deposited in the wwPDB with accession code PDB 9YJZ and 9YK0 respectively. The cryo-EM maps of the apo and **1**-bound *Af*ALS have been deposited in EMDB with accession codes EMD-73040 and EMD-73041.

## Acknowledgements

We would like to thank Dr. Tsokyi Choera, Hong Shi, Yi Zhou, and Dr. Chiraj Dalal for their contributions to this work.

## Notes

### Competing Interest Statement

BP, YC, KD, AF, MFG, SL, TO, RS, TA, YT, VC, and CJBH are all shareholders in Hexagon Bio.

