## Supporting Information for "An antifungal with a novel mechanism of action discovered via resistance gene-guided genome mining"

### Materials and Methods

##### Media and buffer composition

ST buffer: 1.2 M sorbitol, 12.5 mM Tris-HCl, pH 7.5

STC buffer: 1.2 M sorbitol, 10 mM calcium chloride, 12.5 mM Tris-HCl, pH 7.5

Digestion buffer: 280 g/l $MgSO_4 \cdot 7H_2O$, 120 g/l Vino Taste Pro, in a 175 mM sodium phosphate buffer containing 14.79 g/l $NaH_2PO_4 \cdot H_2O$, 17 g/l $Na_2PO_4$, with pH adjusted to 7.0

GMM: 10 g/L glucose, 6 g/L $NaNO_3$, 0.52 g/L KCl, 0.52 g/L $MgSO_4 \cdot 7H_2O$, 1.52 g/L $KH_2PO_4$, 1 mL/L trace elements concentrate, 0.05 w/v Riboflavin

YG: 20 g/l glucose, 10 g/l yeast extract, and 1 mL trace elements concentrate

20X salt solution: 120 g/l $NaNO_3$, 10.4 g/l KCl, 10.4 g/l $MgSO_4 \bullet 7H_2O$, 30.4 g/l $KH_2PO_4$

SMK: 40 g/L soluble starch, 1 g/L yeast extract, 4.3 g/L Murashige and Skoog Basal Salt Mixture

Trace elements concentrate: 1.0 g/L $FeSO_4.7H_2O$, 1.0 g/L $MnSO_4.H_2O$, 0.2 g/L $ZnSO_4.7H_2O$, 0.1 g/L $CaCl_2.2H_2O$, 0.056 g/L $HBO_3$, 0.025 g/L $CuCl_2.2H_2O$, 0.019 g/L $(NH_4)6.Mo_7O_{24}.4H_2O$, 50 mL/L 12M HCl)

YPD medium: 20 g/L peptone, 10 g/L yeast extract, 10 g/L dextrose

TEGM buffer: 50 mM Tris-HCl, pH 7.5, 1 mM EDTA, 0.6 M sorbitol, 1.5 mM β-mercaptoethanol, 20% glycerol (v/v), Pierce$^{TM}$ protease inhibitor

Formulation buffer: 300 mM $K_2HPO_4$, pH 8, 10 µM FAD, 0.5 mM TCEP, 10% glycerol v/v

##### Heterologous production of 1 in *Aspergillus nidulans*

To produce the metabolites encoded in the *sdt*BGC, we refactored the BGC in *A. nidulans* using the asperfuraone(*afo*) regulon with the gene for the positive regulator AN1029 (*afoA*) expressed under the inducible alcA promoter ($P_{alcA}$) as previously described.[1] In brief, intergenic regions of the *afo* regulon were amplified by PCR from the gDNA of *A. nidulans* and 7 genes of the *sdt*BGC were ordered as synthetic DNA from Twist Bioscience and PCR amplified. All relevant amplicons were gel purified and put through intermediate assemblies using NEBuilder HiFi DNA Assembly Master Mix (NEB, #E2621). The assembled DNA fragments were then used as templates for PCR to generate seven large amplicons, ranging in size from 4.2 to 8.5 kb for transformation. The recipient strain was derived from *A. nidulans* A1145 (the Fungal Genetics Stock Center strain number, https://www.fgsc.net/), in which the BGCs for sterigmatocystin (AN7804 – AN7825) and asperfuranone (AN1036 – AN1028) have been deleted. Protoplast preparation and *A. nidulans* transformation of the 7 prepared fragments were carried out as previously described.[2] After transformation, prototrophic colonies were randomly picked and examined by diagnostic PCR.

For each transformation, 6 individual clones were picked and inoculated into each of two deep-well blocks, one with 750 µl of glucose minimal media (GMM) in each well and the second with 750 µl lactose containing minimal media (LMM) per well. All cultures were grown for 48 hours at 30°C with shaking at 750 RPM prior to induction by the addition of 3.75 µl (0.5 % v/v) methylethyl ketone. After a further 72 hrs of growth under the same conditions, all plates were frozen and lyophilized overnight. Lyophilized culture solids were resuspended in 750 µl of methanol, left to soak for 2 hours and filtered through 2 µM filter plates prior to LCMS analysis.

##### Overexpression of the *sdt*BGC in *Aspergillus terreus*

The cluster specific transcription factor gene *sdtE* was cloned under the control of the $P_{COX}$ promoter from the *A. nidulans* cytochrome C oxidase and the terminator from gene AN0717 on a plasmid containing an AMA1 origin of replication and a hygromycin resistance marker. 2 µg of the assembled plasmid was used for each transformation.

To prepare biomass for transformation, Aspergillus pseudoterreus (NRRL4017, isolated from a soil sample taken in Argentina) was cultured for 24 hr at 180 rpm in 100 mL of YG media. Mycelia were collected from this culture and prepared for transformation as described previously[2]. Briefly, mycelia were incubated at 30°C and 100 rpm for 4-5 hr in 100 mL of freshly prepared and filtered digestion buffer. 45 mL of this protoplast suspension was filtered through miracloth, and 5 mL of 0.4 M ST buffer was gently layered on top. Samples were centrifuged at 1800 G for 10 min with deceleration on the gentlest setting. Protoplasts were collected at the interface and washed once with 0.6M KCl solution containing 44.7 g/l KCl, and once with 0.6M KCl, $CaCl_2$ solution containing (g/L): KCl, 44.7; $CaCl_2$ 7.4. Protoplasts were quantified and diluted to $1.0 \times 10^8$ protoplasts/ mL with KCl, $CaCl_2$ solution. Transformation of the prepared protoplasts was undertaken with 2 µg of plasmid DNA as previously described.[2] Each transformation was added to 35 mL of molten minimal media + KCl + hygromycin containing: 15 g/l glucose, 44.7 g/l KCl, 8 g/l agar, 0.4 g/l hygromycin, 1 mL trace elements solution, and 50 mL of 20 X salt solution. This mixture was gently inverted and poured into a 10 cm petri dish. Transformations were incubated at 37°C for 3-5 days at which point colonies were picked, cultured as described above, and subjected to untargeted metabolomic analysis.

##### Untargeted metabolomics analysis

High-resolution LCMS analyses were performed on a Thermo Fisher Scientific Vanquish Horizon UHPLC System integrated with a Thermo Q Exactive hybrid quadrupole-orbitrap high-resolution mass spectrometer with a HESI ion source. Two µL of the filtered culture extract to be analysed was injected and separated using a water-acetonitrile gradient on an Accucore Vanquish C18+ Reversed Phase HPLC Column (1.5 µm, 2.1 mm x 50 mm, Cat# 27101-052130) maintained at 50°C. Solvent A: 0.1% formic acid in water; Solvent B: 0.1% formic acid in acetonitrile. A/B gradient started at 12.5% B increased linearly to 97% B at 8.5 min and held at 97% B for 1 min, using a flow rate 0.6 mL/min. Mass spectrometer parameters: spray voltage 3 kV for positive mode, capillary temperature 380 °C, probe heater temperature 400 °C; sheath, auxiliary, and spare gas 60, 20, and 1, respectively; S-lens RF level 50, resolution 140,000 at m/z 200, AGC target $3\times 10^{10}$. Tandem mass spectrum analysis was carried out with an exclusion list of known endogenous features with the same parameters (vide supra) with the following additions or adjustments: Full MS Resolution: 35,000, AGC target 1 x $10^6$, Max IT: 30 ms, scan range: 150 to 1400 m/z, ddMS2 Resolution: 17,500, AGC target 5 x $10^4$, Max IT: 50 ms, Loop Count: 10, isolation window: 2.0 m/z, stepped collision energy: 20, 40, 60 NCE.

**Expression and purification of NeoA from *E. coli***

The *neoA* gene was amplified with overhang primers from the cDNA of strain *A. nidulans* expressing *neoABDFGH*. The PCR product was cloned into pET-28a(+) digested with NdeI/BamHI using NEBuilder® HiFi DNA Assembly Master Mix (NEB, #E2621). The resulting plasmid was confirmed by DNA sequencing and introduced into *E. coli* BL21(DE3) cells for $His_6$-tagged protein induction and purification. The *E. coli* cells harboring the corresponding plasmid were grown overnight in LB medium with 50 μg/mL kanamycin at 37°C, 220 rpm. 5 mL of the starter culture was inoculated into 1 L of fresh LB medium with 50 μg/mL kanamycin and incubated at 37°C until the optical density at 600 nm ($OD_{600}$ value) reached 0.6. After the culture was submerged in ice water for 25 min, protein expression was induced by addition of 100 μM isopropyl-β-D-thiogalactopyranoside (IPTG) and cultured for an additional 20 hrs at 16°C, 220 rpm. Cells were subsequently harvested by centrifugation at 5,000 rpm, 4°C for 10 min, resuspended in 40 mL lysis buffer (50 mM Tris-HCl, pH 7.5, 10 mM imidazole, 500 mM NaCl and 10% glycerol, v/v), and lysed by sonication on ice. The lysate was centrifuged at 14,000 rpm, 4°C for 30 min to remove the cellular debris. Then the supernatant was subjected to Ni-NTA affinity chromatography at 4°C for protein purification according to the manufacturer's protocols (GE Healthcare). The purified protein was concentrated and exchanged into a storage buffer (50 mM Tris-HCl, 50 mM NaCl, pH 7.5) by using Amicon® Ultra-15 Centrifugal Filters. The purity of protein was checked by SDS-PAGE, and the concentration was determined by Bradford method. The purified proteins were flash-frozen in liquid $N_2$ and stored at -80°C.

**Expression and purification of microsomes containing NeoG from *S. cerevisiae***

*S. cerevisiae* RC01 transformed with a plasmid expressing *neoG* was cultivated in 5 mL selective dropout medium at 28°C for 20 h. 4 mL of the starter culture was inoculated into YPD medium for an additional 48 h at 28°C, 220 rpm. The cells were harvested by centrifugation at 4,300 rpm, 4°C for 5 min, and the cell pellet was resuspended in 4 mL TEGM buffer. Glass beads (0.5 mm in diameter) were added and the cell walls were disrupted by vortexing for 1 min separated by 1 min intervals on ice for a total of 12 min. Then the microsomal fractions were obtained as described previously.[3] The microsomal fractions were resuspended in 50 mM Tris-HCl (pH 7.5, 50 mM NaCl, 10% glycerol, v/v).

**In vitro characterization of the activities of NeoA and NeoG**

Enzymatic assays of NeoA were performed in 50 mM NaPB (pH 7.6) with a final volume of 100 μL containing 10 μM NeoA with 250 μM compound **11** and 10 mM NADPH. The reactions were incubated at 28°C overnight and then quenched with 300 μL $CH_3CN$. The mixtures were centrifuged at 17,000 g for 5 min. LCMS analysis of clarified supernatants was performed on a Shimadzu 2020 EV LC–MS with a reversed-phase column (Phenomenex Kinetex 1.7 μm C18 100 Å, LC Column 100 × 2.1 mm) using positive-and negative-mode electrospray ionization with a linear gradient of 5–95% $CH_3CN$-$H_2O$ with 0.1% FA in 15 min followed by 95% $CH_3CN$ for 3 min with a flow rate of 0.3 ml/min.

Assays of microsomes containing NeoG were performed in 50 mM NaPB (pH 7.6) with a final volume of 500 µL containing 50 µL microsomal fractions with 50 µM **12** and 500 µM NADPH. The reactions were incubated at 28°C for 13 h and extracted with 550 µL EtOAc. The EtOAc extracts were evaporated to dryness and then redissolved in methanol for LC-MS analysis. The samples were analyzed by an Agilent 1260 Infinity II LC equipped with an InfinityLab Poroshell 120 EC-C18 column (2.7 µm, 3.0 × 50 mm) and a 6545 QTOF high resolution mass spectrometer using the following gradient: 1% $CH_3CN-H_2O$ 2 min, then 1−95% $CH_3CN-H_2O$ (both with 0.1% formic acid (FA), v/v) in 9 min followed by 95% $CH_3CN-H_2O$ for 6 min at a flow rate of 0.8 mL/min.

**ALS biochemical assay (*Sc*ALS)**

*Saccharomyces cerevisiae* acetohydroxyacid synthase (*Sc*ALS) lacking the N-terminal mitochondrial-targeting sequence (amino acids 58-687) was overproduced and isolated from *Escherichia coli* as an N-terminally $His_6$-tagged protein. Assays (50 µL final volume) contained 100 nM $His_6$-*Sc*ILV2 (58-687), 8 mM pyruvate, 1 mM thiamine pyrophosphate, and 10 µM flavin adenine dinucleotide, 10 mM MgCl2, 1 mM DTT, 1% DMSO (± inhibitor) in 500 mM $K_2HPO_4$ buffer at pH 7.5. Following a 30 min pre-incubation of the enzyme and inhibitor in the reaction mixture at room temperature, reactions were initiated with the addition of pyruvate. *Sc*ALS assays were carried out at 30˚C for 60 min. The reaction was quenched with the addition of 5 µL of 10% $H_2SO_4$ (v/v in deionized water). Reactions were incubated at 60˚C at 800 rpm in an incubating microplate shaker for 15 min to convert the product of the ALS reaction, acetolactate, to acetoin. Subsequently, the concentration of acetoin was determined by the addition of 7.5 µL 6 M NaOH, 25 µL 0.5% creatine (w/v in deionized water), and 25 µL 4% 1-naphthol (w/v in 2.5 M NaOH). This mixture was incubated at 60˚C at 800 rpm in an incubating microplate shaker for 15 min. 40 µL of this reaction mixture was transferred to a half area UV-star 96-well plate (Grenier) and the absorbance at 525 nm was measured on a Spark multimode plate reader (Tecan). The percent inhibition was calculated by the comparison of the signal to DMSO (0% inhibition) and no enzyme (100% inhibition) controls. $IC_{50}$ values were calculated from the dose response curves using Prism 9 (four parameter, variable slope).

**ALS biochemical assay (*Af*ALS)**

*Aspergillus fumigatus* acetohydroxyacid synthase (*Af*ILV2) lacking the N-terminal mitochondrial-targeting sequence (amino acids 112-752) was overproduced and isolated from *E. coli* as an N-terminally $His_6$-tagged protein. Assays (50 µL final volume) contained 200 nM $His_6$-*Af*ALS, 8 mM pyruvate, 1 mM thiamine pyrophosphate, and 10 µM flavin adenine dinucleotide, 10 mM MgCl2, 1 mM DTT, 1% DMSO (± inhibitor) in 200 mM KH2PO4 buffer at pH 7.5. Following a 30 min pre-incubation of the enzyme and inhibitor in the reaction mixture at room temperature, reactions were initiated with the addition of pyruvate. *Af*ALS assays were carried out at 30˚C for 120 min. The reaction was quenched with the addition of 5 µL of 10% $H_2SO_4$ (v/v in deionized water). Reactions were incubated at 60˚C at incubating microplate shaker for 15 min to convert the product of the ALS reaction, acetolactate, to acetoin. Subsequently, the concentration of acetoin was determined by the addition of 7.5 µL 6 M NaOH, 25 µL 0.5% creatine (w/v in deionized water), and 25 µL 4% 1-naphthol (w/v in 2.5 M NaOH).

This mixture was incubated at 60˚C in an incubating microplate shaker for 15 min. 40 µL of this reaction mixture was transferred to a half area UV-star 96-well plate (Grenier) and the absorbance at 525 nm was measured on a Spark multimode plate reader (Tecan). The percent inhibition was calculated by the comparison of the signal to DMSO (0% inhibition) and no enzyme (100% inhibition) controls. $IC_{50}$ values were calculated from the dose response curves using Prism 9 (four parameter, variable slope).

**Expression and purification of *Sc*ALS**

*S. cerevisiae* ILV2 (the gene or *Sc*ALS) lacking the N-terminal mitochondrial-localization tag (amino acids 58-687) was codon optimized for expression in *E. coli* and synthesized by Genscript. The gene was cloned into a pET28a vector to afford a TEV-protease cleavable N-terminal $His_6$-tagged translational fusion protein. *E. coli* BL21 (DE3) cells were transformed with pET28a-ScILV2 and transformants were selected on LB agar plates supplemented with 50 µg/mL kanamycin. A single colony was used to inoculate 7 mL of LB supplemented with kanamycin and cultures were grown for 18-20 h at 37 ˚C and 180 rpm. The entire overnight culture was used to inoculate 750 mL of kanamycin-supplemented LB in baffled 2 L flasks and cultures were grown at 37˚C and 180 rpm until $OD_{600}$ reached 0.6. Cultures were then placed on ice for 15 min before *Sc*ILV2 expression was induced with the addition of 0.4 mM IPTG. Cultures were grown for an additional 18 h before cells were harvested by centrifugation at 4000 x g for 15 min. Cell pellets were resuspended in 35 mL of ice cold lysis buffer (50 mM Tris pH 7.5, 500 mM NaCl, 5% glycerol, 5 mM imidazole) supplemented with 1 mg/mL lysozyme and complete EDTA-free protease inhibitor cocktail (Roche). Cell suspensions were lysed by sonication for 5 min (2 sec on, 4 sec off; 35% power) using a Qsonica sonicator and insoluble debris was removed by centrifugation at 20,000 x g for 30 min at 4˚C. The soluble fraction was applied to 2 mL of TALON fast flow resin (Cytiva) pre-equilibrated with lysis buffer. The resin was washed with 20 column volumes of lysis buffer and bound protein was eluted in 30 mL of elution buffer (200 mM $KH_2PO_4$ pH7.5, 10 µM FAD, 0.5 mM TCEP, 125 mM imidazole, 10% glycerol). Protein was concentrated in a 30 kDa MWCO Amicon centrifugal filter (Millipore) and subjected to size exclusion chromatography with a superdex 16/600 200 pg column (Cytiva) equilibrated with 200 mM $KH_2PO_4$ pH 7.5, 10 µM FAD, 0.5 mM TCEP, 10% glycerol. Fractions containing *Sc*ALS were pooled and concentrated to 10-20 mg/mL in a 30 kDa MWCO Amicon centrifugal filter (Millipore). Purity was assessed by coomassie-stained SDS-PAGE and protein concentration was determined by absorbance at 280 nm and Bradford.

**Expression and purification of *Af*ALS**

*A fumigatus* ILV2 (the gene for *Af*ALS) lacking the N-terminal mitochondrial-localization tag (amino acids 112-752) was codon optimized for expression in *E. coli* and synthesized by Genscript. The gene was cloned into a pET28a vector to afford a TEV-protease cleavable N-terminal $His_6$-tagged translational fusion protein. *E. coli* BL21 (DE3) cells were transformed with pET28a-*Af*ILV2 and transformants were selected on LB agar plates supplemented with 50 µg/mL kanamycin. A single colony was used to inoculate 50 mL of LB supplemented with kanamycin and cultures were grown for 18-20 h at 37˚C and 180 rpm. The entire overnight culture was used to inoculate 1 L of kanamycin-supplemented TB in baffled 2 L flasks and cultures were grown at 37˚C and 180 rpm until $OD_{600}$ reached 1. Cultures were then placed on ice for 15 min before *Af*ALS expression was induced with the addition of 0.3 mM IPTG. Cultures were grown for an additional 20 h before cells were harvested by centrifugation at 4000 x g for 15 min. Cell pellets were resuspended in 35 mL of ice cold lysis buffer (50 mM $K_2HPO_4$, pH 8.0, 300 mM NaCl, 5 mM imidazole, 10 µM FAD) supplemented with 1 mg/mL lysozyme and complete EDTA-free protease inhibitor cocktail (Roche). Cell suspensions were lysed by sonication for 10 min (2 sec on, 5 sec off; 45% power) using a Qsonica sonicator and insoluble debris was removed by centrifugation at 20,000 x g for 30 min at 4 ˚C. The soluble fraction was applied to 2 mL of TALON fast flow resin (Cytiva) pre-equilibrated with lysis buffer. The resin was washed with 20 column volumes of lysis buffer and bound protein was eluted in 30 mL of elution buffer (500 mM $K_2HPO_4$, pH 8, 10 µM FAD, 0.5 mM TCEP, 300 mM imidazole, 10% glycerol). Protein was concentrated in a 30 kDa MWCO Amicon centrifugal filter (Millipore) and subjected to size exclusion chromatography with a superdex 16/600 200 pg column (Cytiva) equilibrated with 300 mM $K_2HPO_4$, pH 8, 10 µM FAD, 0.5 mM TCEP. Fractions containing *Af*ALS were pooled and concentrated to 10-20 mg/mL in a 30 kDa MWCO Amicon centrifugal filter (Millipore). Purity was assessed by coomassie-stained SDS-PAGE and protein concentration was determined by absorbance at 280 nm and Bradford.

**Forward genetic screening of 1**

Ethyl methanesulfonate (EMS) mutagenesis was performed on *S. cerevisiae* to facilitate the raising of resistant clones. Briefly, overnight yeast cultures were grown in YPD broth at 30°C with shaking at 180 rpm. Cells were harvested by centrifugation, washed once with sterile water and twice with 0.1 M $NaH_2PO_4$, and $10^8$ cells addedre to 1.5 mL eppendorf tubes containing 1.7 mL 0.1 M $NaH_2PO_4$. Mutagenesis was initiated by addition of 25 µL EMS to each tube followed by incubation for 1 h at 30°C with agitation, after which the reaction was quenched with 8 mL 5% sodium thiosulfate. Mutagenized cells were washed and resuspended in sterile water to prepare for spreading on selection plates.

Selection plates were prepared by spreading 30 µL of DMSO containing increasing concentrations of **1** dispensed into each well of a 6-well agar plate. Concentrations tested were 5 µM, 10 µM, 20 µM, 40 µM, and 80 µM. The solution was spread evenly over the wells and plates were dried overnight. Mutagenized cells were then plated onto drug-containing agar and incubated at 30°C for 4–5 days until resistant colonies appeared.

**Antifungal screening of 1**

Antifungal screening was performed by Micromyx, LLC (Kalamazoo, Michigan, USA). Test organisms were either reference strains from the American Type Culture Collection (ATCC; Manassas, VA), the Centers for Disease Control and Prevention (CDC; Atlanta, GA), or clinical isolates from the Micromyx (MMX) collection. Upon receipt at Micromyx, the isolates were streaked under suitable conditions onto agar medium appropriate to each organism. The yeast isolates were incubated for 24 hr at 35ºC and the fungi were incubated for 7 to 13 days at 35ºC, as appropriate. Colonies harvested from these growth plates were resuspended in the appropriate medium containing a cryoprotectant. Aliquots of each suspension were then frozen at -80ºC.

Prior to testing, the yeast isolates were streaked from frozen vials onto Sabouraud Dextrose Agar (SAB; Becton Dickenson [BD]/BBL; Sparks, MD, Lot No. 2228660) and were incubated at 35°C for 24 hr. Additional fungal isolates were streaked from frozen vials onto potato dextrose agar (BD/BBL; Sparks, MD, Lot No. 2046481) and incubated under optimal conditions for growth.

All MIC measurements were performed in RPMI1640 from Hyclone Laboratories (Logan, UT; Cat. No. SH3001104; Lot No. AC10257966A) buffered with MOPS to pH-7 from Millipore (Billerica, MA; Cat No. 475898; Lot No. 3462216).

MIC values were determined using a broth microdilution procedure described by CLSI. [4,5].

Automated liquid handlers (Multidrop 384, Labsystems, Helsinki, Finland; Biomek 3000 and Biomek FX, Beckman Coulter Fullerton CA) were used to conduct serial dilutions and liquid transfers.

Solutions of **1** were added in an 11 point 2x serial dilution in DMSO with a maximum tested concentration of 32 µg/ml in the culture medium. Drug solution was added at 1% v/v into 200 µl of culture in all wells.

A standardized inoculum of each test organism was prepared in appropriate media per CLSI methods.[4,5] The plates were then inoculated with 10 µL of the diluted inoculum using the Biomek 3000 from low to high drug concentration, resulting in a final concentration of approximately 0.5 to 2.5 x $10^3$ CFU/mL for yeast and 0.2 to 2.5 x $10^4$ CFU/mL for the other fungi. An un-inoculated plate was incubated for the purpose of assessing solubility of the drug in the test medium.

The plates were stacked 3 to 4 high, covered with a sterile lid on the top plate, and incubated aerobically at 35ºC for 24, 48, and 72 h. For yeast, both the MIC based on complete inhibition and 50% inhibition were reported at 24 and 48 h. For other fungal isolates, the MIC values were recorded at 24 and 48 hr and were based on complete inhibition and 50% inhibition.

**Intact protein LCMS analysis**

10 µM of recombinant *S. cerevisiae* ALS was incubated with 10 µM compound **1** in buffer (50 mM Tris pH 7.5, 100 mM KCl, 10 uM FAD, 1 mM TPP, 10 mM $MgCl_2$ (with 2.5% DMSO from compound)) at 22 °C for 30 min. Unreacted **1** was removed by passing the reaction mixture through a 30 kDa MWCO filter (Millipore). The collected protein was then denatured and reduced by incubation with 6 M urea and 10 mM DTT at 37ºC for 1 h followed by 20 mM iodoacetamide alkylation for 30 min in dark. This reaction mixture then underwent buffer exchange into 100 mM Tris-HCl pH 8.0 and 10 mM $CaCl_2$ buffer using a 30 kDa MWCO filter. The protein was then incubated with sequencing grade chymotrypsin (Promega) (15:1,w/w) for 2 h at 37 ºC.

After digestion, the peptide mixture was immediately analysed by LC-MS/MS using a Waters I-class UPLC equipped with an ACQUITY UPLC BEH C18 column (2.1x50 mm, 1.7 µm, Waters) at 50ºC coupled to a Waters Synapt G2Si MS. The sample was injected and subjected to an LC gradient starting from 0.5% mobile phase B (ACN with 0.1% formic acid) for 5 min with linear increase to 40% B over 35 min. Mobile phase A was 0.1% formic acid water. Flow rate was maintained at 0.3 ml/min. MS/MS data was acquired in positive ionization mode with data dependent MS/MS acquisition. Peptide mapping was performed with PEAKs Studio 8.0

(Bioinformatics Solutions Inc.) followed by manual review to identify peptides putatively modified by covalent addition of **1**.

##### Protein expression and Purification for Cryo-EM

*Af*ALS(residues S112-K752) with an N-terminal $His_6$ tag followed by a TEV protease cleavage site was cloned in a pET28a vector and transformed into *E. coli* BL21 (DE3) cells for protein expression. These expression stains were grown at 37°C in Terrific Broth to mid-exponential phase ($OD_{600}$ = 0.6) followed by induction with 0.4 mM isopropyl β-D-1-thiogalactopyranoside (IPTG) at 16°C for 18 hours.

Cells were harvested and resuspended in buffer (50 mM $K_2HPO_4$, pH 8.0, 300 mM NaCl, 10 μM FAD, 5 mM imidazole). The slurry was then lysed by sonication and clarified by centrifugation. The supernatant was then applied to a TALON column and washed with wash buffer (50 mM $K_2HPO_4$, pH 8.0, 300 mM NaCl, 10 μM FAD, 10% glycerol, 0.5 mM TCEP). *Af*ALS was then eluted with elution buffer (500 mM $K_2HPO_4$, pH 8, 10 μM FAD, 0.5 mM TCEP, 200 mM imidazole, 10% glycerol). The eluate was further purified by size exclusion chromatography (Superdex 200 Increase 10/300 GL, Cytiva) with formulation buffer. Purity of the proteins was analyzed by SDS-PAGE after each step and the purified proteins were concentrated and stored at -80°C.

##### Cryo-EM sample preparation, data collection and processing

The *Af*ALS complex with **1** was prepared by mixing 1mg/mL *Af*ALS with 0.7 mM **1** followed by incubation on ice for 0.5 h. Both Apo and **1** bound-*Af*ALS were diluted to 0.5mg/mL with formulation buffer for cryoEM grid freezing. 3 μl of sample was applied to Ultra Au 300 1.2/1.3 (Quantifoil, Inc) pretreated by glow discharging. The grid was blotted at 4°C with 100% humidity and plunge-frozen in liquid ethane using FEI Vitrobot Mark IV (Thermo Fisher). The grids were stored in liquid nitrogen before screening and data collection.

Images were acquired on a Krios G4 Cryo-Transmission electron microscope (Thermo Fisher) equipped with Falcon 4i Direct Electron Detector in counting mode, at a calibrated magnification of 165,000X with the physical pixel size corresponding to 0.743 Å. Detailed data collection statistics have been indicated in **Table S2**. Motion correction of each micrograph, contrast transfer function (CTF) estimation, particle picking, 2D classification, ab initio 3D reconstruction, heterogeneous refinement, homogeneous refinement,, and local refinement were carried out by standard pipeline in cryoSPARC[6], 3D classification and 3D auto-refine were carried out in RELION.[7] For **1**-bound *Af*ALS, 3,065,801 initial particles were extracted from 6014 movies, and after iterative 2D classification 411,943 particles were selected for heterogeneous refinement, from which a class of 263,024 particles was further selected for homogeneous refinement. The particles were then imported to RELION for RELION SIRM reconstruction. The generated model, which still had preferred orientation, was then further processed by 3D classification, and a subset of 125,472 particles was selected for 3D auto-refine. The particles and volume were then imported back to cryoSPARC for final polishing. Homogeneous refinement and local refinement plus CTF refinement generated a final reconstruction with C2 symmetry at a resolution of 2.36 Å, based on the Fourier shell correlation (FSC) cutoff at 0.143 between the two half maps. The Apo *Af*ALS reconstruction was obtained through the similar image processing workflow, and a final reconstruction with C2 symmetry at a resolution of 2.76 Å was obtained from 64,577 particles. The final map was of excellent quality consistent with the resolution (**Figure S10**).

**Model building and refinement**

A homogenous *Af*ALS structure was obtained by SWISS model[8] using the *Sc*ALS crystal structure (PDB ID 5IMS[9]) as the template. Then the *Af*ALS model was docked into the cryo-EM map using Chimera.[10] Both Apo and **1**-bound *Af*ALS structures were further built and refined in COOT.[11] The structure of **1** was built by eLBOW in PHENIX and docked into the map for further refinement. Both structures were further refined in PHENIX[12] with secondary structure restraints.[13] The final model with good geometry and fit to the map (**Figure S10B**) was validated using the comprehensive cryo-EM validation tool implemented in PHENIX[12] (**Table S2**). All structural figures were generated using PyMol[14] and ChimeraX.[15]

**Construction of *A. fumigatus* AF293 *ilv2Δ* and the reconstituted strain**

The *A. fumigatus ilv2Δ* (Afu3g10310) mutant was generated in the reference strain AF293[16] using CRISPR-Cas9 technology as described previously.[17] Briefly, two crRNAs adjacent to PAM motifs (g201: AATTGGCCTTTGGCCGCGCT and g202: AGTGATCATGTAGACGCTTT) that target Cas9 cleavage in the upstream and downstream of the ilv2 gene were designed. On the day of transformation, crRNA-tracrRNA duplex was generated by mixing equimolar concentrations of each crRNA with tracrRNA in duplex buffer, boiling for 5 minutes, and then allowing them to anneal by cooling to room temperature. This resulting duplex was combined with the Cas9 enzyme to generate the Cas9-RNP complex. We generated the repair construct by amplifying the hygromycin expression cassette with 35bp overhang primers (RAC 7458 and RAC 7459, **Table S4**) adjacent to the Cas9 targeted sites. Next, the Cas9-RNP complex and 2 µg of repair construct were transformed into AF293 protoplasts.[18] The transformants were selected on glucose minimal media (GMM)[19] with 1.2 M soribitol (SMM), hygromycin (175 µg/mL), and 5 mM each of isoleucine and valine.

Replacement of *ilv2* was confirmed using junction PCRs (RAC 7462/RAC6355, and RAC6562/RAC7463, **Table S4**) and by checking for the absence of the open reading frame (using RAC7460 and RAC7461, **Table S4**) in the mutant. The mutant's auxotrophy was confirmed by its ability to grow only on GMM supplemented with 5mM each of isoleucine and valine (**Figure S8**).

For the generation of the reconstituted strain, the *ilv2* locus (including 949bp of the promoter and 652bp of the terminator/3'UTR region) was amplified using primers RAC7498 and RAC7499 (**Table S4**), and the resulting product was integrated into a plasmid containing pyrathimaine (*ptrA*) selection marker using HiFi (NEB), such that the selection marker lies downstream of *ilv2*. The repair construct was amplified from the plasmid using RAC7502/RAC7503 (**Table S4**), which included a 35-bp overhang on each primer flanking the Cas9 cut site at the *aft4* safe haven site.[20] The sequence for the crRNA that targets the *aft4* site is g65 - TCTCCTTCATAAGCGACCAG. The Cas9-RNP complex and 2 µg of repair construct were transformed into Afu3g10310 protoplasts, and the transformants were selected on SMM containing 100 µg/L pyrithiamine hydrobromide. Integration of the gene at the *aft4* site was confirmed by junction PCRs (RAC6583/RAC7503, RAC6558/RAC6584, **Table S4**) and a PCR amplifying a region within the ORF of the gene (RAC7460 and RAC7461, **Table S4**).

To confirm the restoration of gene expression, RT-qPCR was performed on four candidates, using AF293 and Afu3g10310 as controls. Briefly, $10^5$ conidia/ml of each strain were cultured in GMM containing 5 mM each of isoleucine and valine. After 16 hours, the adhered biofilms were scraped and resuspended in Trisure reagent. All samples were bead-beaten for 1 minute, and manufacturer instructions were followed thereafter. 500 ng of RNA was used for cDNA synthesis. cDNA was then diluted 1:4 with water and 2 µL per reaction was used for qPCR. The gene expression was normalized to housekeeping genes (tubulin and actin), and fold change was calculated relative to wild-type.

##### Assessing the pathogenicity of *A. fumigatus* AF293 variants in a mouse model of invasive aspergillosis

All strains used in the animal experiment were grown on GMM supplemented with 5 mM each of isoleucine and valine for 3 days at 37°C, 5% $CO_2$. On the day of inoculation, the conidia were collected in 0.01% v/v Tween 80 in water, washed, counted, and then resuspended in PBS for inoculation.

Outbred 20-24 g female CD-1 mice (Charles River Laboratory, Raleigh, NC, USA) were housed in autoclaved cages (3-4 per cage) with HEPA-filtered air, food, and autoclaved water available *ad libitum*. For immunosuppression, the mice were subcutaneously administered 40 mg/kg Kenalog-10 (triamcinolone acetonide; Bristol-Myer Squibb, Princeton, NJ).[21] After 24 hrs, the mice were intranasally administered with $2 \times 10^6$ conidia of each strain in 40 µl sterile PBS or PBS alone and monitored for survival.

To determine the fungal burden in the lungs, the mice were sacrificed 72 hours after inoculation. The lungs were harvested, flash frozen, and the relative fungal burden was assessed through qPCR quantitation of *A. fumigatus* 18S rDNA, as previously described.[22,23]

##### Ethics statement

The animal protocol was approved by the Institutional Animal Care and Use Committee (IACUC; federal-wide assurance number A3259-01) at Dartmouth College. The experiments were carried out with strict adherence to the recommendations in Guide for the Care and Use of Laboratory Animals.[24]

##### Spectroscopic Analyses

HPLC purifications were performed using either a Biotage Selekt Flash Chromatography, a Thermo Vanquish Flex HPLC or an Agilent 1260 HPLC. HPLC-HRMS analyses were acquired using a Thermo Vanquish HPLC system coupled to a Thermo Orbitrap Exploris 120 high-resolution mass spectrometer at 120 K resolution for MS1 and 17.5 K resolution for MS/MS. All chromatographic methods used 0.1% aqueous formic acid (A) and acetonitrile with

0.1% formic acid (B) as mobile phases unless described otherwise. For elucidation of chemical structures, 1D and 2D NMR spectra were obtained on Bruker AV600 spectrometer ($^1$H 600 MHz, $^{13}$C 150 MHz) equipped with a BBO cryoprobe. Chemical shifts were referenced using the solvent peak or TMS (δ 0).

### Isolation of compounds:

##### Isolation of compounds 1, 13, 14, 15, 16, 17 and 19

*A. nidulans* expressing *sdtABCDFGH* were cultured in 2 L flasks containing 1 L LMM media (12 L total). Each liter of culture was extracted by adding a sealed nylon bag containing 10 gm each of Amberlite XAD7HP and Diaion HP20 resin and shaken at 150 rpm for 16 h at room temperature. Resin bags from each flask were collected using long tweezers, combined and extracted with ethyl acetate for 3 h. The organic extract was filtered and dried. The crude extract was subjected to flash chromatography using a C18 column (120 g, pore size), with a flow rate of 50 mL/min, 5% solvent B was used for 1.5 column volumes (CV), a gradient of 5-30% solvent B for 3.5 CV, a gradient of 30-100% solvent b for 3 CV, and a wash of 100% solvent B for 2 CV. Target fractions containing metabolites of interest were identified by HPLC-HRMS using a linear gradient of 5-100% B in 10 minutes. Fractions containing compounds **1**, **13** and **19** were combined and further fractionated by semi-preparative HPLC (XBridge BEH Shield RP18 OBD, 10 × 250 mm, 10μm) and an isocratic method with 43%B for 60 mins, yielding **1** (0.5 mg/L), **13** (0.1 mg/L) and **19** (0.1 mg/L). Fractions containing **14** and **16** were combined and further fractionated by preparative HPLC using a C18 column (Eclipse XBD-C18 21.2 x 250 mm, 7 um), with a gradient of 50-100%B over 34 min (0-5 min, 50%B; 5-25 mins, 50-60% B; 25-27 mins, 60-100%B ; 27-34 min, 100%B) at 25 mL/min, yielding **14** (3.0 mg/L) and **16** (0.5 mg/L). Fractions containing **15** were combined and further fractionated using the same C18 column, with a gradient of 50-100%B over 27 min (0-5 min, 50%B; 5-20 mins, 50-62% B; 20-22 min, 62-100%B ; 22-27 min, 100%B) at 25 mL/min, yielding **15** (1.0 mg/L). Fractions containing **17** were combined and further fractionated also using the same C18 column with a gradient of 50%-100%B over 35 min (0-5 min, 50%B; 5-25 min, 50%-68%B; 25-28 min, 68%-100%B; 28-35 min, 100%B) at 25 mL/min, yielding 17 (0.1 mg/L)

##### Isolation of compound 10

*Aspergillus nidulans* expressing *sdtBDF* were cultured in 2 L flasks containing 1 L LMM media (12 L total). Each liter of culture was extracted by adding a sealed nylon bag containing 10 grams each of Amberlite XAD7HP and Diaion HP20 resin and shaken at 150 rpm for 16 h at room temperature. Resin bags from each flask were collected using long tweezers, combined and extracted with ethyl acetate for 3 hours. The organic extract was filtered and dried. The crude extract was subjected to flash chromatography using a C18 column (120 g, pore size) with a flow rate of 50mL/min, 5% solvent B was used for 1.5 column volumes (CV), a gradient of 5-30% solvent B for 3.5 CV, a gradient of 30-100% solvent b for 3 CV, and a wash of 100% solvent B for 2 CV. Target fractions containing metabolites of interest were identified by LC-HRMS using a linear gradient of 5-100% B in 10 minutes, Fractions were further fractionated by preparative HPLC using a C18 column (Eclipse XBD-C18 21.2 x 250 mm, 7 um),with a gradient of 70-100%B over 35 minutes (0-5 min, 70%B; 5-25 mins, 70-90% B; 25-27 mins, 90-100%B ; 27-35 min, 100%B) at 25mL.min$^{-1}$, yielding **10** (2 mg/L)

##### Isolation of compounds 11 and 12

For isolation of **11**, spores of the *A. nidulans* expressing *neoBDFH* were inoculated into 5 L CD-ST agar media and grown for 4 days at 28°C. The culture was exhaustively extracted with EtOAc, and the organic solvent was evaporated to dryness under vacuum. The resulting crude extract was fractionated by flash column chromatography subjected to an acetonitrile-water gradient (5:95 to 100:0). Fractions containing **11** were purified by semi-preparative HPLC using a C18 column (AR-II) and a gradient of 70-95% B over 30 min to yield **11** (0.4 mg/L).
For isolation of **12**, spores of the *A. nidulans* expressing *neoABDFH* were inoculated into 5 L CD-ST agar media and grown for 4 days at 28°C. The culture was exhaustively extracted with EtOAc, and the organic solvent was evaporated to dryness under vacuum. The resulting crude extract was fractionated by flash column chromatography subjected to an acetonitrile-water gradient (5:95 to 100:0). Fractions containing **12** were purified by semi-preparative HPLC using a C18 column (AR-II) and a gradient of 70-95% B over 30 min to yield **12** (0.4 mg/L).

##### Structure elucidation of isolated compounds

Purified HB-35018 (**1)** was isolated as a off-white solid. HRMS m/z 386.2328 for the $[M+H]^+$ ion was consistent with a molecular formula $C_{23}H_{31}NO_4$. NMR data was acquired for **1** in $CD_3OD$ (**Table S5**, **Figures S11-S16**). $^1H$, $^{13}C$ and $^1H$-$^{13}C$ HSQC showed the presence of 23 carbons and 29 observable protons. Of the 23 carbons, seven are sp$^2$ hybridized, among them 2 carbonyls at δ 209.0 and δ 211.0, one carboxyl at δ 173.5 and four olefinic carbons at δ 126.1, δ 130.6, δ 130.7 and δ 137.5. The remaining 16 (sp$^3$) carbons were identified as one aminomethine, one oxymethylene, four methines, five methylenes, three methyls and two quaternary carbons. COSY cross-peaks allowed the determination of spin systems for H-2-H-3, H-5-H-9, H-11-$H_3$-14 and H-4’-$H_3$-7’. A weak COSY correlation between H-3-and H-8 allowed the establishment of the bicyclic decalin scaffold and suggested a *cis*-decalin configuration based on J coupling constants. HMBC correlations from $H_3$-15 to C-2’, C-2, C-3 and C-4, and from $H_3$-16 to C-9, C-10 and C-11 allowed the exact placement of methyl groups. The HMBC correlation from H-11 to C-1, from $H_3$-15 to C-2’ and from H-2 to both C-1 and C-2’ provided the basis for establishing a rare hexahydro-acenaphthene-1-one tricyclic core. HMBC correlations from $H_2$-5’ to C-3’ and C-4’, and from H-4’ to C-1’ and C-3’ indicated an alkyl tetramic acid scaffold. While there is no direct NMR evidence, the only possible connection between these two ring systems would be through a carbocyclic spiro formation at the quaternary carbon C-2’. The observation of NOESY correlation among H-3, H-8 and $H_3$-15, as well as between H-2 with both Ha-12 and Ha-13 allowed the assignment of relative configurations for C-2, C-3, C-4, C-8 and C-11. The absolute configuration was determined by X-ray analysis, confirmed as 2R, 3R, 4R, 8R, 11R, 2’R, 4’R.
Compounds **13** and **19** were also obtained as off-white solids. HRMS indicated the same molecular formula of **1**, $C_{23}H_{31}NO_4$. NMR data indicated a scaffold identical to **1** for both compounds. The main differences observed were NOE correlations from H-4’ to H-5 and $H_3$-15

in both **13** and **19**. This spatial proximity could be attributed to a direct stereochemical change at the C-4’carbon, or by an inversion of the stereocenter at the spirocyclic carbon C-2'. Careful analysis of the $^{1}H$ NMR shifts in **13** and **19** revealed that while there is minimal impact to other proton chemical shifts in **19**, there is a noticeable upfield shift for H-5 and downfield shift for $H_3$-15 for compound **13**. The absolute configuration of **1** indicates that the keto carbonyl at C-3’ faces the same side as $H_3$-15, while the amide carbonyl at C-1’stacks with H-5. An inversion in the spirocyclic carbon stereocenter changes the overlap between the carbonyl carbons C-1’and C-3 and the protons at H-5 and $H_3$-15. As such, we have designated compound **13** as the 2'S isomer of **1**, and **19** as the 4'S isomer of **1**.

Unfortunately, despite extensive effort, the 2'S, 4'S isomer of **1** was not obtained in sufficient amount for NMR characterization. However, a set of four intermediates (**14-17**) were isolated, representing all four possible stereoisomers. They were all isolated as off-white solids and showed a HRMS m/z of 400.2118 as the $[M+H]^+$ ion, indicating a molecular formula of $C_{23}H_{29}NO_5$. As expected, strong agreement in 1D chemical shifts and 2D correlations for all compounds were observed, similar to **1** and its isomers, with the major difference identified as the presence of a carboxylic acid at C-7’. Analysis of the chemical shifts of H-5, $H_3$-15 and H-4’and the presence or absence of NOE among them allowed the identification of **14** as the C-7’COOH analog of **19** **(**2'S, 4'R), **16** as the C-7’COOH analog of **1** **(**2'S, 4'S), **17** as the C-7’COOH analog of **13** **(**2'R, 4'S)**,** and **15** as the 2'R, 4'R isomer.

### Supporting Tables

**Table S1:** Allele frequencies of all mutations observed in *S. cerevisiae* ALS from clones raised to be resistant to **1**.

| Mutation (S.cerevisiae) | Number of clones expressing this mutation | Frequency | Residue number in *Afl*LV2 |
|---|---|---|---|
| A117V | 1 | 4.3% | A176 |
| V191L | 1 | 4.3% | V250 |
| P192S | 3 | 13.0% | P251 |
| P192L | 15 | 65.2% | P251 |
| A200V | 1 | 4.3% | S259 |
| W586R | 1 | 4.3% | W648 |
| S596F | 1 | 4.3% | S658 |

**Table S2:** Cryo-EM data collection, refinement and validation statistics

| Data collection and processing | apo AfALS | *AfALS + 1* |
|---|---|---|
| Magnification | 165,000 | 165,000 |
| Voltage (kV) | 300 | 300 |
| Electron exposure (e–/Å2) | 65 | 65 |
| Defocus range (μm) | -1 - -2.4 | -1 - -2.4 |
| Pixel size (Å) | 0.743 | 0.743 |
| Symmetry imposed | C2 | C2 |
| Initial particle images (no.) | 3,284,254 | 3,065,801 |
| Final particle images (no.) | 64,577 | 125,472 |
| Map resolution (Å)<br>FSC threshold | 2.76<br>0.143 | 2.36<br>0.143 |
| Map resolution range (Å) | 2.5 – 3.5 | 2.0-2.7 |
| **Refinement** | | |
| Initial model used (PDB code) | 5IMS | 5IMS |
| Model resolution (Å)<br>FSC threshold | 2.74<br>0.143 | 2.55<br>0.143 |
| Model resolution range (Å) | 2.74 | 2.55 |
| Model composition<br>Nonhydrogen atoms<br>Protein residues<br>Ligands | <br>9122<br>1164<br>4 | <br>9092<br>1154<br>6 |
| *B factors (Å2)*<br>*Protein*<br>*Ligand* | <br>56.68<br>52.63 | <br>83.69<br>90.87 |
| R.m.s. deviations<br>Bond lengths (Å)<br>Bond angles (°) | <br>0.009<br>1.079 | <br>0.002<br>0.635 |
| Validation<br>MolProbity score<br>Clashscore<br>Poor rotamers (%) | <br>2.54<br>20.06<br>4.18 | <br>1.75<br>7.12<br>1.48 |
| Ramachandran plot<br>Favored (%)<br>Allowed (%)<br>Disallowed (%) | <br>96.02<br>3.98<br>0.00 | <br>96.51<br>3.49<br>0.00 |

**Table S3:** Physical chemical and pharmacokinetic parameters for **1**

| | | |
|---|---|---|
| **Calculated properties** | MW \| ClogP \| tPSA (Å$^2$) | 386 \| 2.7 \| 83 |
| **Solubility** | PBS kinetic solubility, 5% DMSO (μM) | 260 |
| **Buffer stability** | PBS pH 7.4 $t_{1/2}$ (min) | 141 |
| **Plasma stability** | Plasma $t_{1/2}$ (min) Mouse \| Human | 88 \| 79 |
| **Liver microsome stability** | Liver microsome $t_{1/2}$ (min) Mouse \| Human | 3 \| 3 |

**Table S4:** Primers used in the construction and validation of *A. fumigatus ilv2Δ* (Afu3g10310)

| Primer name | Binds to | sequence |
|---|---|---|
| RAC7458 | Repair forward | ACATCGGAGTTCGCTGTCCAATAATGGCGTTAAGGTAC CGGTCGCCTCAAACAATGCTCT |
| RAC7459 | Repair reverse | GAAAGATGAAAGATTAGTAAGCAACGTTGCAAAGCCGG TCTGAGAGGAGGCACTGATGCG |
| RAC7460 | ORF forward | GAGTCATCCCTCGCTTACCA |
| RAC7461 | ORF reverse | GCAATCTCGAAAGCCTCCTG |
| RAC7462 | 5' utr forward | AGGTCCATACTTGTGCTCTGT |
| RAC7463 | 3' utr reverse | AATGTCCGACAGAACCAGGG |
| RAC6355 | Hygromycin reverse | cagaaagaacgcatccatca |
| RAC 6562 | Hygromycin forward | gactgaggaatccgctcttg |
| RAC7498 | ilv2p forward | AAGAACCGGTGAATTTGGCT |
| RAC7499 | Ilv2t reverse | CCGCGGCTTTCAAGGATTT |
| RAC7502 | Recon repair fwd | TCAGGTGCATCTTCCAGTTCTGGATATAGCATATAGATCT ATCTACCCCGCTTCTGATGC |
| RAC6321 | Recon repair rev | TTTGGCCTCCATACTCCCCTGATCTCAATCCAGTTGAGA Acggccagtgccaagctctag |
| RAC6583 | *aft4 site fwd* | GCCTCGATAACTGCCTTCAC |
| RAC7503 | *ilv2 rev* | CTTTCTGTGTGCGGCCTATC |
| RAC6584 | *aft4 site rev* | TCAGGCCGGGTTATAAGAGA |
| RAC6558 | *ptrA fwd* | gcatgaaccggatttgtctt |
| RAC007 | tubulin fwd | ATAATGTTCAGACCGCCCTCTGCT |
| RAC008 | tubulin rev | GACGGATGTGGAATTGCCCACAAA |
| RAC158 | actin fwd | TCACTGCCCTTGCTCCCTCGTC |
| RAC159 | actin rev | GCACTTGCGGTGAACGATCGAA |

**Table S5:** NMR spectroscopic data for **1**

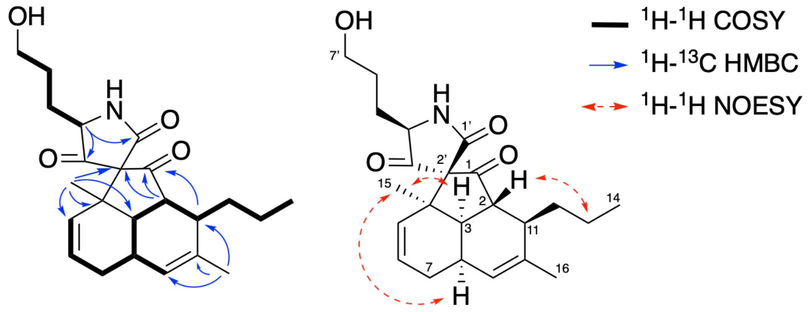

| | Found (MeOD) | |
|---|---|---|
| Position | $\delta_C$ | $\delta_H$, m(J in Hz)1 |
| 1 | 210.2 | - |
| 2 | 49.6 | 2.59, dd(14.1,10.4) |
| 3 | 46.0 | 2.30. ddd(14.2, 4.3, 1.7) |
| 4 | 47.1 | - |
| 5 | 131.1 | 5.80, ddt (10.4, 3.1, 1.6) |
| 6 | 125.0 | 5.63, ddd(10.4, 5.3, 2.4) |
| 7 | 29.0 | Ha: 2.36, dddd (18.8, 7.0, 5.3, 1.7)/ Hb: 1.73, ddt(18.9, 10.1, 2.7)/ |
| 8 | 29.6 | 2.50, m |
| 9 | 130.8 | 5.75, d(2.8) |
| 10 | 137.4 | - |
| 11 | 42.7 | 2.43, m |
| 12 | 32.0 | Ha: 1.93, m/ Hb: 1.59, m |
| 13 | 17.8 | Ha: 1.24, m/ Hb: 1.03, m |
| 14 | 14.9 | 0.89, t(7.2) |
| 15 | 24.7 | 1.18, s |
| 16 | 20.8 | 1.66, s |
| 1' | 172.1 | - |
| 2' | 76.2 | - |
| 3' | 209.9 | - |
| 4' | 64.0 | 3.92, dd(8.3,5.2) |
| 5' | 30.1 | 1.60, m/1.81, m |
| 6' | 29.7 | 1.65, m |
| 7' | 62.2 | 3.58, t(6.1) |

In MeOD, 600 MHz for $^1H$ and 150 MHz for $^{13}C$ NMR; Chemical shifts are reported in ppm. All shifts are determined by $^1H$, $^{13}C$, COSY, HSQC, HMBC and NOESY correlations.

**Table S6:** NMR spectroscopic data for **10**

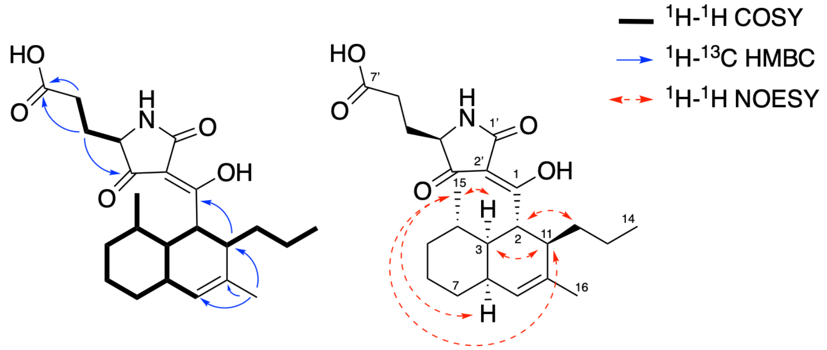

| Position | δC | δH, m(J in Hz) |
|---|---|---|
| 1 | 195.1 | - |
| 2 | 47.9 | 3.8, m |
| 3 | 52.5 | 1.18 ,m |
| 4 | 40.0 | 1.38 ,m |
| 5 | 38.0 | Ha: 1.69, m/ Hb: 1.02, m |
| 6 | 27.2 | Ha: 1.73, m/ Hb: 1.41, m |
| 7 | 34.0 | Ha: 1.93, m/ Hb: 1.22, m |
| 8 | 41.6 | 1.75, m |
| 9 | 131.1 | 5.41, s |
| 10 | 138.3 | - |
| 11 | 43.0 | 2.34, m |
| 12 | 32.6 | Ha: 1.46, m/ Hb: 1.35, m |
| 13 | 23.1 | 1.35, m |
| 14 | 14.7 | 0.86, m |
| 15 | 21.1 | 0.84, m |
| 16 | 20.0 | 1.71, m |
| 1' | 177.1 | - |
| 2' | 102.9 | - |
| 3' | 197.3 | - |
| 4' | 62.2 | 3.91, m |
| 5' | 28.1 | Ha: 2.07, m/ Hb: 1.88, m |
| 6' | 29.8 | 2.34, m |
| 7' | 176.2 | - |

In MeOD, 600 MHz for $^1$H and 150 MHz for $^{13}$C NMR; Chemical shifts are reported in ppm. All shifts are determined by $^1$H, $^{13}$C, COSY, HSQC, HMBC and NOESY correlations.

**Table S7:** NMR spectroscopic data **for 11**

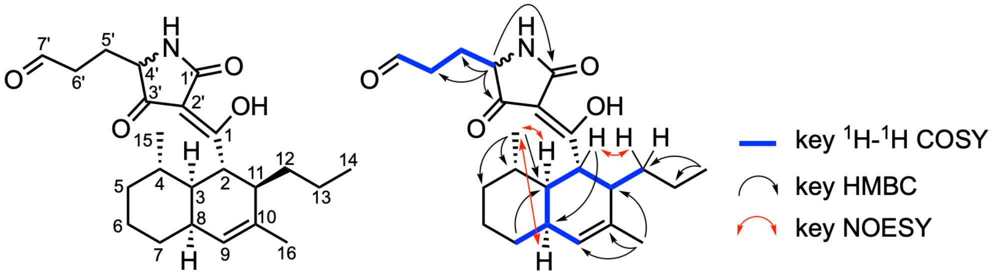

**11**

| **Position** | **δC** | **δH, m(J in Hz)** |
| --- | --- | --- |
| 1 | 192.8 | - |
| 2 | 40.7 | 4.07 m |
| 3 | 43.0 | 1.86 m |
| 4 | 31.2 | 1.66 m |
| 5 | 27.6 | 1.51 m |
| 6 | 20.7 | 1.40 m |
| 7 | 30.6 | 1.15 m |
| 8 | 32.6 | 2.25 m |
| 9 | 129.8 | 5.53 m |
| 10 | 134.7 | - |
| 11 | 43.3 | 2.52 m |
| 12 | 32.6 | Ha: 1.51, m/ Hb: 1.35, m |
| 13 | 18.0 | Ha: 1.43, m/ Hb: 1.03, m |
| 14 | 14.8 | 0.80 m |
| 15 | 19.5 | 1.00 m |
| 16 | 21.2 | 1.63 s |
| 1' | 176.3 | - |
| 2' | 104.8 | - |
| 3' | 196.3 | - |
| 4' | 61.7 | 3.88 m |
| 5' | 24.6 | Ha: 2.06, m/ Hb: 1.90, m |
| 6' | 39.1 | Ha: 2.49, m/ Hb: 2.44, m |
| 7' | 202.6 | 9.67 m |

In $CDCN_3$, 500 MHz for $^{1}H$ and 125 MHz for $^{13}C$ NMR; Chemical shifts are reported in ppm. All shifts are determined by $^{1}H$, $^{13}C$, COSY, HSQC, HMBC and NOESY correlations.

**Table S8:** NMR spectroscopic data for **12**

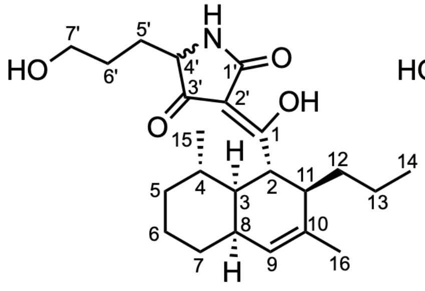

**12**

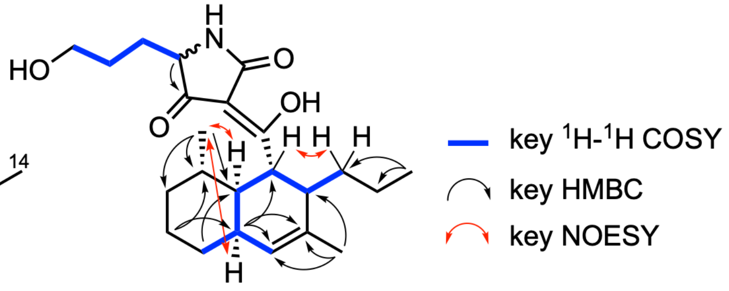

| Position | δC | δH, m(J in Hz) |
|---|---|---|
| 1 | 192.5 | - |
| 2 | 40.3 | 4.08 t (10.7) |
| 3 | 42.5 | 1.89 m |
| 4 | 30.8 | 1.64[a] |
| 5 | 27.4 | 1.54[a] |
| 6 | 20.4 | 1.42 m |
| 7 | 28.5 | 1.18 m |
| 8 | 32.4 | 2.24 m |
| 9 | 129.3 | 5.52 d (5.8) |
| 10 | 134.1 | - |
| 11 | 42.9 | 2.54 m |
| 12 | 32.2 | Ha: 1.49, m/ Hb: 1.21, m |
| 13 | 17.7 | Ha: 1.43, m/ Hb: 1.05, m |
| 14 | 14.8 | 0.81 t (7.1) |
| 15 | 19.4 | 1.00 d (7.2) |
| 16 | 21.2 | 1.63 s |
| 1' | 175.4 | - |
| 2' | 104.1 | - |
| 3' | 195.5 | - |
| 4' | 62.4 | 3.87 m |
| 5' | 30.2 | Ha: 1.95, m/ Hb: 1.69, m |
| 6' | 29.4 | Ha: 1.68, m/ Hb: 1.58, m |
| 7' | 62.7 | 3.66 m |

In $CD_2Cl_2$, 500 MHz for $^1H$ and 125 MHz for $^{13}C$ NMR; Chemical shifts are reported in ppm. All shifts are determined by $^1H$, $^{13}C$, COSY, HSQC, HMBC and NOESY correlations.

**Table S9:** NMR spectroscopic data for **13**

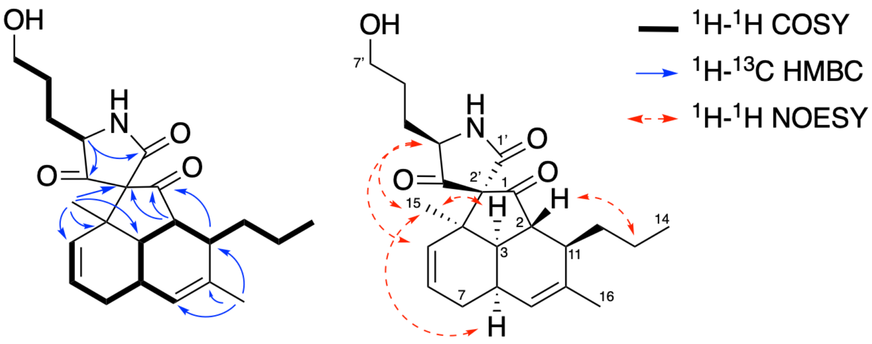

| | Found (MeOD) | |
|---|---|---|
| Position | $\delta_C$ | $\delta_H$, m(J in Hz) |
| 1 | 211.0 | - |
| 2 | 49.7 | 2.46, m |
| 3 | 46.0 | 2.34, m |
| 4 | 48.0 | - |
| 5 | 130.7 | 5.4 d(10.5) |
| 6 | 126.1 | 5.67 ddd(10.4, 5.3, 2.4) |
| 7 | 29.1 | Ha: 2.37, m/ Hb: 1.74, m |
| 8 | 29.9 | 2.49, m |
| 9 | 130.6 | 5.74, d (6.3) |
| 10 | 137.5 | - |
| 11 | 42.3 | 2.43, m |
| 12 | 32.0 | Ha: 1.93, m/ Hb: 1.6, m |
| 13 | 17.9 | Ha: 1.20, m/ Hb: 1.03, m |
| 14 | 14.9 | 0.89 t(7.2) |
| 15 | 24.2 | 1.30, s |
| 16 | 20.8 | 1.65, s |
| 1’ | 173.5 | - |
| 2’ | 76.2 | - |
| 3’ | 209.0 | - |
| 4’ | 64.8 | 4.03, t (5.8) |
| 5’ | 28.8 | Ha: 1.88, m/ Hb: 1.72, m |
| 06’ | 29.0 | Ha: 1.64, m/ Hb: 1.60, m |
| 7’ | 62.5 | 3.57, t (6.0) |

In MeOD, 600 MHz for $^1$H and 150 MHz for $^{13}$C NMR; Chemical shifts are reported in ppm. All shifts are determined by $^1$H, $^{13}$C, COSY, HSQC, HMBC and NOESY correlations.

**Table S10:** NMR spectroscopic data for **14**

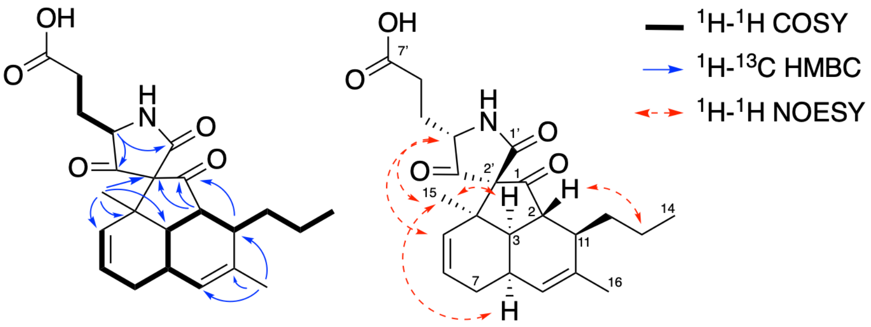

| | Found (MeOD) | |
|---|---|---|
| Position | $\delta_C$ | $\delta_H$, m(J in Hz) |
| 1 | 211.1 | - |
| 2 | 49.5 | 2.53. m |
| 3 | 46.0 | 2.26, dd(13.9, 1.5) |
| 4 | 47.9 | - |
| 5 | 130.6 | 5.70, d(10.6) |
| 6 | 125.5 | 5.65, ddd(10.4, 5.2, 2.2) |
| 7 | 29.0 | Ha: 2.34, t(6.01)/Hb: 1.73, ddt(19.0, 10.1, 2.6) |
| 8 | 29.7 | 2.49, m |
| 9 | 130.7 | 5.75, d(6.3) |
| 10 | 137.5 | - |
| 11 | 42.4 | 2.44, m |
| 12 | 32.0 | Ha: 1.94, m/Hb: 1.61, ddt(14.1, 12.5, 4.2) |
| 13 | 17.8 | Ha: 1.24, m/Hb: 1.03, m |
| 14 | 14.9 | 0.89, t(7.3) |
| 15 | 24.2 | 1.19, s |
| 16 | 20.8 | 1.66, s |
| 1’ | 172.0 | - |
| 2’ | 76.0 | - |
| 3’ | 210.6 | - |
| 4’ | 63.8 | 4.05, t(5.9) |
| 5’ | 27.8 | Ha: 2.07, m/Hb: 1.94, m |
| 6’ | 30.3 | 2.43, m |
| 7’ | 176.38 | - |

In MeOD, 600 MHz for $^1$H and 150 MHz for $^{13}$C NMR; Chemical shifts are reported in ppm. All shifts are determined by $^1$H, $^{13}$C, COSY, HSQC, HMBC and NOESY correlations.

**Table S11:** NMR spectroscopic data for **15**

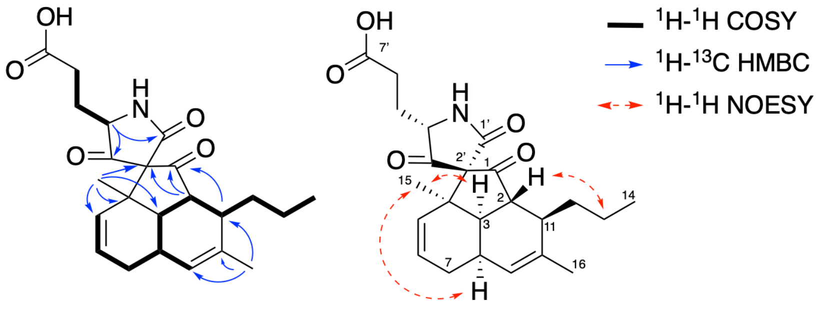

| | Found (MeOD) | |
|---|---|---|
| **Position** | **δC** | **δH, m(J in Hz)** |
| 1 | 210.6 | - |
| 2 | 49.7 | 2.44, m |
| 3 | 46 | 2.37, m |
| 4 | 47.6 | - |
| 5 | 130.6 | 5.44, m |
| 6 | 125.7 | 5.68, m |
| 7 | 28.9 | Ha: 2.36, m; Hb: 1.73, ddt (18.5, 10.0, 2.1) |
| 8 | 29.7 | 2.51, m |
| 9 | 130.6 | 5.75, d(4.6) |
| 10 | 137.4 | - |
| 11 | 42.6 | 2.45, m |
| 12 | 32.1 | Ha:1.92, m/Hb: 1.60, m |
| 13 | 17.9 | Ha: 1.20, m/Hb: 1.04, m |
| 14 | 14.9 | 0.89, t(7.1) |
| 15 | 24.4 | 1.34, br s |
| 16 | 20.8 | 1.66, s |
| 1’ | 173.3 | - |
| 2’ | 76.1 | - |
| 3’ | 208.4 | - |
| 4’ | 63.6 | 3.97, t(7.4) |
| 5’ | 28.4 | Ha: 1.98, m/Hb: 1.84, m |
| 6’ | 31.0 | 2.46, m |
| 7’ | 176.5 | - |

In MeOD, 600 MHz for $^1$H and 150 MHz for $^{13}$C NMR; Chemical shifts are reported in ppm. All shifts are determined by $^1$H, $^{13}$C, COSY, HSQC, HMBC and NOESY correlations.

**Table S12:** NMR spectroscopic data for **16**

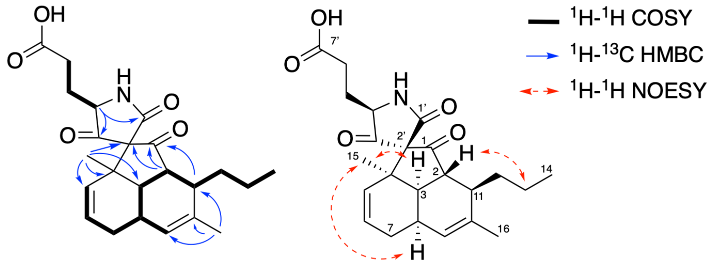

| | Found (MeOD) | |
|---|---|---|
| Position | $\delta_C$ | $\delta_H$, m(J in Hz) |
| 1 | 210.1 | - |
| 2 | 49.6 | 2.59, m |
| 3 | 46.0 | 2.28, m |
| 4 | 47.2 | - |
| 5 | 131.0 | 5.80, d(10.5) |
| 6 | 125.0 | 5.65, m |
| 7 | 28.9 | Ha: 2.36, m/Hb: 1.73 dd( 18.9, 10) |
| 8 | 29.6 | 2.5, m |
| 9 | 130.7 | 5.75, d(5.93) |
| 10 | 137.4 | - |
| 11 | 42.7 | 2.43, m |
| 12 | 32.0 | Ha: 1.93, m/Hb: 1.61, m |
| 13 | 17.8 | Ha: 1.23, dp(12.6, 5.9) /Hb: 1.03, m |
| 14 | 14.9 | 0.89, t(7.1) |
| 15 | 24.7 | 1.19, s |
| 16 | 20.8 | 1.66, s |
| 1’ | 172.0 | - |
| 2’ | 76.2 | - |
| 3’ | 209.6 | - |
| 4’ | 63.2 | 3.97, t(5.9) |
| 5’ | 28.7 | Ha: 2.03, m/Hb: 1.81, dt(14.0, 7.3) |
| 6’ | 31.0 | 2.46, m |
| 7’ | 176.3 | - |

In MeOD, 600 MHz for $^1$H and 150 MHz for $^{13}$C NMR; Chemical shifts are reported in ppm. All shifts are determined by $^1$H, $^{13}$C, COSY, HSQC, HMBC and NOESY correlations.

**Table S13:** NMR spectroscopic data for **17**

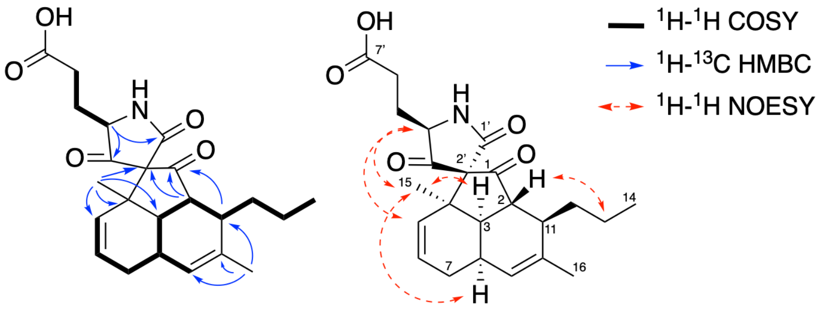

| | Found (MeOD) | |
|---|---|---|
| **Position** | **δC** | **δH, m(J in Hz)** |
| 1 | 211.1 | - |
| 2 | 49.7 | 2.47, m |
| 3 | 45.9 | 2.34, m |
| 4 | 48 | - |
| 5 | 130.5 | 5.44 d(10.5) |
| 6 | 126 | 5.68 ddd(14.9, 10.4, 3.9) |
| 7 | 29 | Ha: 2.35, m/ Hb: 1.76, m |
| 8 | 30.3 | 2.46, m |
| 9 | 130.7 | 5.75, d (5.6) |
| 10 | 137.5 | - |
| 11 | 42.3 | 2.46, m |
| 12 | 32 | Ha: 1.92, m/ Hb: 1.62, m |
| 13 | 17.9 | Ha: 1.22, m/ Hb: 1.05, m |
| 14 | 14.9 | 0.90 t(7.2) |
| 15 | 24.1 | 1.34, s |
| 16 | 20.9 | 1.67, s |
| 1’ | 173.4 | - |
| 2’ | 76.2 | - |
| 3’ | 208.9 | - |
| 4’ | 63.9 | 4.08, t (5.9) |
| 5’ | 27.4 | Ha: 2.07, m/ Hb: 1.95, m |
| 6’ | 30.2 | 2.46 |
| 7’ | 176.4 | - |

In MeOD, 600 MHz for $^1H$ and 150 MHz for $^{13}C$ NMR; Chemical shifts are reported in ppm. All shifts are determined by $^1H$, $^{13}C$, COSY, HSQC, HMBC and NOESY correlations.

**Table S14:** NMR spectroscopic data for **19**

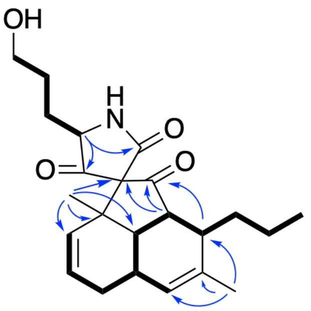

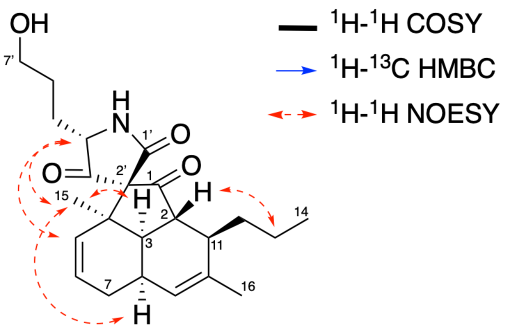

| | Found (MeOD) | |
|---|---|---|
| Position | $\delta_C$ | $\delta_H$, m(J in Hz) |
| 1 | 210.8 | - |
| 2 | 49.4 | 2.51, m |
| 3 | 45.9 | 2.26, dd(14.1, 3.5) |
| 4 | 47.9 | - |
| 5 | 130.8 | 5.72, d(10.7) |
| 6 | 125.5 | 5.65, m |
| 7 | 29.1 | Ha: 2.36, m/ Hb: 1.74, m |
| 8 | 29.8 | 2.49, m |
| 9 | 130.7 | 5.75, d(5.1) |
| 10 | 137.5 | - |
| 11 | 42.5 | 2.44, m |
| 12 | 32.0 | Ha: 1.94, m/ Hb: 1.59, m |
| 13 | 17.8 | Ha: 1.24, m/ Hb:1.00, d(8.3) |
| 14 | 14.9 | 0.89, t(7.1) |
| 15 | 24.13 | 1.20, s |
| 16 | 20.7 | 1.66, s |
| 1’ | 172.9 | - |
| 2’ | 76.1 | - |
| 3’ | 211.0 | - |
| 4’ | 64.6 | 4.00, t(5.9) |
| 5’ | 29.2 | Ha: 1.87, m/ Hb: 1.72, m |
| 6’ | 29.0 | Ha: 1.65, m/ Hb: 1.59, m |
| 7’ | 62.4 | 3.57, t(6.1) |

In MeOD, 600 MHz for $^1$H and 150 MHz for $^{13}$C NMR; Chemical shifts are reported in ppm. All shifts are determined by $^1$H, $^{13}$C, COSY, HSQC, HMBC and NOESY correlations.

### Supporting Figures

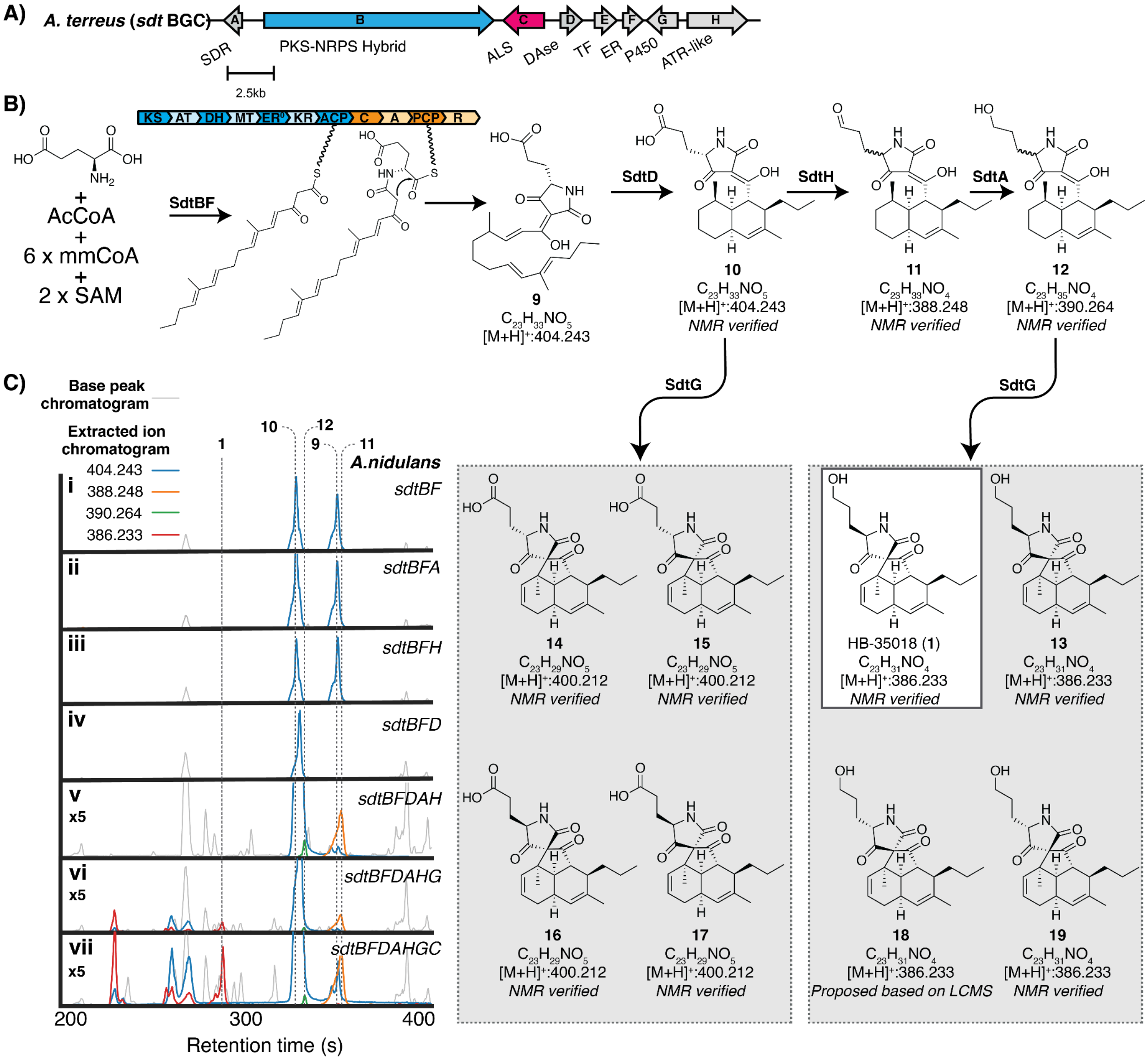

**Figure S1:** Detailed proposed biosynthesis of **1** inclusive of all isolated shunts and intermediates. Abbreviations: short chain reductase reductase (SDR), polyketide synthase/non-ribosomal synthetase hybrid (PKS/NRPS hybrid), Diels-Alderase (DAse), transcription factor (TF), enoyl-reductase (ER), cytochrome P450 (P450), adenylation-thiolation-reductase-like (ATR-like), ketosynthase (KS), acyltransferase (AT), dehydratase (DH), methyltransferase (MT), ketoreductase (KR), acyl carrier protein (ACP), condensation (C), adenylation (A), peptidyl carrier protein (PCP), reductase (R).

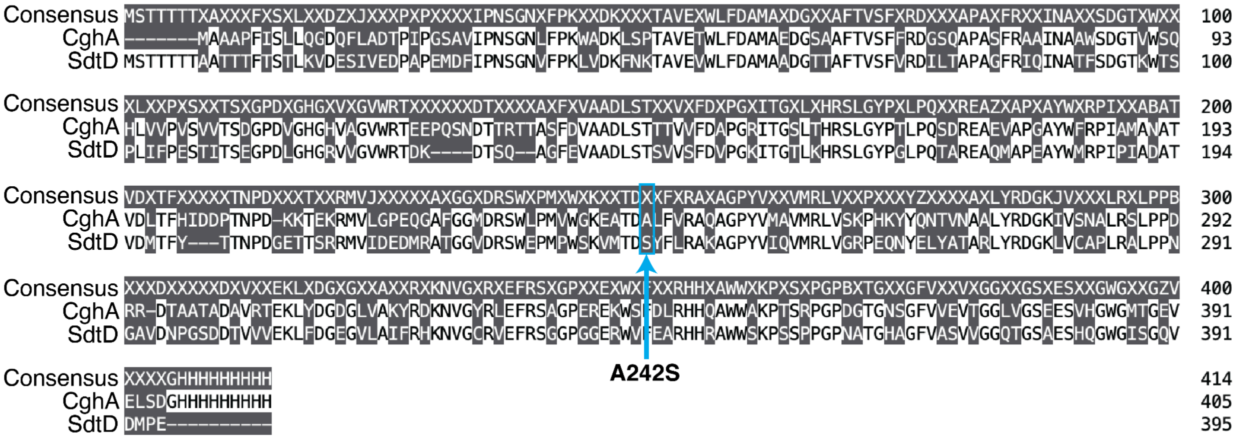

**Figure S2:** Sequence alignment of CghA and SdtD highlighting the A to S difference at position 242.

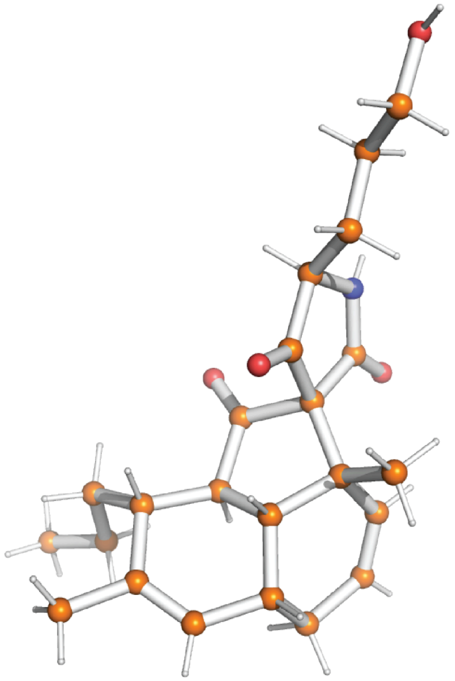

| | |
|---|---|
| Crystal system | orthorhombic |
| Space group | $P2_12_12_1$ |
| *a* [Å] | 7.6508 ± 0.0005 |
| *b* [Å] | 9.8384 ± 0.0006 |
| *c* [Å] | 28.002 ± 0.002 |
| *α* [º] | 90 |
| *β* [º] | 90 |
| *γ* [º] | 90 |

**Figure S3:** Small molecule X-ray structure of **1.**

.

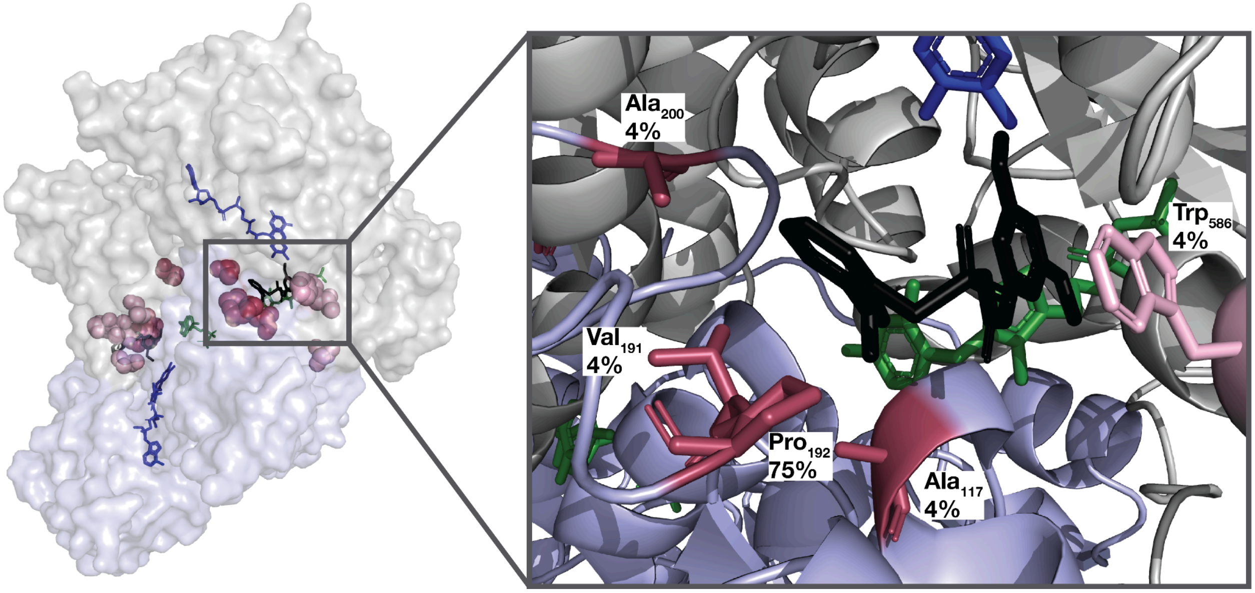

**Figure S5:** Mutations in ALS observed in *S. cerevisiae* clones resistant to **1**. Resistance mutations (shown in pink and raspberry) are mapped onto the crystal structure of **2** (black) bound to S. cerevisiae ALS (PDB:5FEM30, α-chain in gray, β-chain in blue). Essential thiamine pyrophosphate (TPP,green) and FAD (blue) cofactors shown. All mutations are annotated with the percentage of clones in which the residue was mutated.

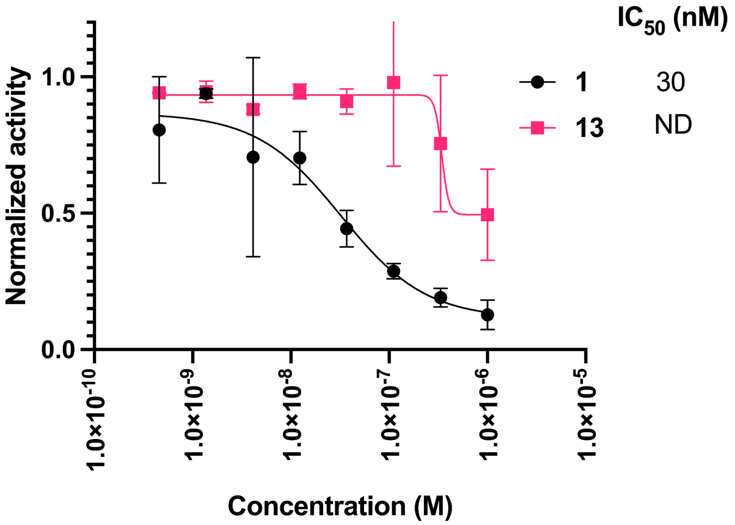

**Figure S4:** Inhibition of recombinant *S. cerevisiae* ALS by **1** and its C12 diastereomer **13**

A)

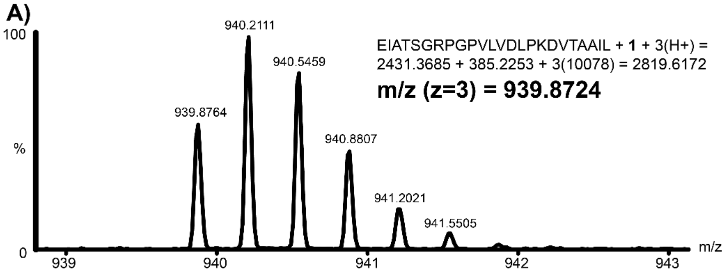

B)

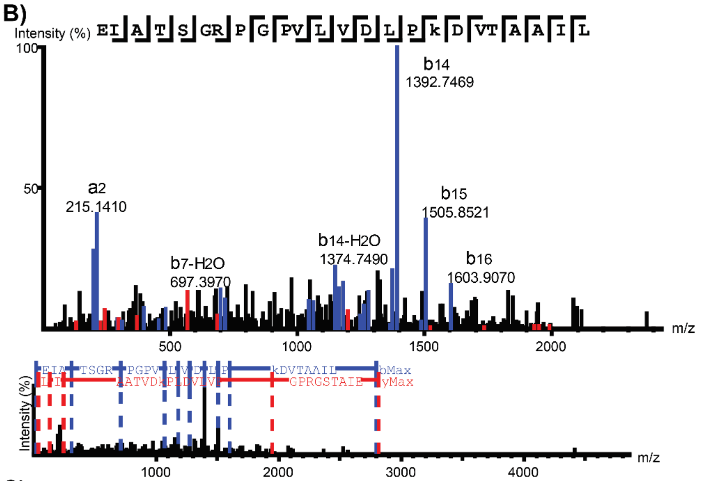

C)

| # | b | b-H2O | a | a-NH3 | Seq | y | y-H2O | z | # |
|---|---|---|---|---|---|---|---|---|---|
| 1 | 130.0504 | 112.0399 | 102.0554 | 85.0284 | E | | | | 24 |
| 2 | 243.1345 | 225.1239 | 215.1410 | 198.1230 | I | 2688.5586 | 2670.5481 | 2671.5315 | 23 |
| 3 | 314.1680 | 296.1620 | 286.1766 | 269.1496 | A | 2575.4744 | 2557.4639 | 2558.4473 | 22 |
| 4 | 415.2192 | 397.2040 | 387.2242 | 370.1973 | T | 2504.4373 | 2486.4268 | 2487.4102 | 21 |
| 5 | 502.2513 | 484.2560 | 474.2563 | 457.3000 | S | 2403.3896 | 2385.3792 | 2386.3625 | 20 |
| 6 | 559.2728 | 541.2622 | 531.2778 | 514.2508 | G | 2316.3577 | 2298.3472 | 2299.3306 | 19 |
| 7 | 715.4030 | 697.3970 | 687.3788 | 670.3519 | R | 2259.3362 | 2241.3257 | 2242.3091 | 18 |
| 8 | 812.4266 | 794.4161 | 784.4316 | 767.4047 | P | 2103.2351 | 2085.2246 | 2086.2080 | 17 |
| 9 | 869.4481 | 851.4375 | 841.4531 | 824.4261 | G | 2006.1824 | 1988.0790 | 1989.1554 | 16 |
| 10 | 966.5009 | 948.4903 | 938.5059 | 921.4789 | P | 1949.0940 | 1931.1504 | 1932.1541 | 15 |
| 11 | 1065.5601 | 1047.5861 | 1037.5742 | 1020.5472 | V | 1852.1082 | 1834.0977 | 1835.0812 | 14 |
| 12 | 1178.6530 | 1160.6429 | 1150.6160 | 1133.6313 | L | 1753.0397 | 1734.9520 | 1736.0127 | 13 |
| 13 | 1277.7070 | 1259.7040 | 1249.7271 | 1232.6997 | V | 1639.9556 | 1621.9451 | 1622.9286 | 12 |
| 14 | 1392.7469 | 1374.7490 | 1364.7679 | 1347.7267 | D | 1540.8872 | 1522.8820 | 1523.8602 | 11 |
| 15 | 1505.8521 | 1487.8690 | 1477.8378 | 1460.8108 | L | 1425.8602 | 1407.8497 | 1408.8333 | 10 |
| 16 | 1602.9041 | 1584.8750 | 1574.8905 | 1557.8635 | P | 1312.7762 | 1294.7657 | 1295.7493 | 9 |
| 17 | 2116.2058 | 2098.1953 | 2088.2109 | 2071.1838 | K(+385.23) | 1215.7234 | 1197.6650 | 1198.6964 | 8 |
| 18 | 2231.2327 | 2213.2222 | 2203.2378 | 2186.2107 | D | 702.4032 | 684.3800 | 685.3762 | 7 |
| 19 | 2330.3010 | 2312.2905 | 2302.3062 | 2285.2791 | V | 587.3763 | 569.3460 | 570.3493 | 6 |
| 20 | 2431.3486 | 2413.3381 | 2403.3538 | 2386.3267 | T | 488.3079 | 470.2973 | 471.2809 | 5 |
| 21 | 2502.3857 | 2484.3752 | 2474.3909 | 2457.3638 | A | 387.2602 | 369.1800 | 370.2332 | 4 |
| 22 | 2573.4229 | 2555.4124 | 2545.4280 | 2528.4009 | A | 316.2231 | 298.1430 | 299.1961 | 3 |
| 23 | 2686.5071 | 2668.4966 | 2658.5122 | 2641.4851 | I | 245.1880 | 227.1820 | 228.1590 | 2 |
| 24 | | | | | L | 132.1030 | 114.0913 | 115.0749 | 1 |

D)

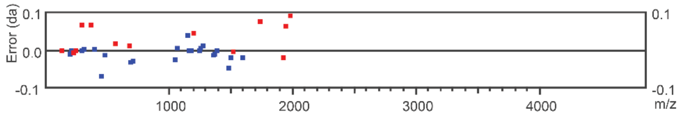

**Figure S6:** Peptide mapping of *S. cerevisiae* ALS treated with **1** localizes the site of covalent modification to the SGRPGPVLVDLPKDVTAAIL peptide. **A)** MS spectrum of the target peptide. **B)** Annotated MS2 spectrum of the target peptide. **C)** Peptide table from $MS^2$ analysis of the target peptide. **D)** Error map of the $MS^2$ analysis.

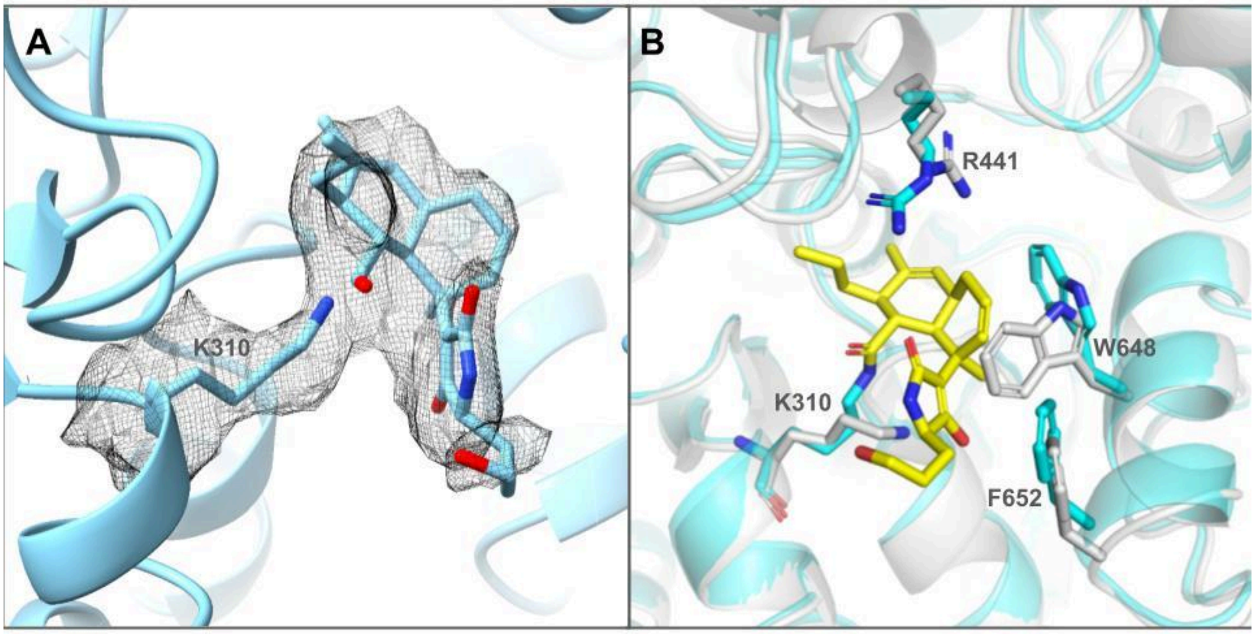

**Figure S7:** cryoEM density of compound **1** and the conformational changes it causes at the Af ALS substrate channel. **A)** The continuous cryoEM density between Af ALS K310 and compound 1 indicates a covalent bond formed between them. **B)** the superposition of Apo (gray) and compound **1** (yellow)-bound (cyan) Af ALS structures.

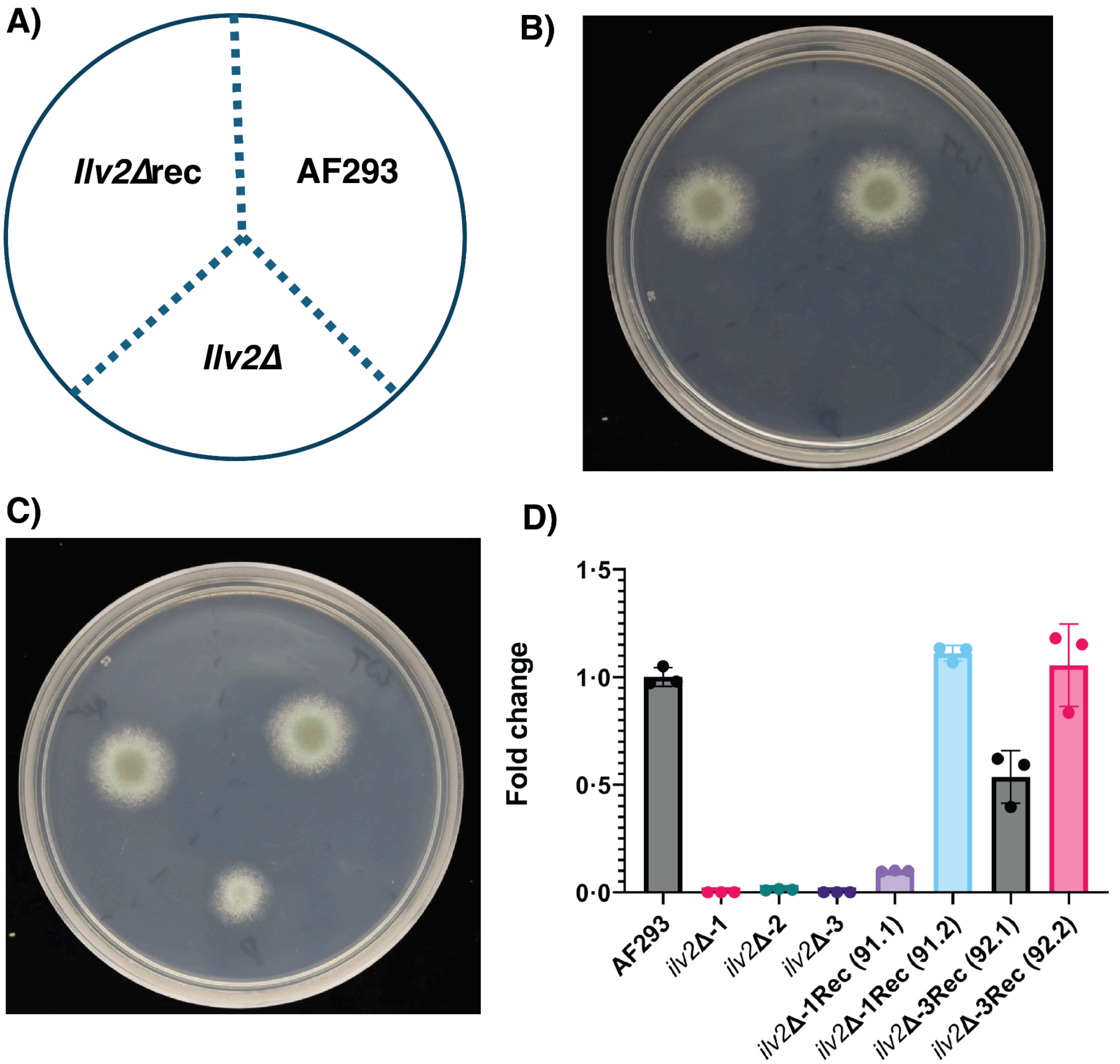

**Figure S8: Validation of *A. fumigatus* mutants A)** Plate map for growth plating experiments **B)** Growth of *A. fumigatus* strains on glucose minimal medium with no supplemented amino acids. **C)** Growth of strains on glucose minimal medium supplemented with 5 mM each of valine and isovaline. All images taken 48 hrs after plating of 1000 spores per strain **D)** RT-qPCR validation of *ilv2Δ* and *ilv2Δ*Rec strains. Clones *ilv2Δ-1* and *ilv2Δ*Rec (91.2) were taken forward for studies in mice.

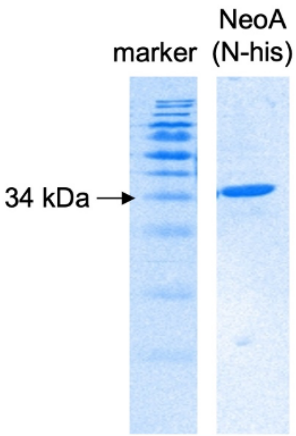

**Figure S9:** SDS page gel of recombinant NeoA

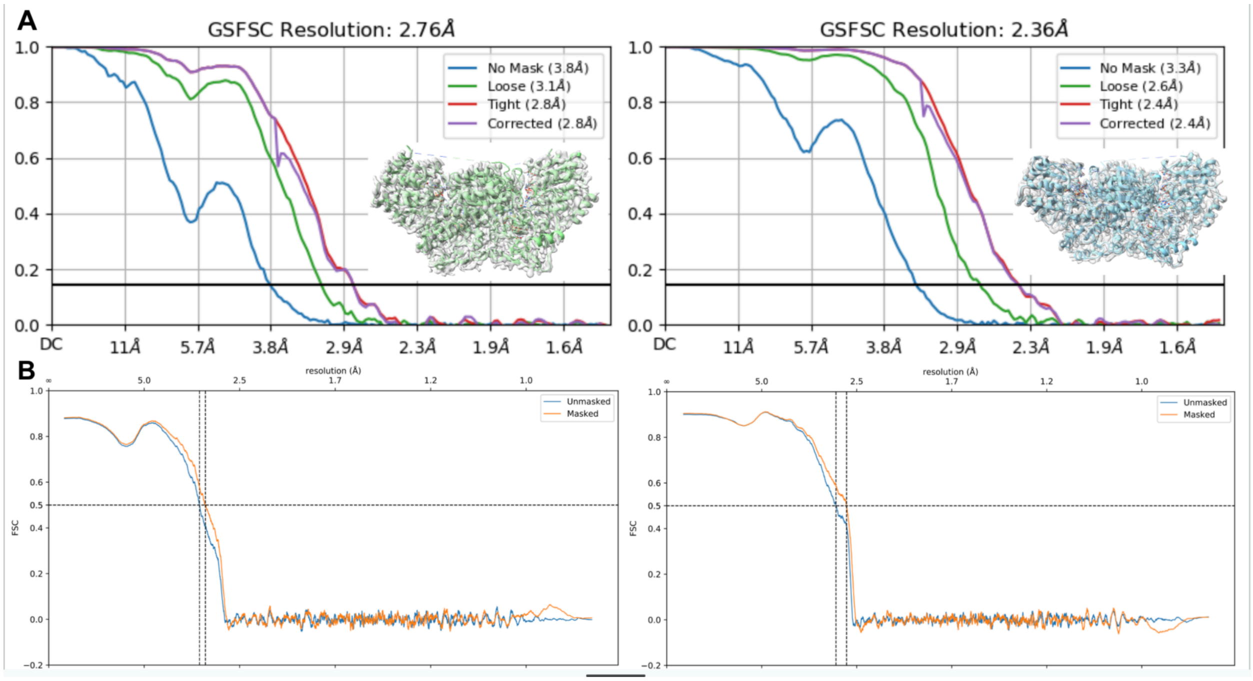

**Figure S10: Quality of the cryo-EM reconstruction of the Apo and 1-bound *Af*ALS. A)** The Fourier shell correlation (FSC) curves of the cryo-EM reconstructions. An overview of the model docked cryoEM map was shown as an inset. Left, Apo *Af*ALS; Right, **1**-bound *Af*ALS. **B)** The model-to-map FSC curves of Apo *Af*ALS (left) and **1**-bound AfALS (right) structures.

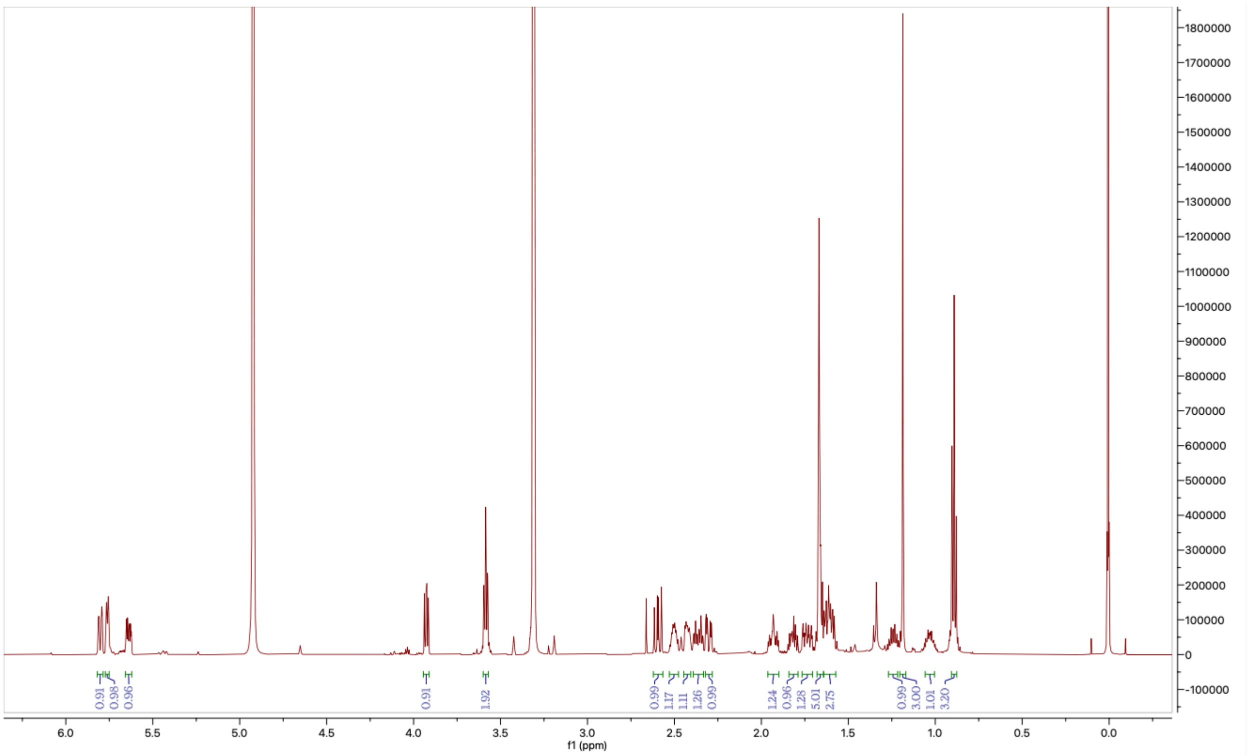

**Figure S11:** The ¹H NMR Spectrum of **1** in MeOD (600 MHz)

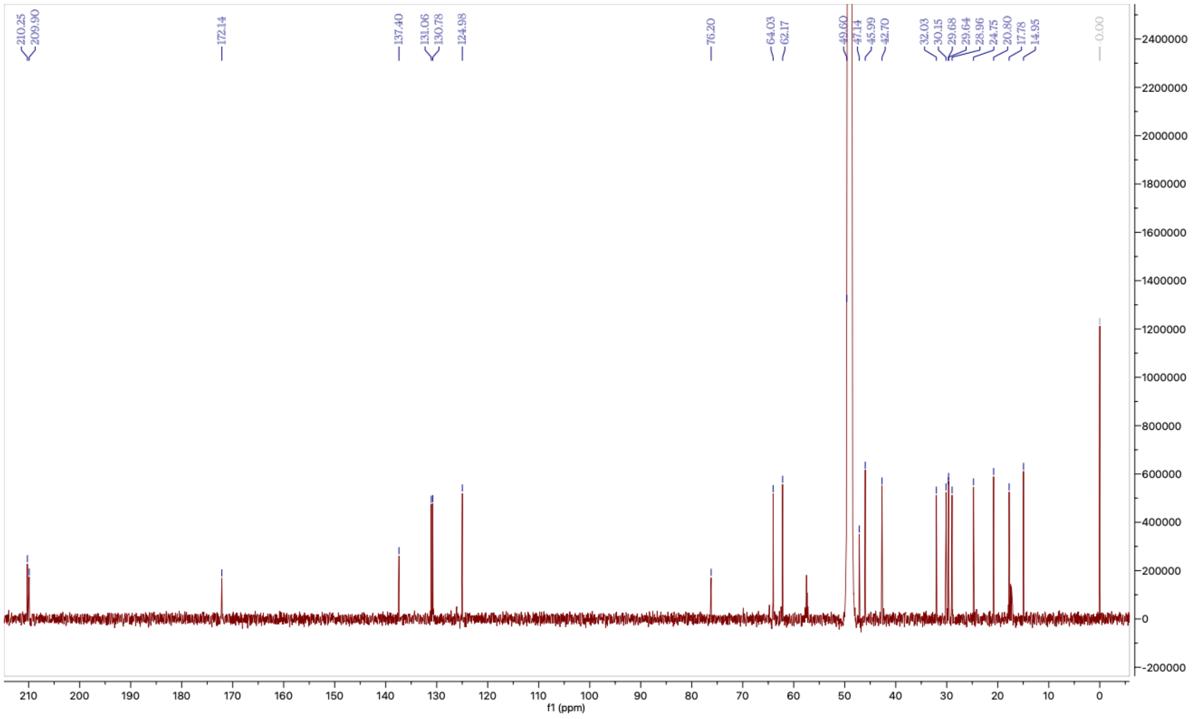

**Figure S12: The** ¹³C NMR Spectrum of **1** in MeOD (150 MHz)

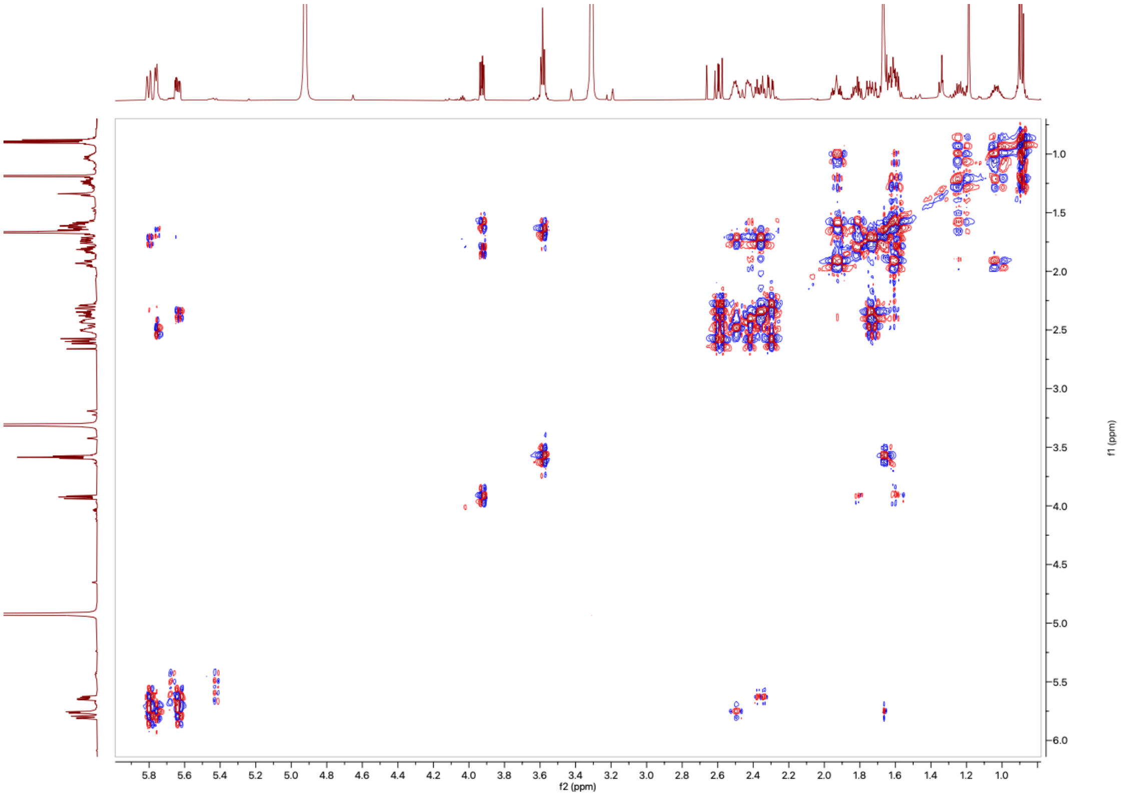

**Figure S13:** The $^1$H-$^1$H COSY NMR Spectrum of **1** in MeOD (600 MHz)

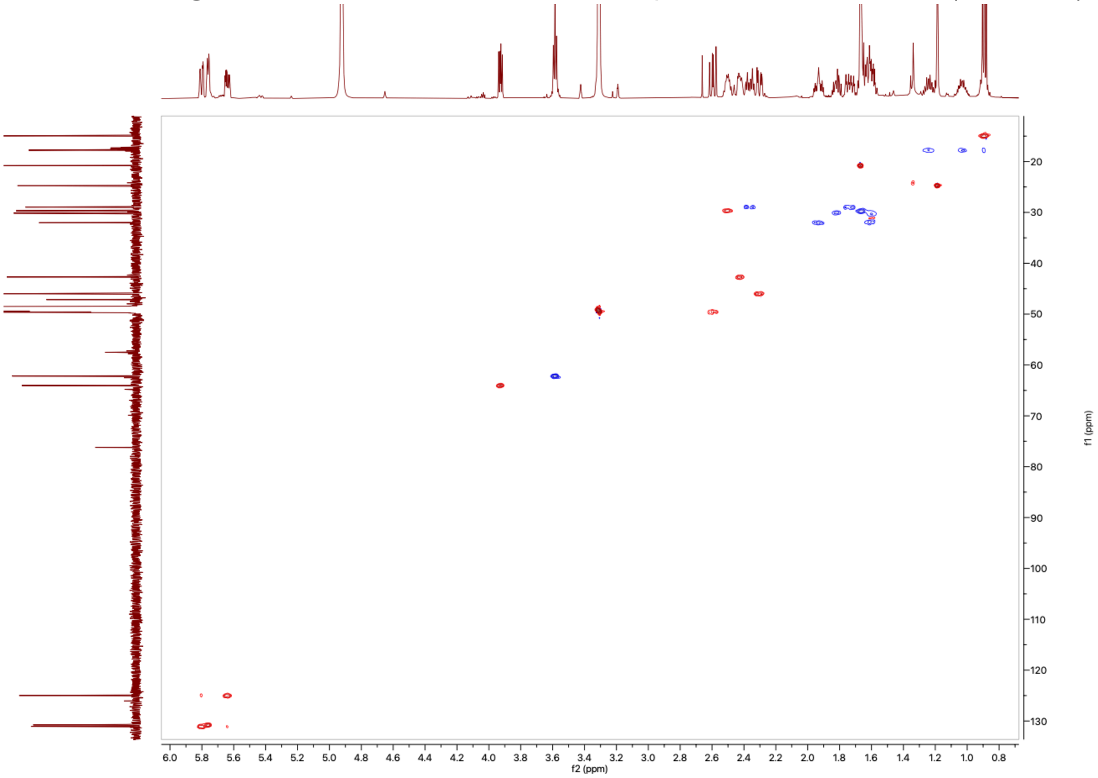

**Figure S14:** The $^1$H-$^{13}$C HSQC NMR Spectrum of **1** in MeOD (600 MHz)

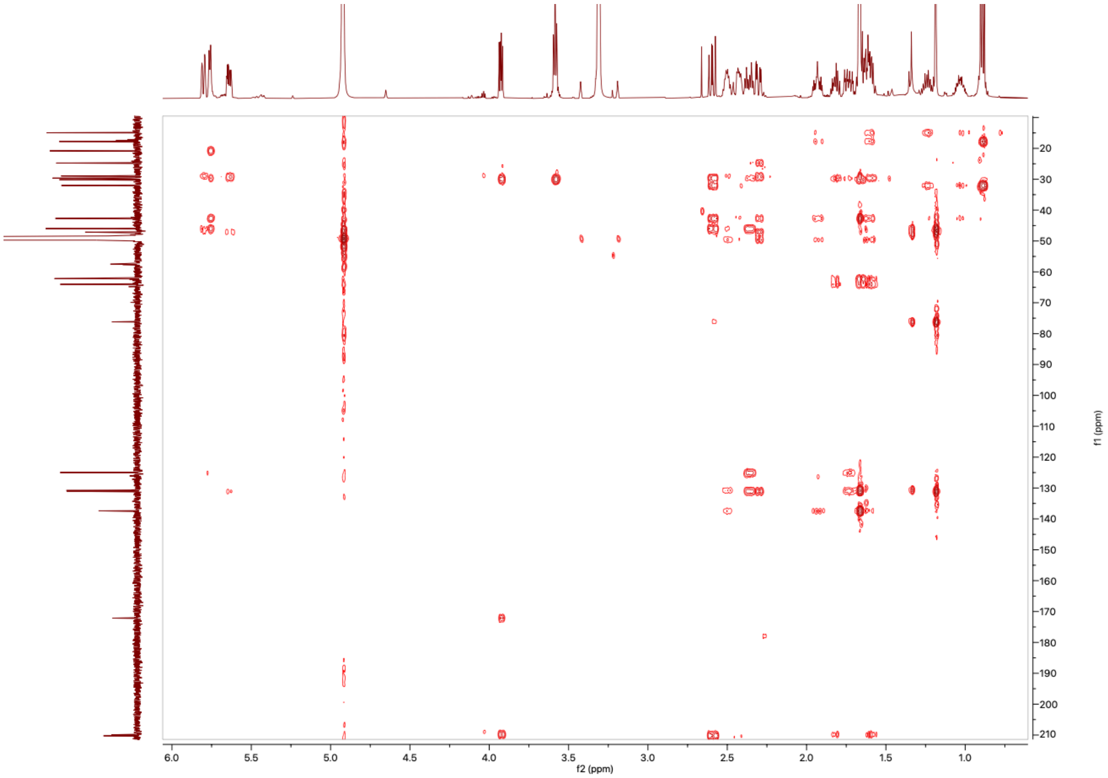

**Figure S15:** The $^{1}H$-$^{13}C$ HMBC NMR Spectrum of **1** in MeOD (600 MHz)

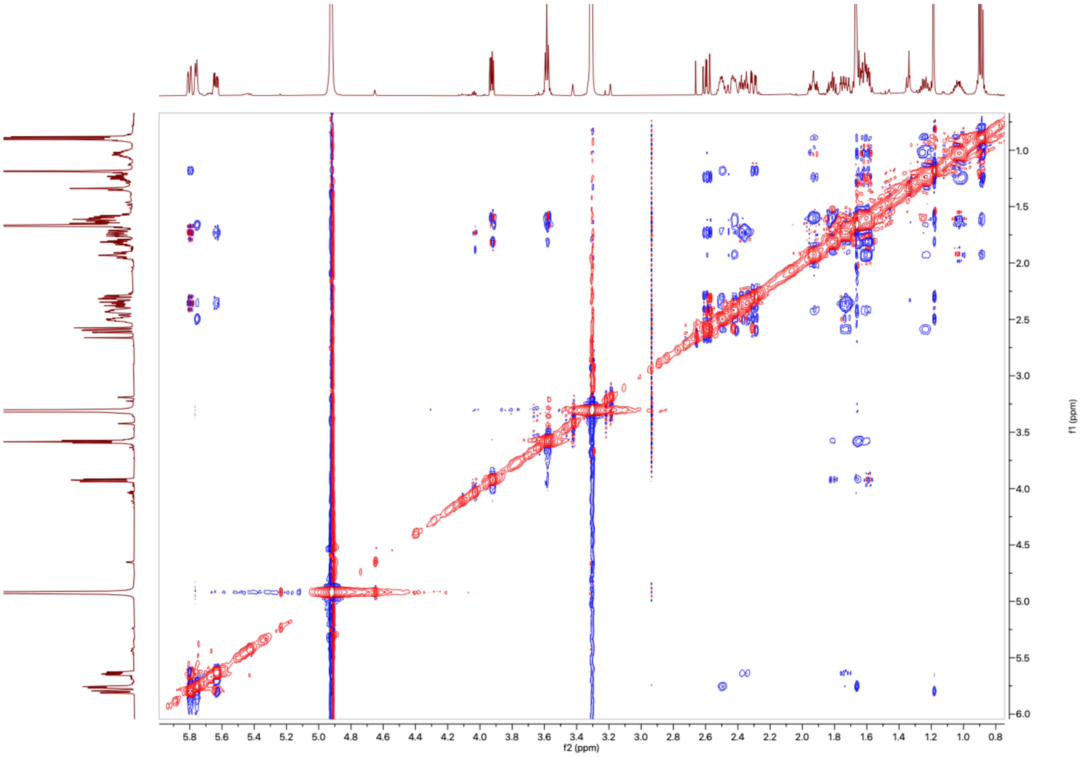

**Figure S16:** The $^{1}H$-$^{1}H$ NOESY NMR Spectrum of **1** in MeOD (600 MHz)

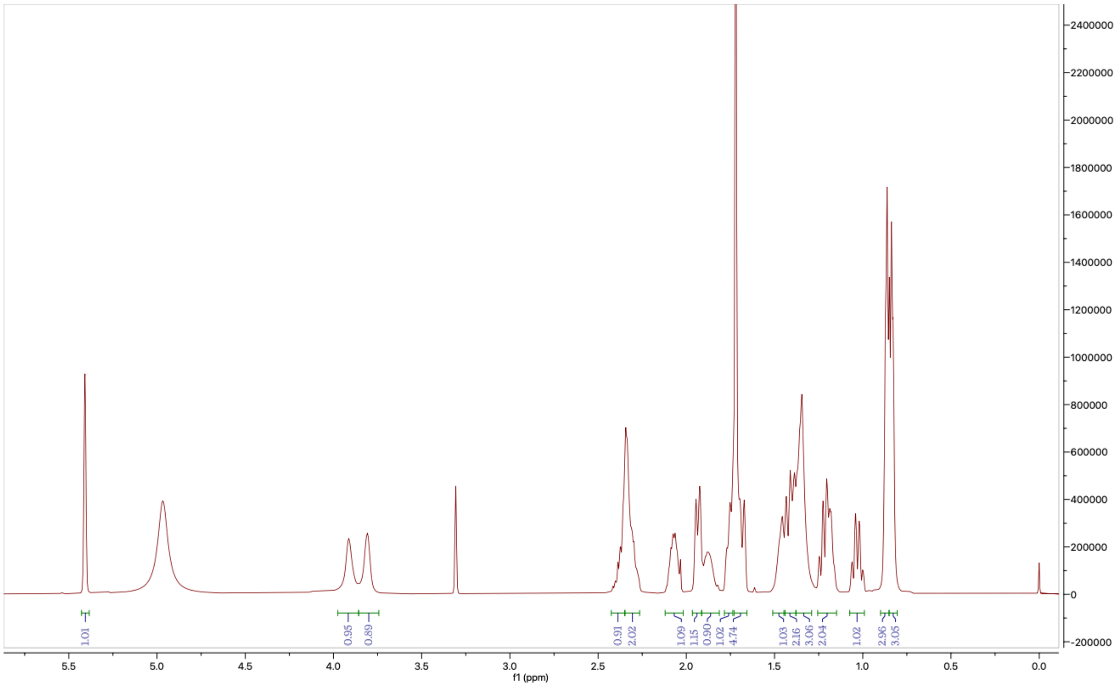

**Figure S17:** The $^{1}$H NMR Spectrum of **10** in MeOD (600 MHz)

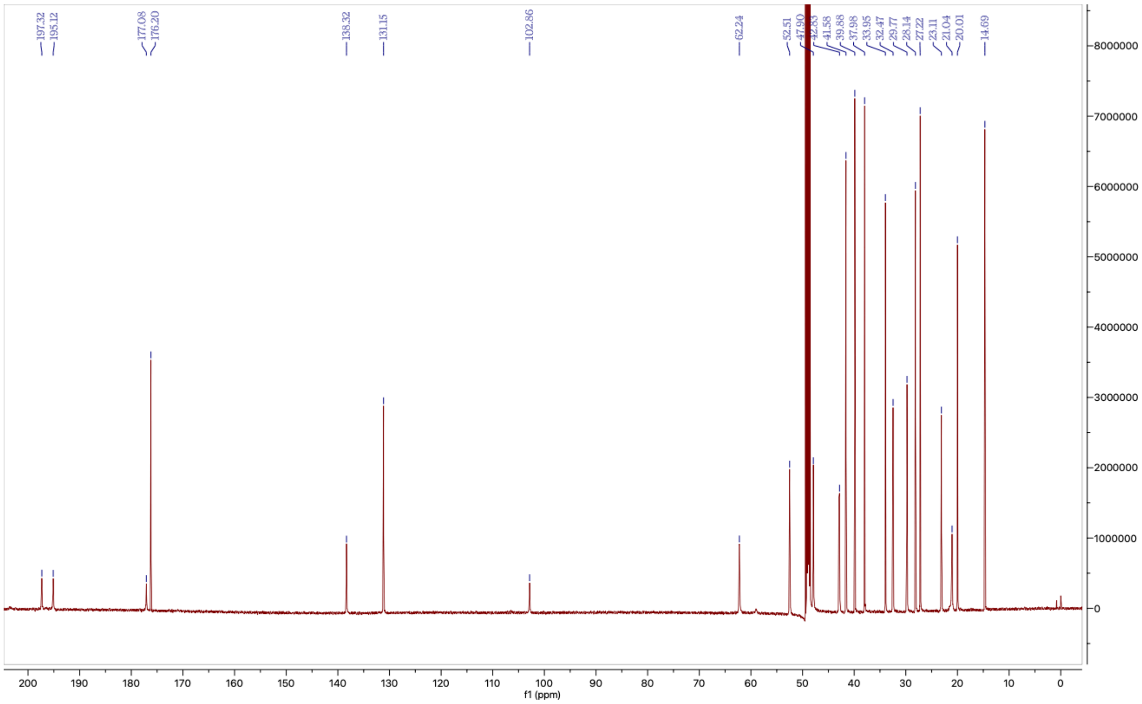

**Figure S18:** The $^{13}$C NMR Spectrum of **10** in MeOD (600 MHz)

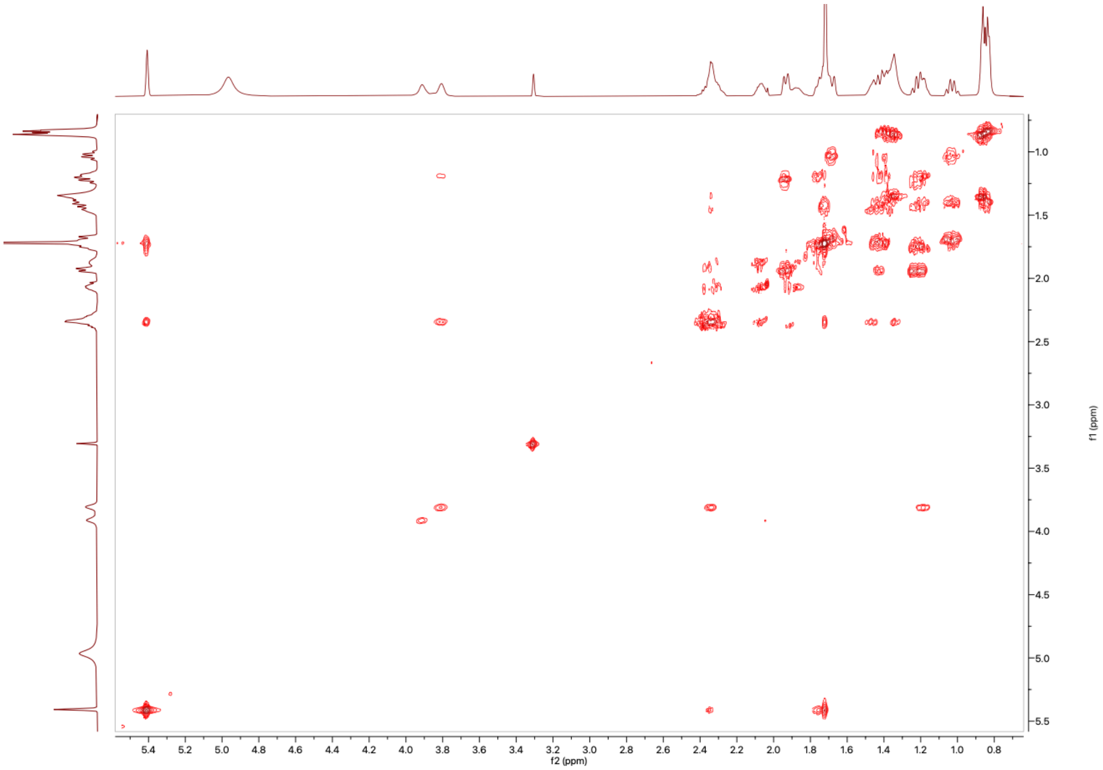

**Figure S19:** The $^1$H-$^1$H COSY NMR Spectrum of **10** in MeOD (600 MHz)

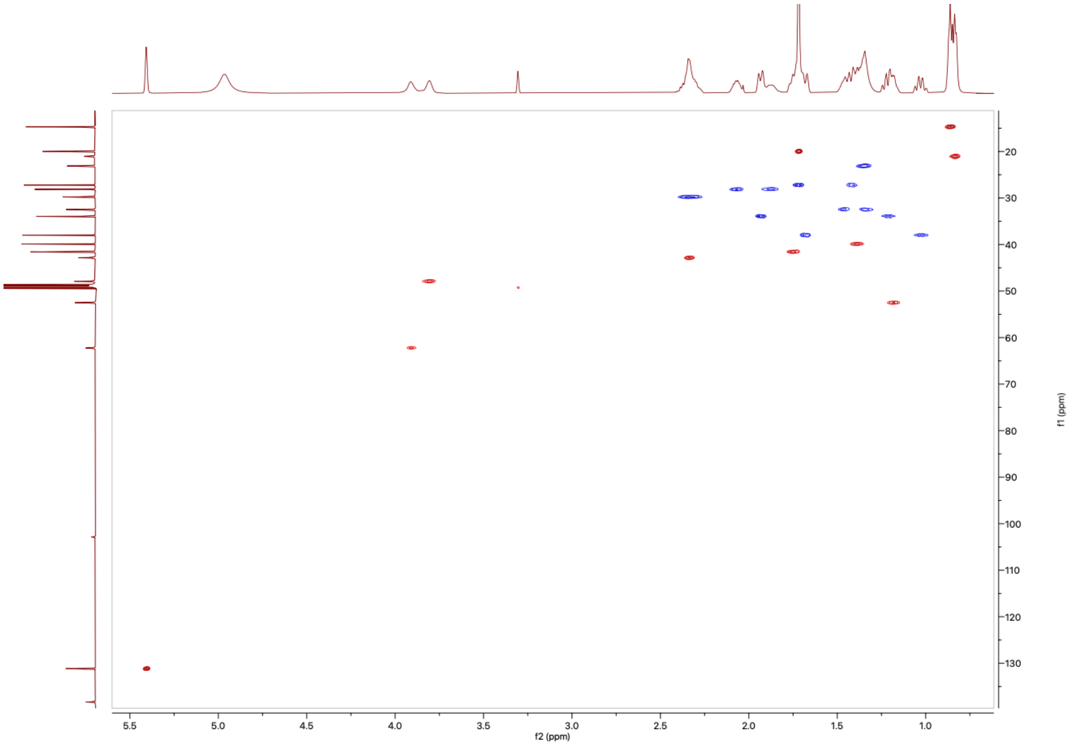

**Figure S20:** The $^1$H-$^{13}$C HSQC NMR Spectrum of **10** in MeOD (600 MHz)

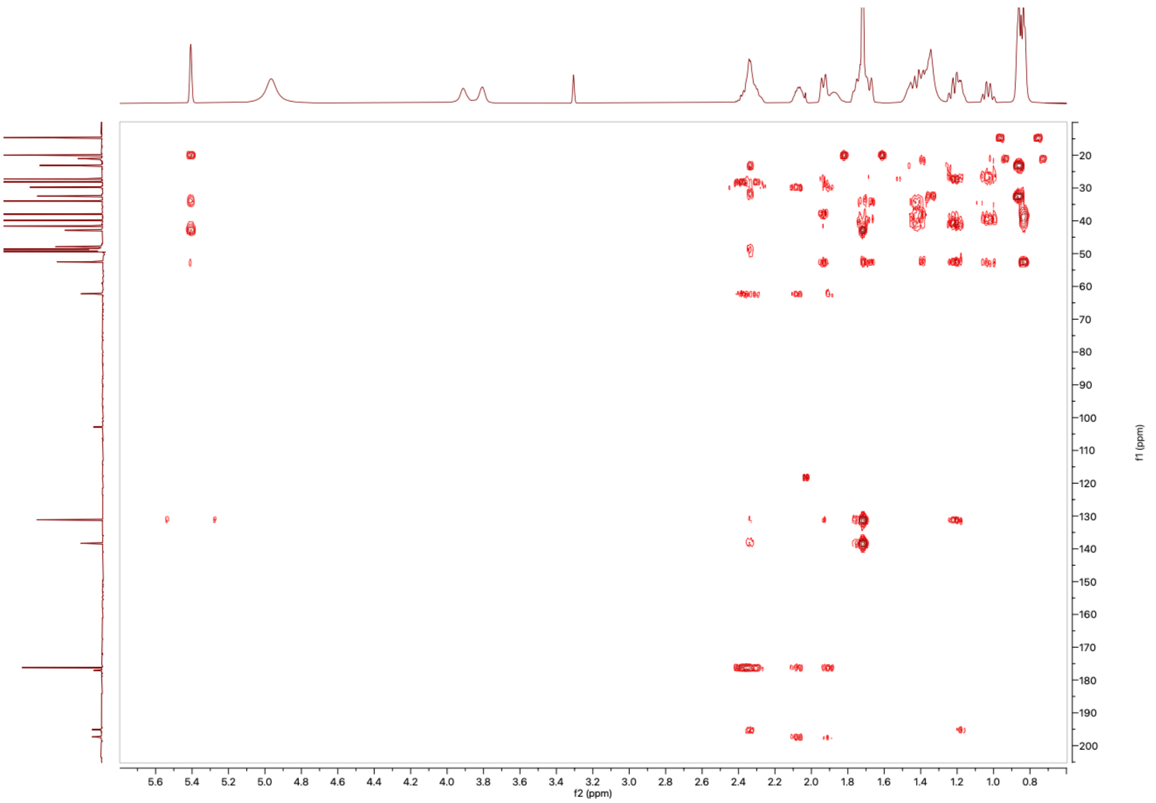

**Figure S21:** The $^1$H-$^{13}$C HMBC NMR Spectrum of **10** in MeOD (600 MHz)

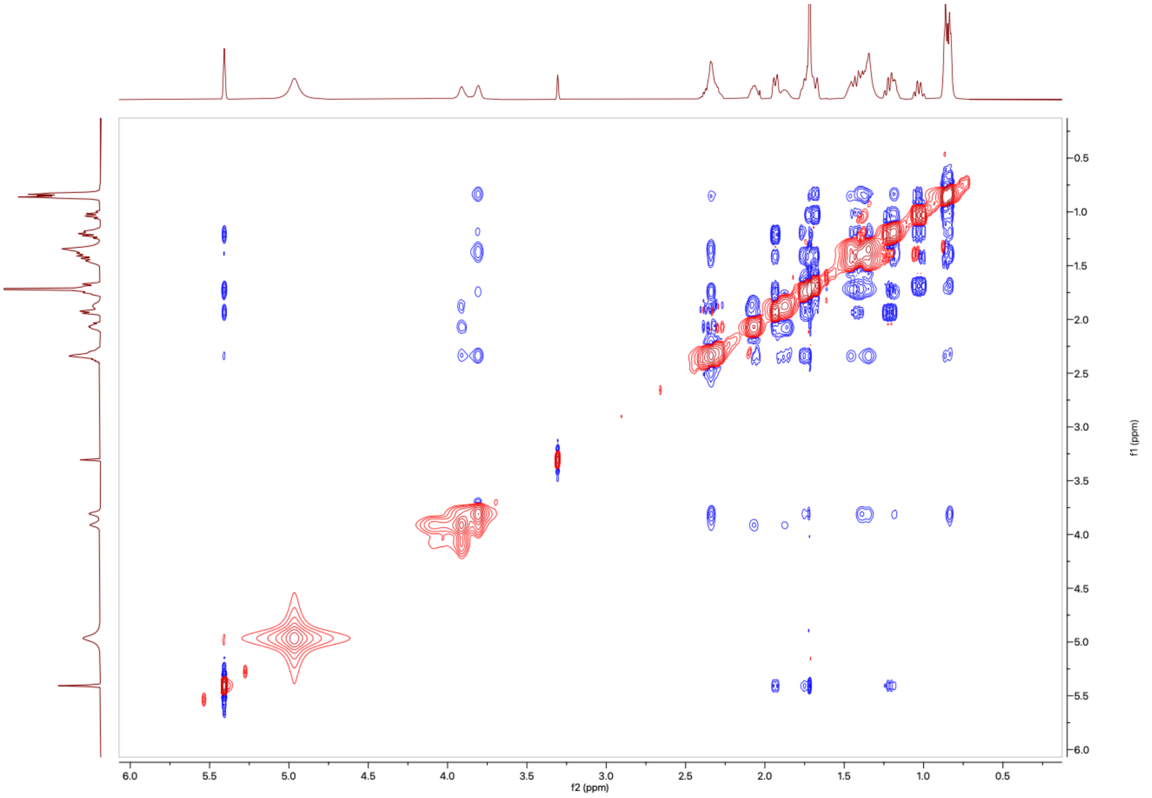

**Figure S22:** The $^1$H-$^1$H NOESY NMR Spectrum of **10** in MeOD (600 MHz)

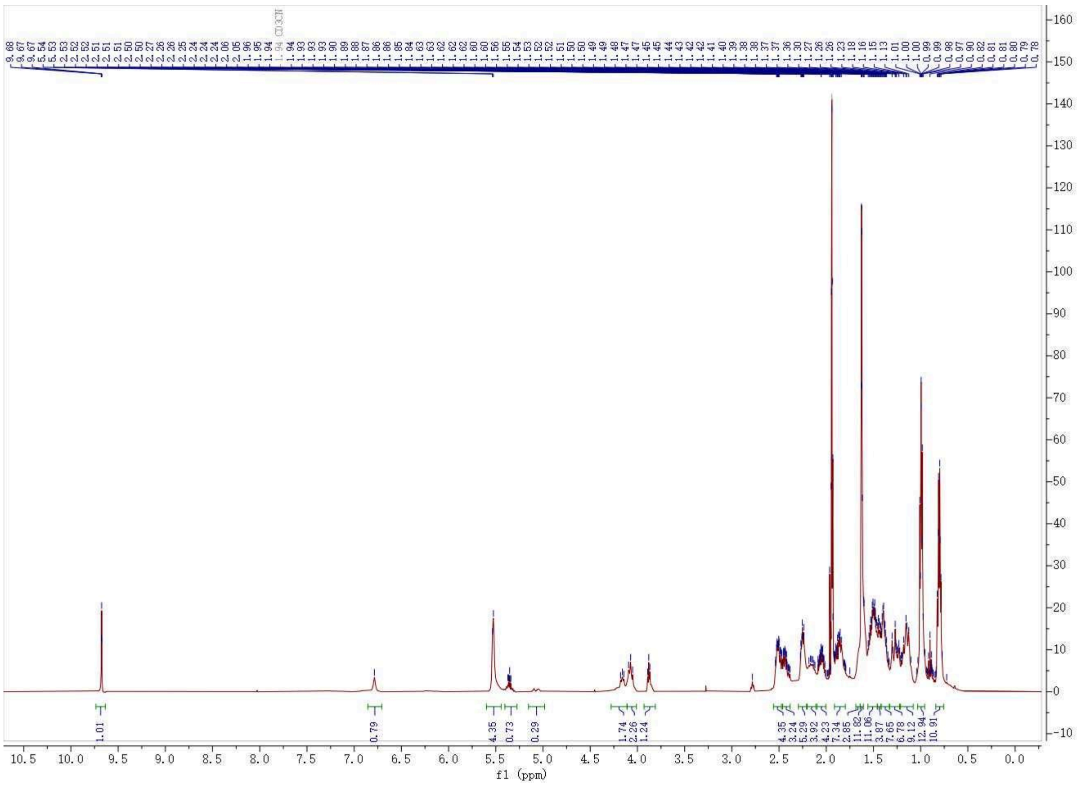

**Figure S23:** The [1]H NMR Spectrum of **11** in $CDCN_3$ (500 MHz)

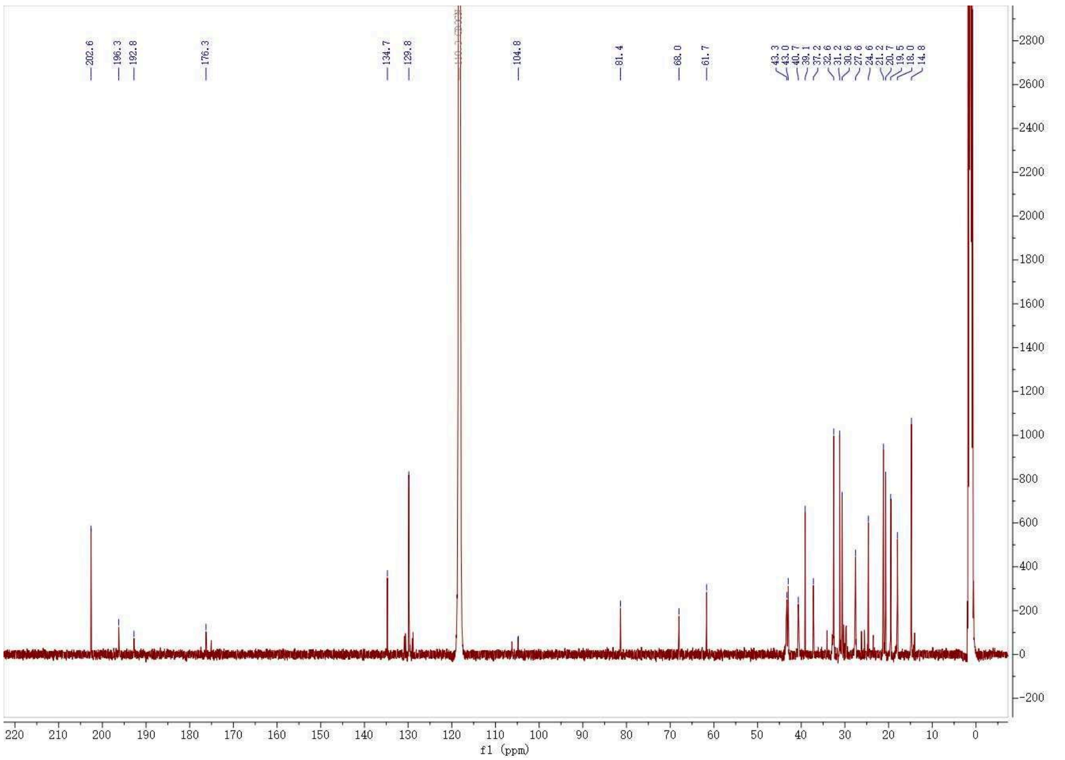

**Figure S24:** The [13]C NMR Spectrum of **11** in $CDCN_3$ (500 MHz)

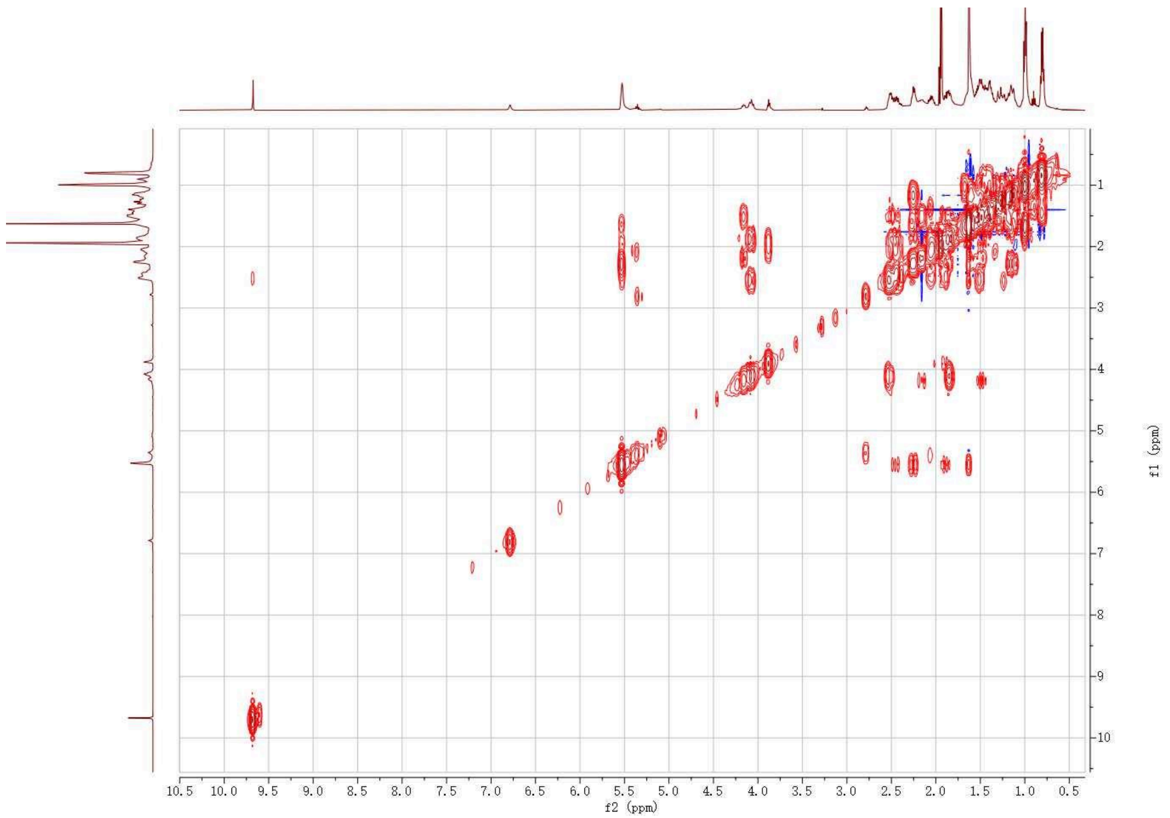

**Figure S25:** The ¹H-¹H COSY NMR Spectrum of **11** in $CDCN_3$ (500 MHz)

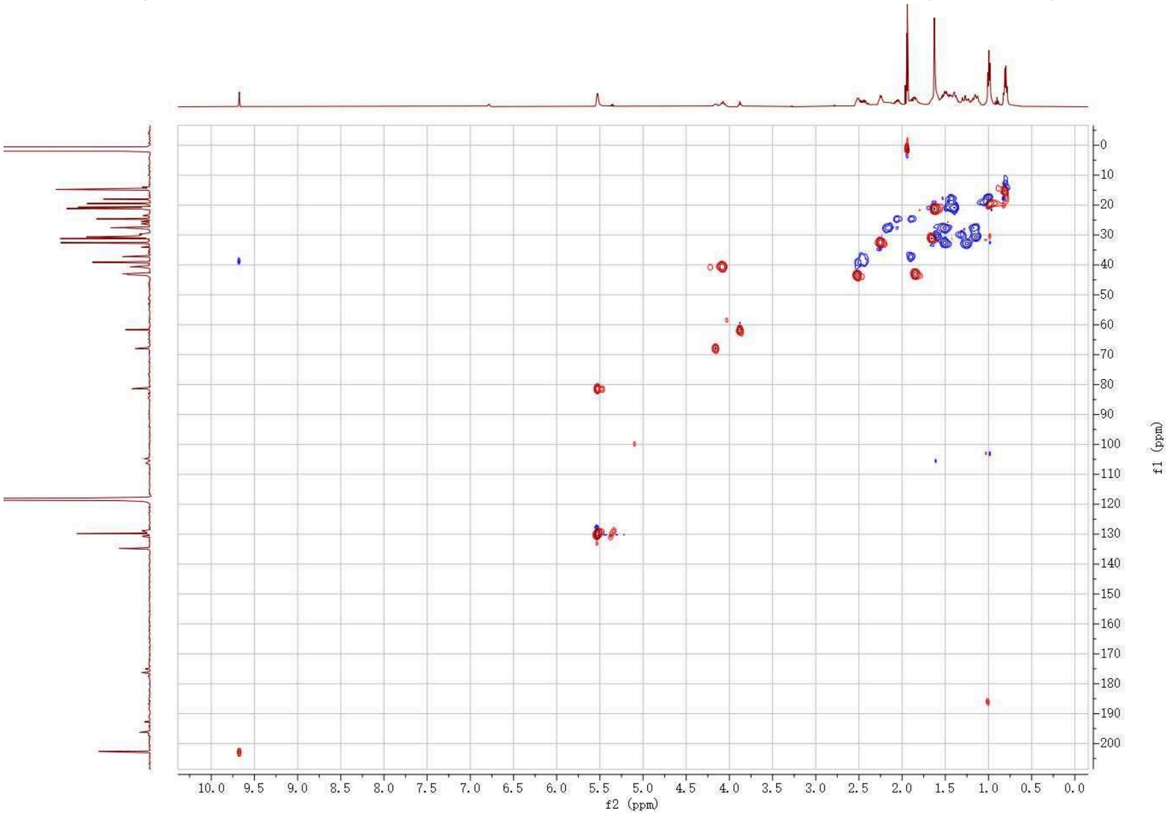

**Figure S26:** The ¹H-¹³C HSQC NMR Spectrum of **11** in $CDCN_3$ (500 MHz)

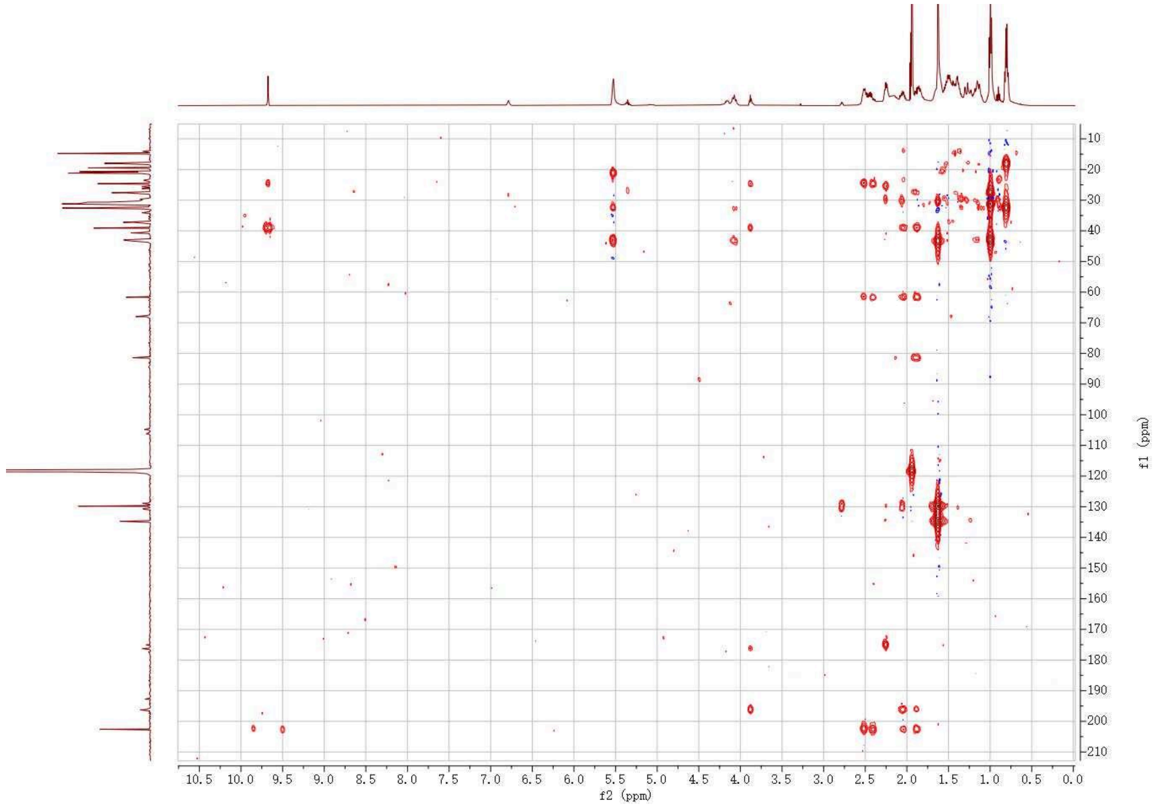

**Figure S27:** The $^{1}H$-$^{13}C$ HMBC NMR Spectrum of **11** in $CDCN_3$ (500 MHz)

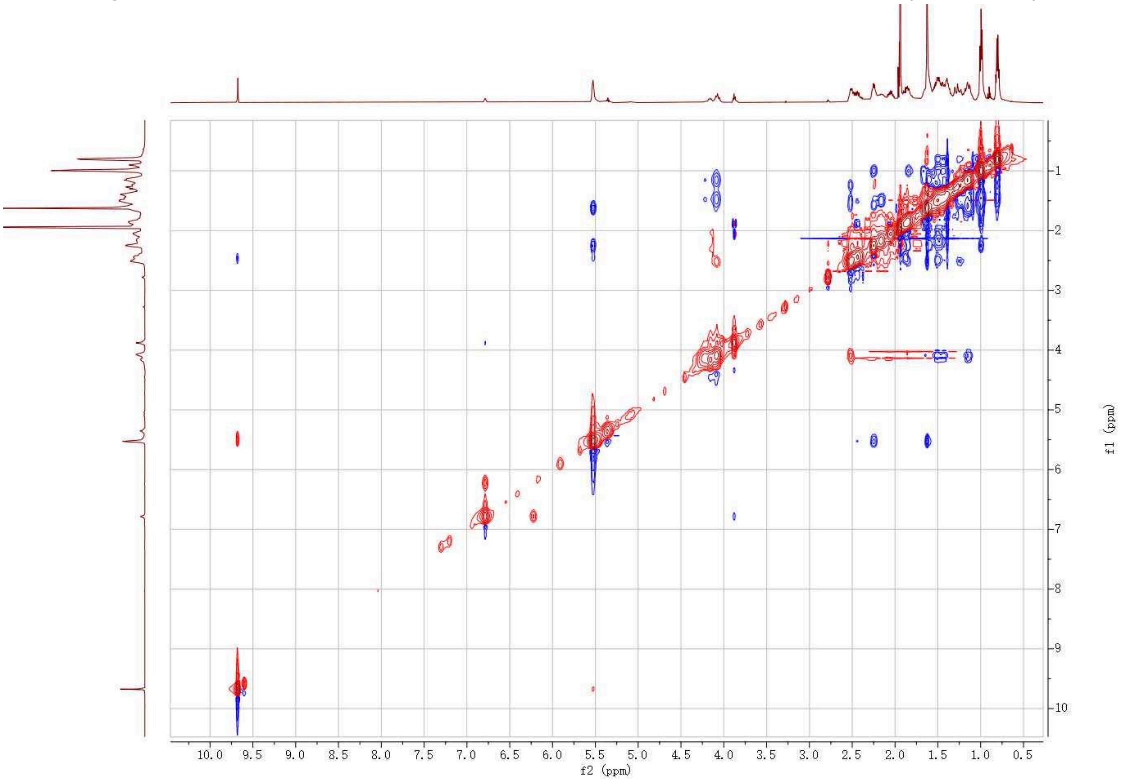

**Figure S28:** The $^{1}H$-$^{1}H$ NOESY NMR Spectrum of **11** in $CDCN_3$ (500 MHz)

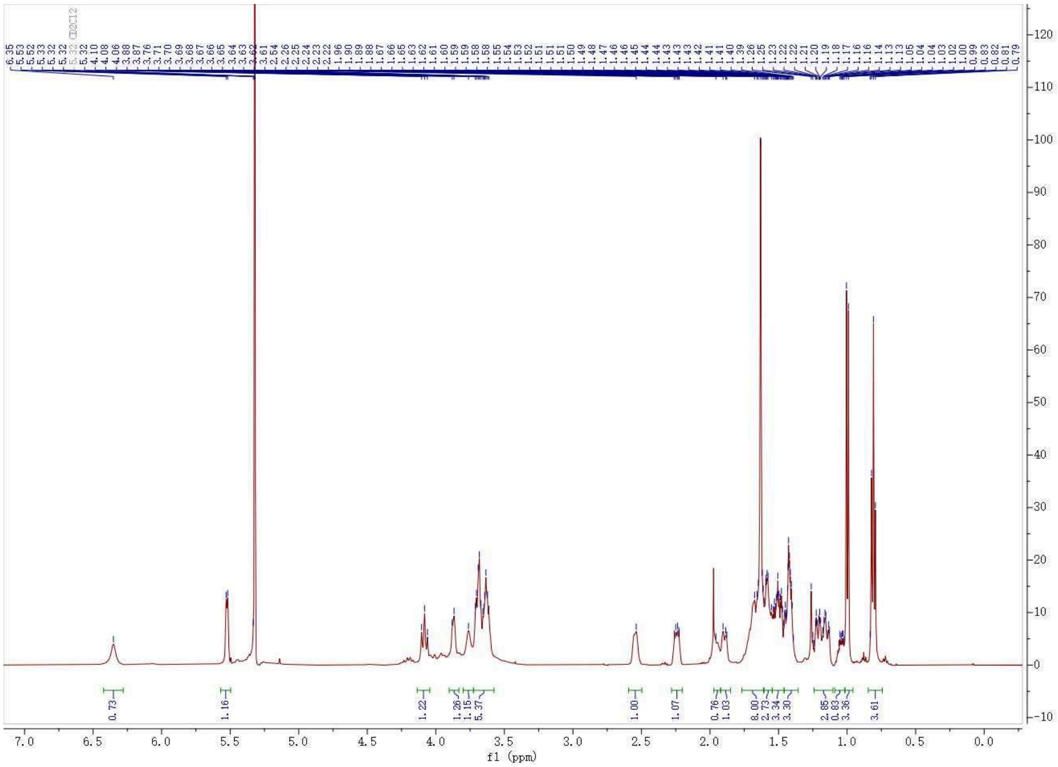

**Figure S29:** The ¹H NMR Spectrum of **12** in $CD_2Cl_2$ (500 MHz)

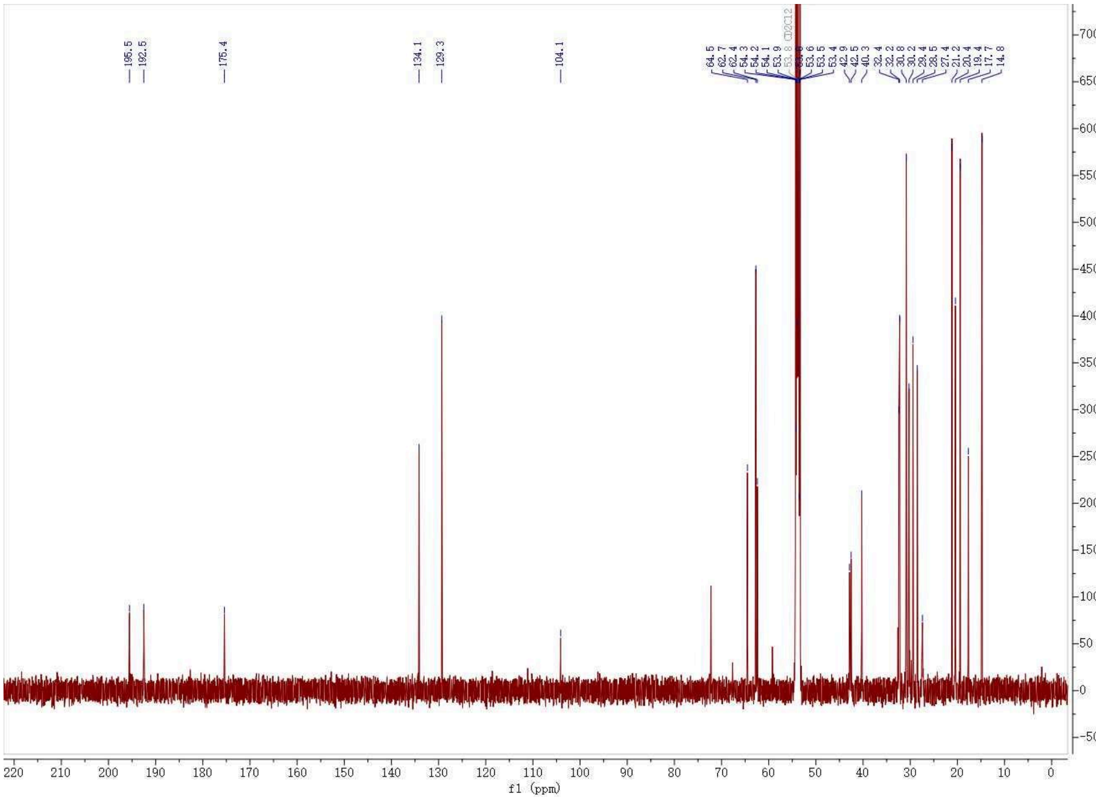

**Figure S30:** The ¹³C NMR Spectrum of **12** in $CD_2Cl_2$ (500 MHz)

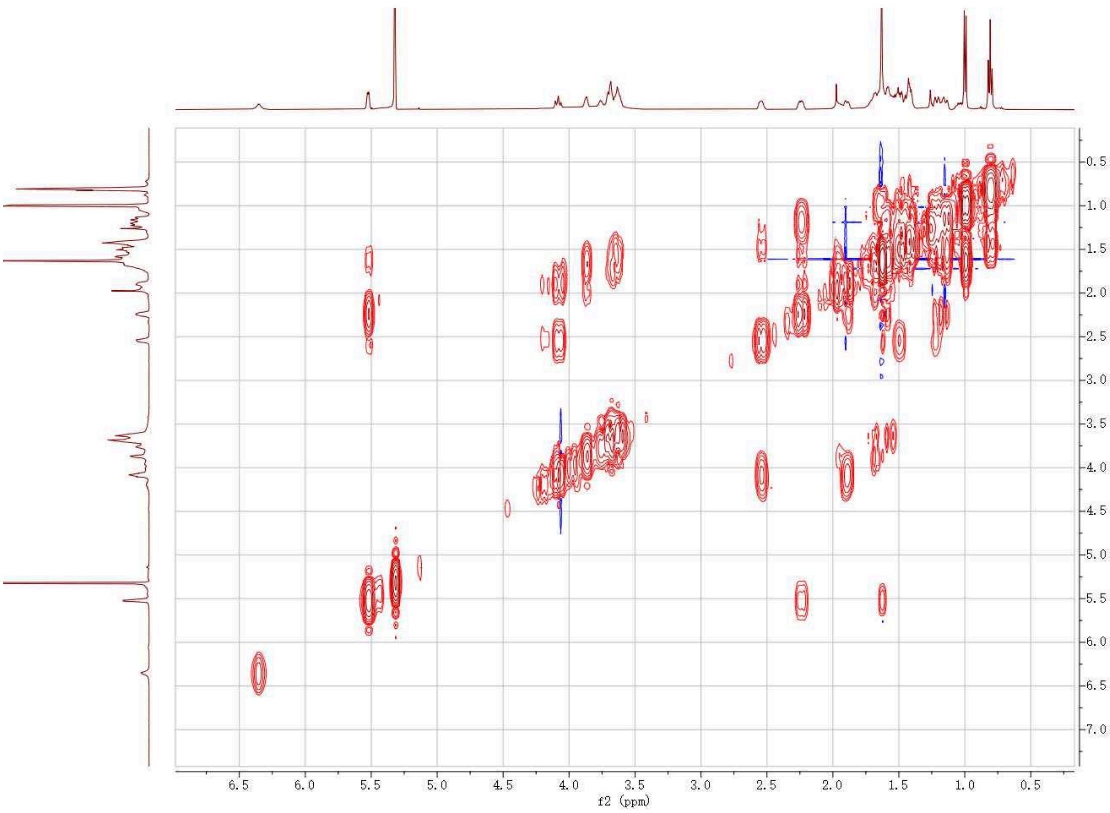

**Figure S31:** The $^1$H-$^1$H COSY NMR Spectrum of **12** in $CD_2Cl_2$(500 MHz)

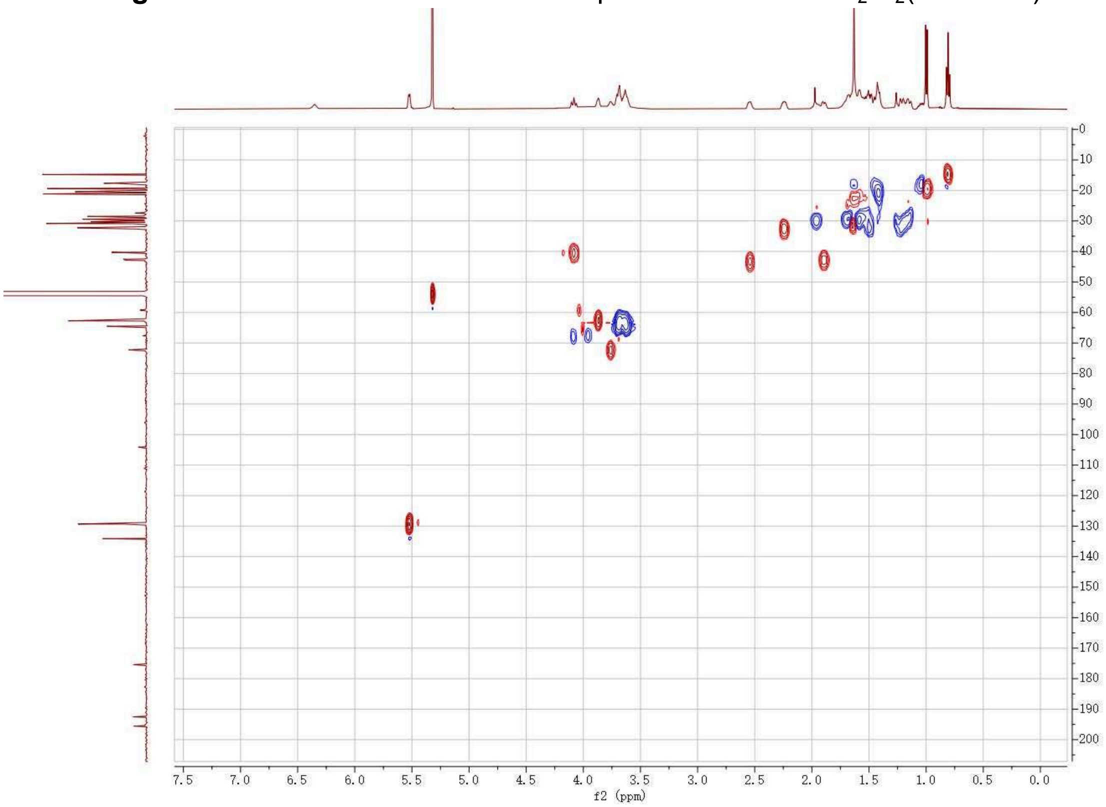

**Figure S32:** The $^1$H-$^{13}$C HSQC NMR Spectrum of **12** in $CD_2Cl_2$ (500 MHz)

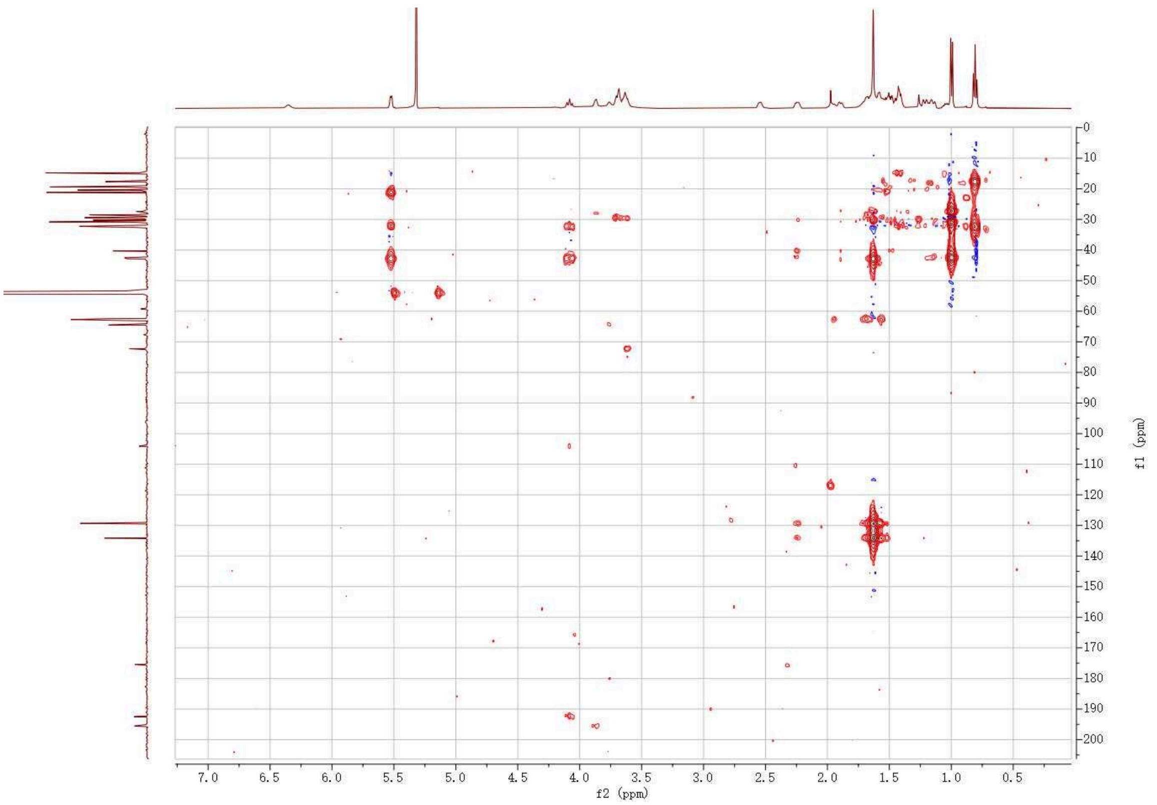

**Figure S33:** The $^{1}H$-$^{13}C$ HMBC NMR Spectrum of **12** in $CD_2Cl_2$ (500 MHz)

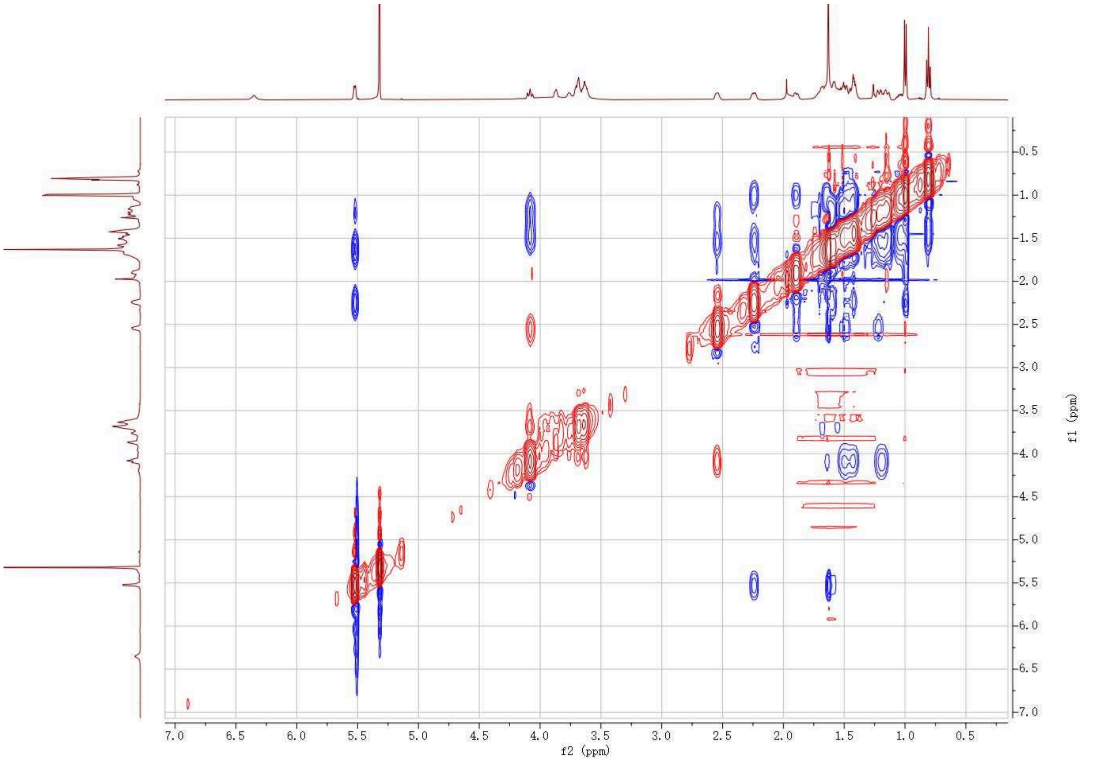

**Figure S34:** The $^{1}H$-$^{1}H$ NOESY NMR Spectrum of **12** in $CD_2Cl_2$ (500 MHz)

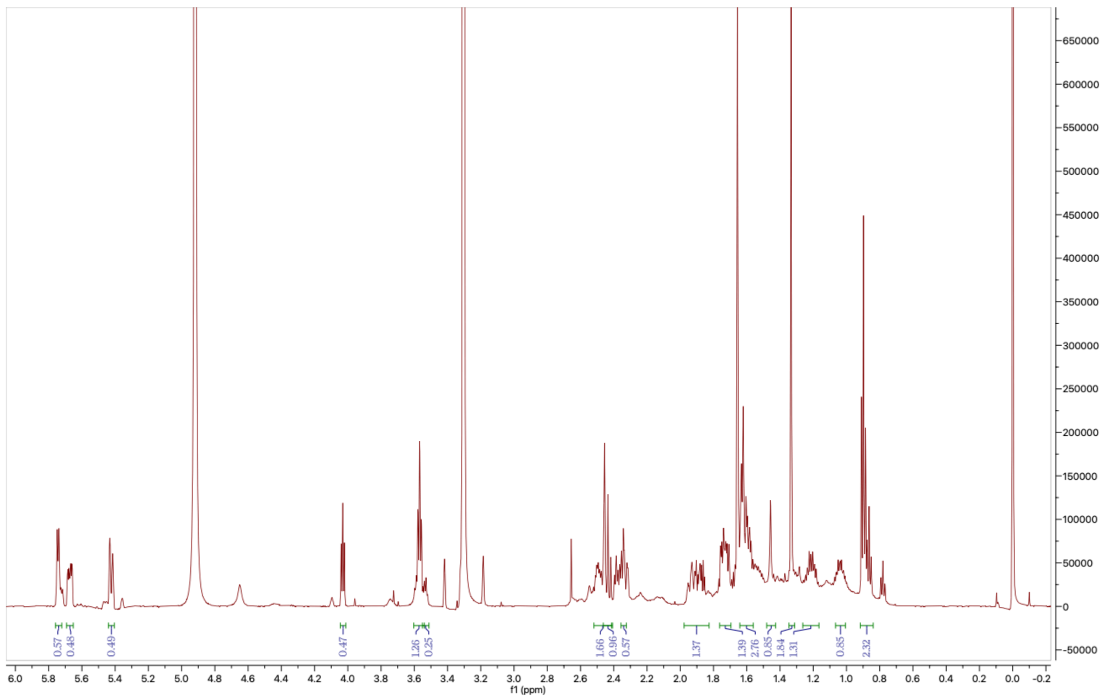

**Figure S35:** The $^{1}H$ NMR Spectrum of **13** in MeOD (600MHz)

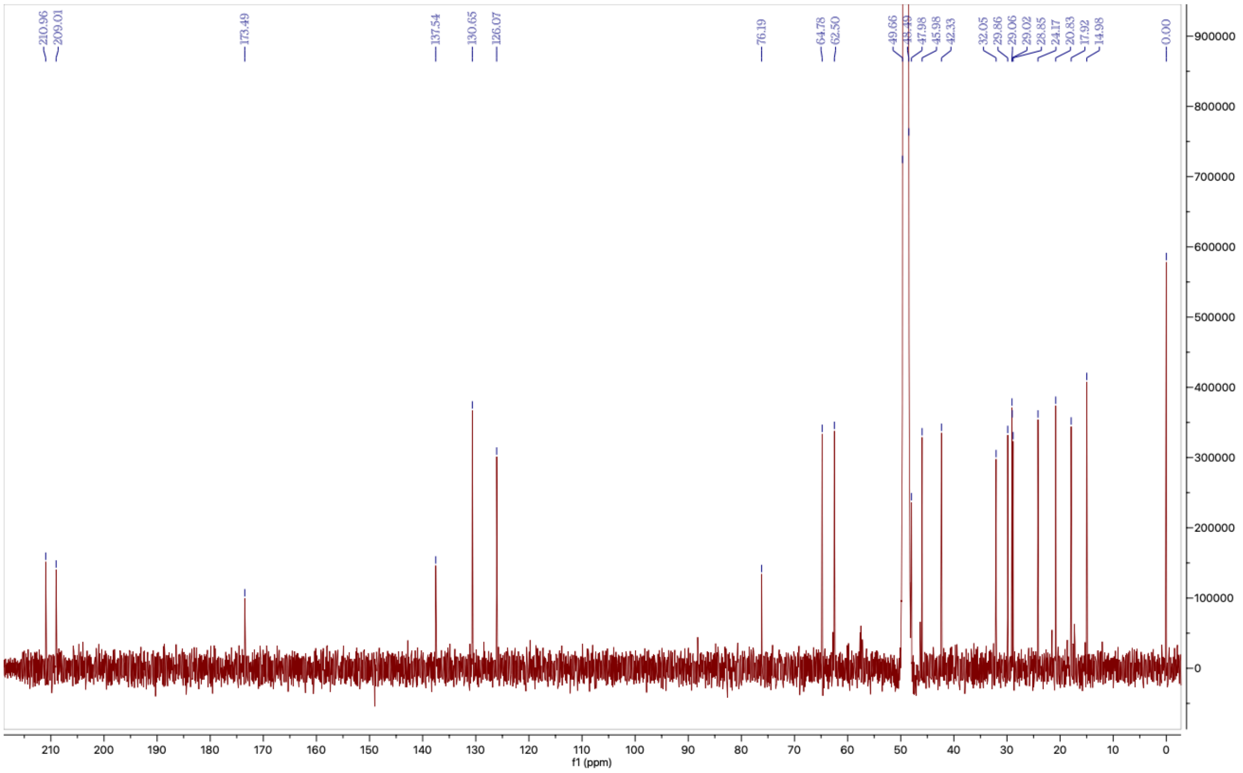

**Figure S36:** The $^{13}C$ NMR Spectrum of **13** in MeOD (150MHz)

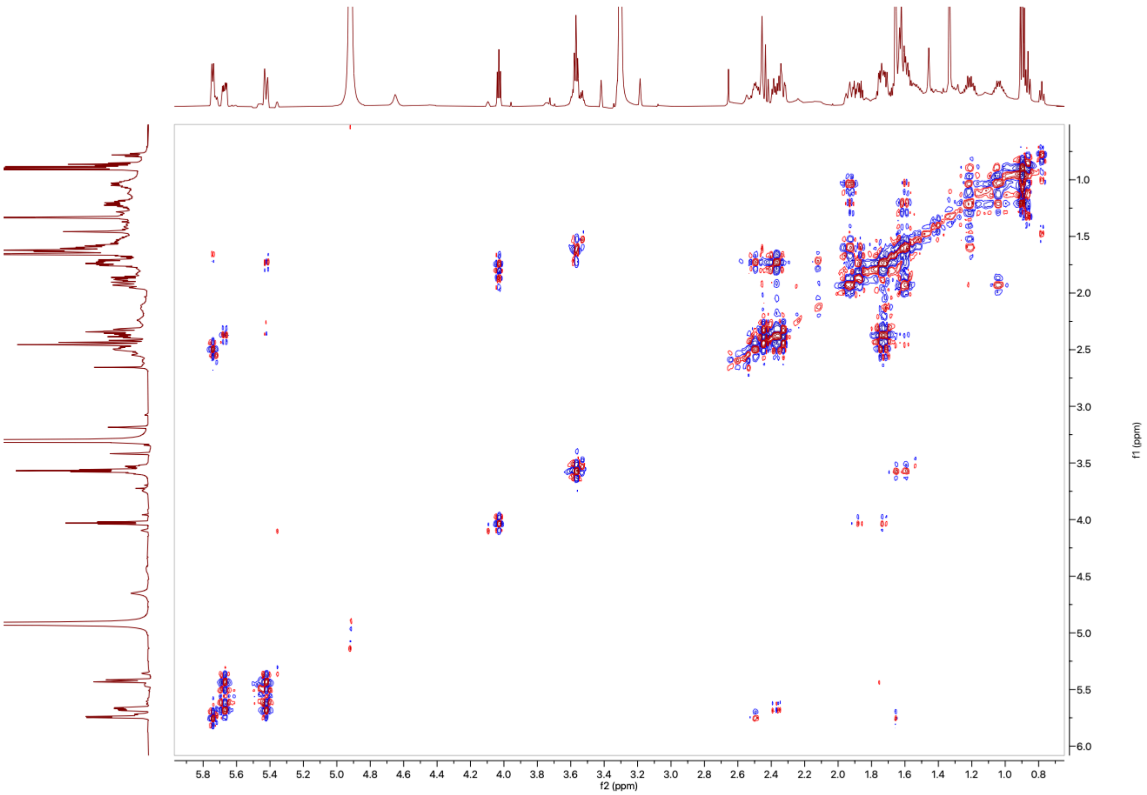

**Figure S37:** The [1]H-[1]H COSY NMR Spectrum of **13** in MeOD (600 MHz)

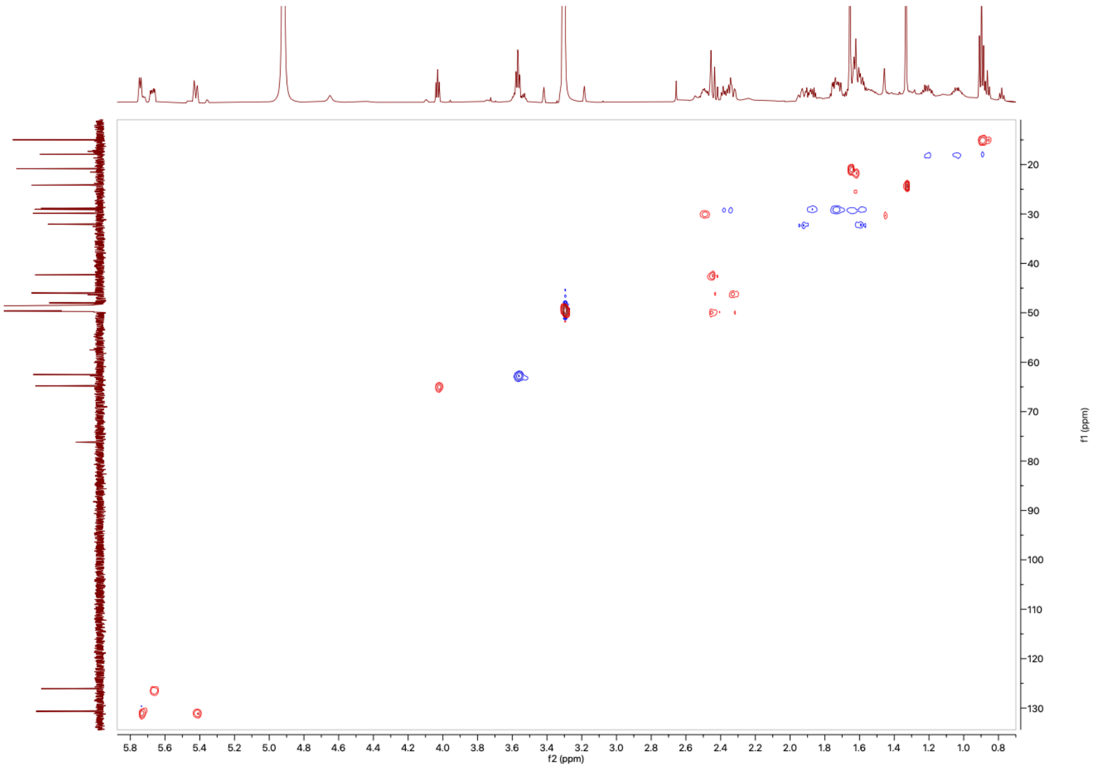

**Figure S38:** The [1]H-[13]C HSQC NMR Spectrum of **13** in MeOD (600 MHz)

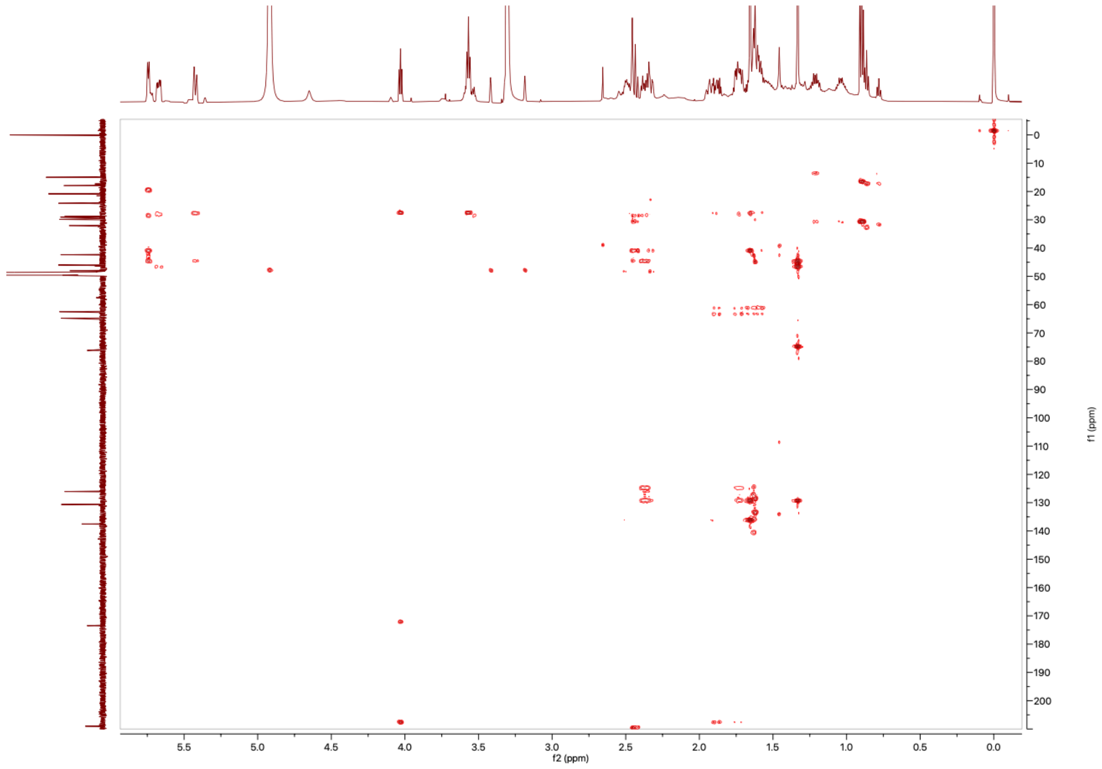

**Figure S39:** The $^{1}H$-$^{13}C$ HMBC NMR Spectrum of **13** in MeOD (600 MHz)

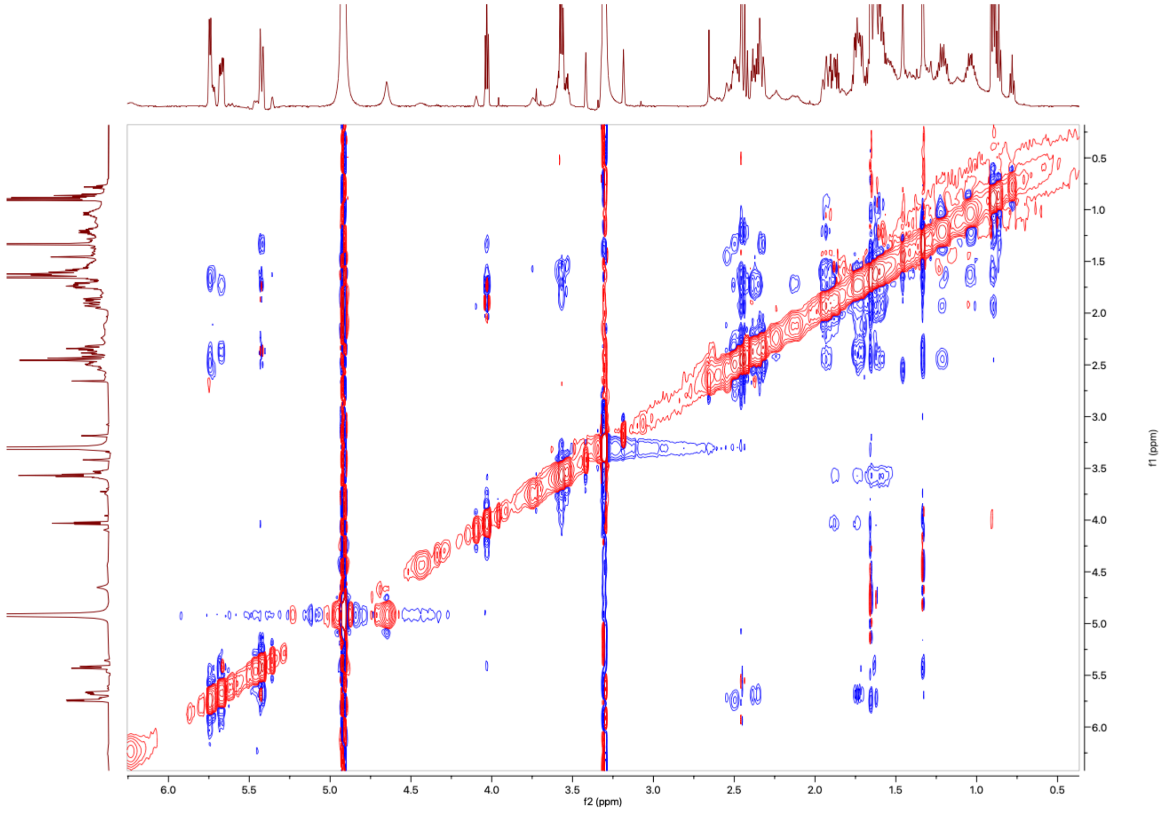

**Figure S40:** The $^{1}H$-$^{1}H$ NOESY NMR Spectrum of **13** in MeOD (600 MHz)

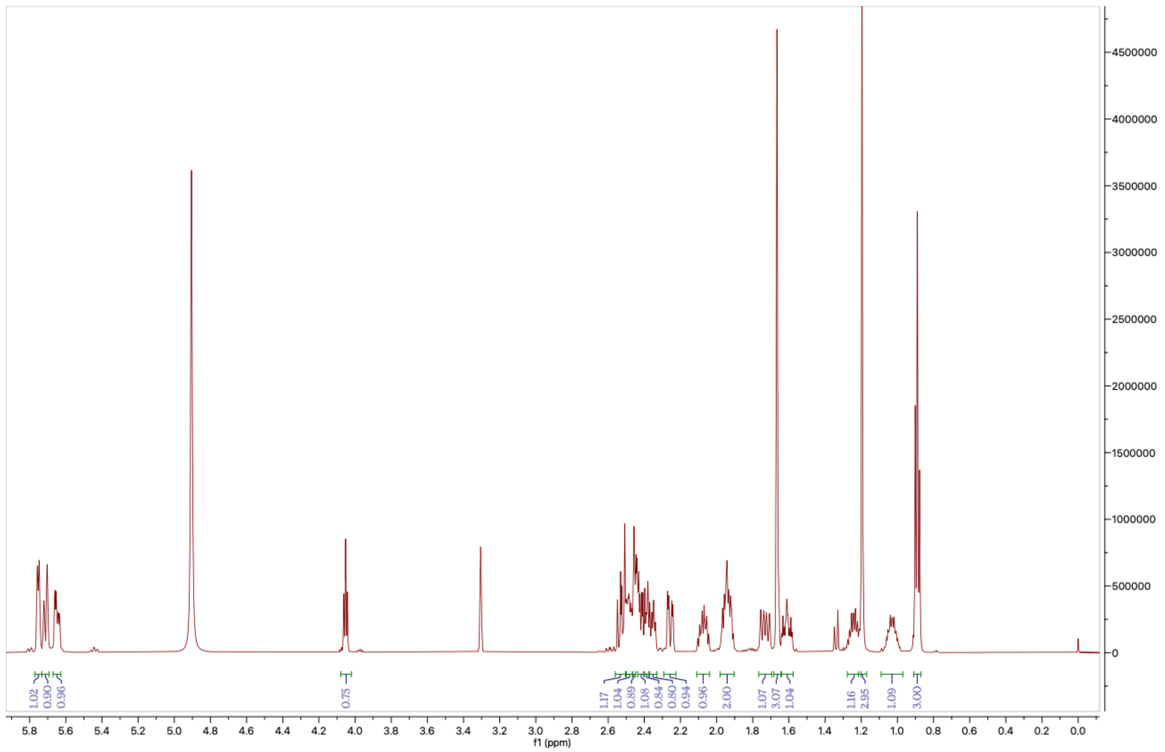

**Figure S41:** The ¹H NMR Spectrum of **14** in MeOD (600MHz)

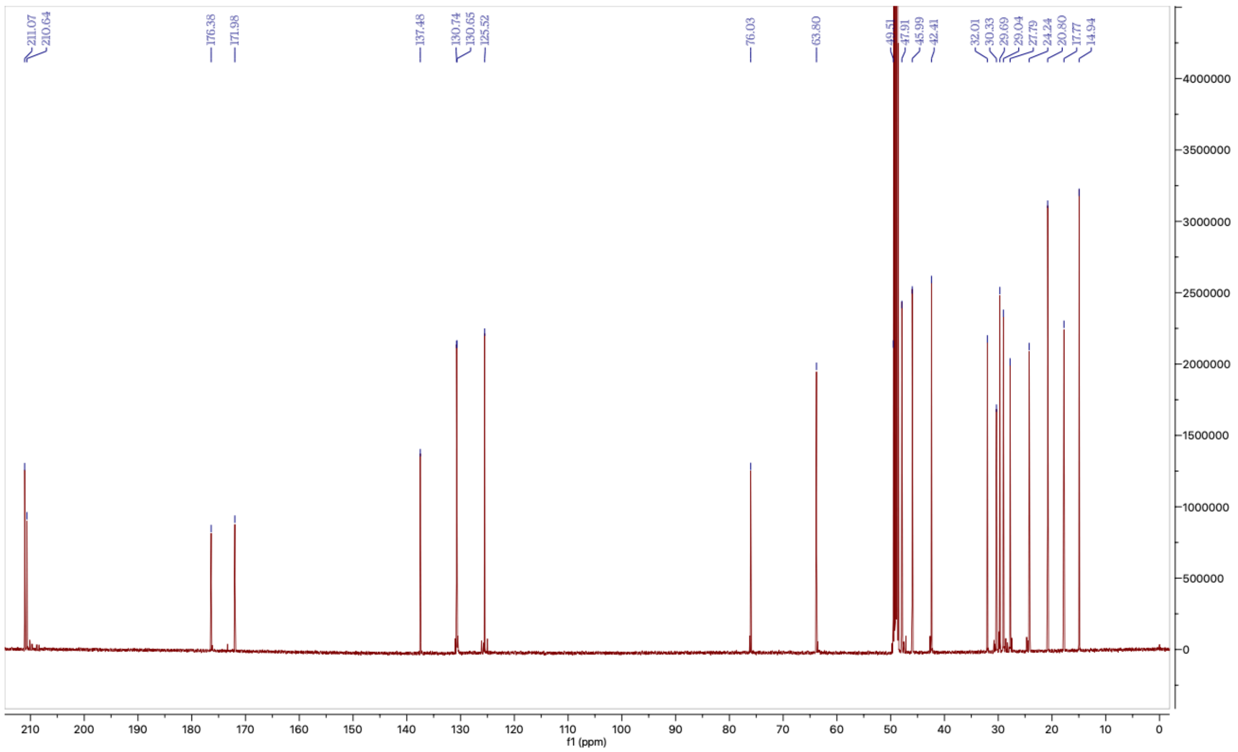

**Figure S42:** The ¹³C NMR Spectrum of **14** in MeOD (600MHz)

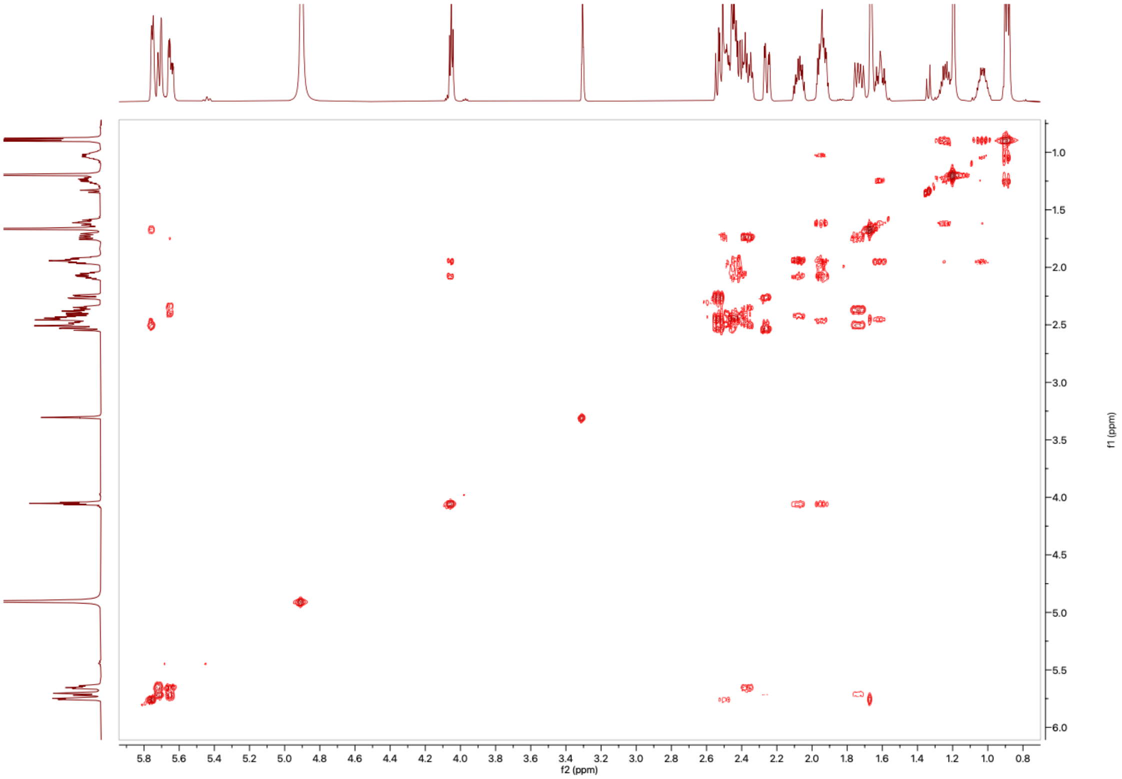

**Figure S43:** The $^{1}H$-$^{1}H$ COSY NMR Spectrum of **14** in MeOD (600 MHz)

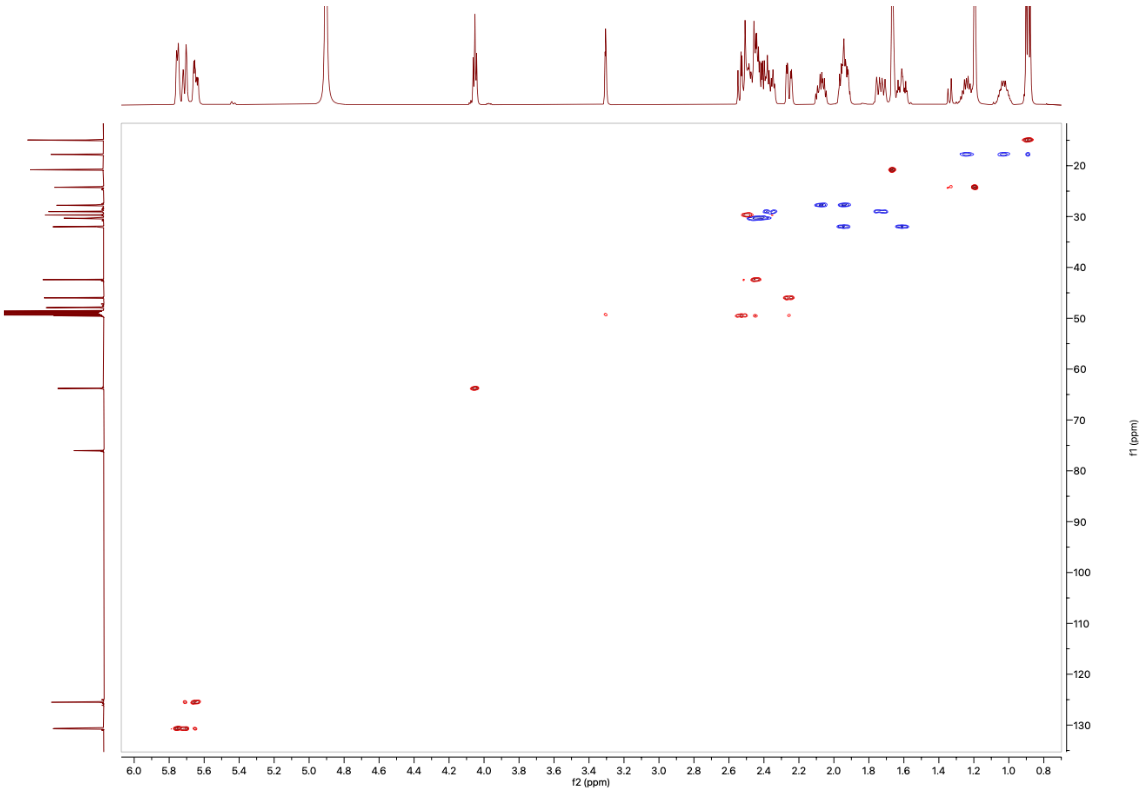

**Figure S44:** The $^{1}H$-$^{13}C$ HSQC NMR Spectrum of **14** in MeOD (600 MHz)

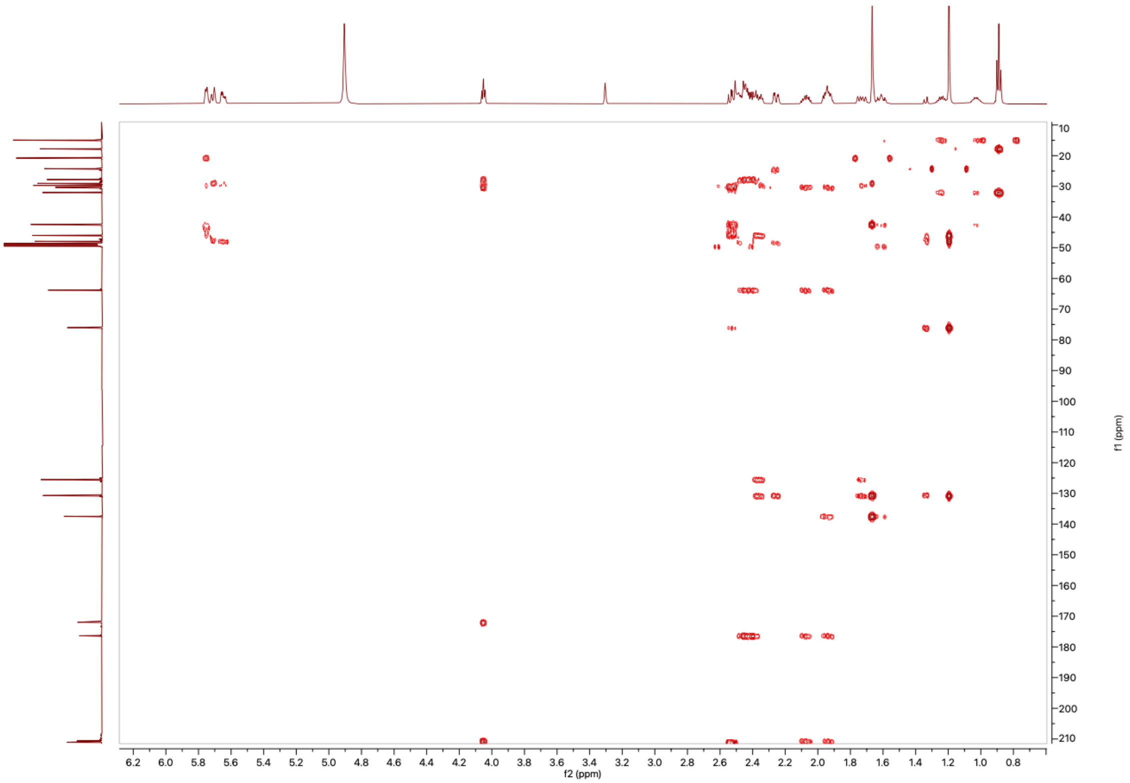

**Figure S45:** The $^{1}H$-$^{13}C$ HMBC NMR Spectrum of **14** in MeOD (600 MHz)

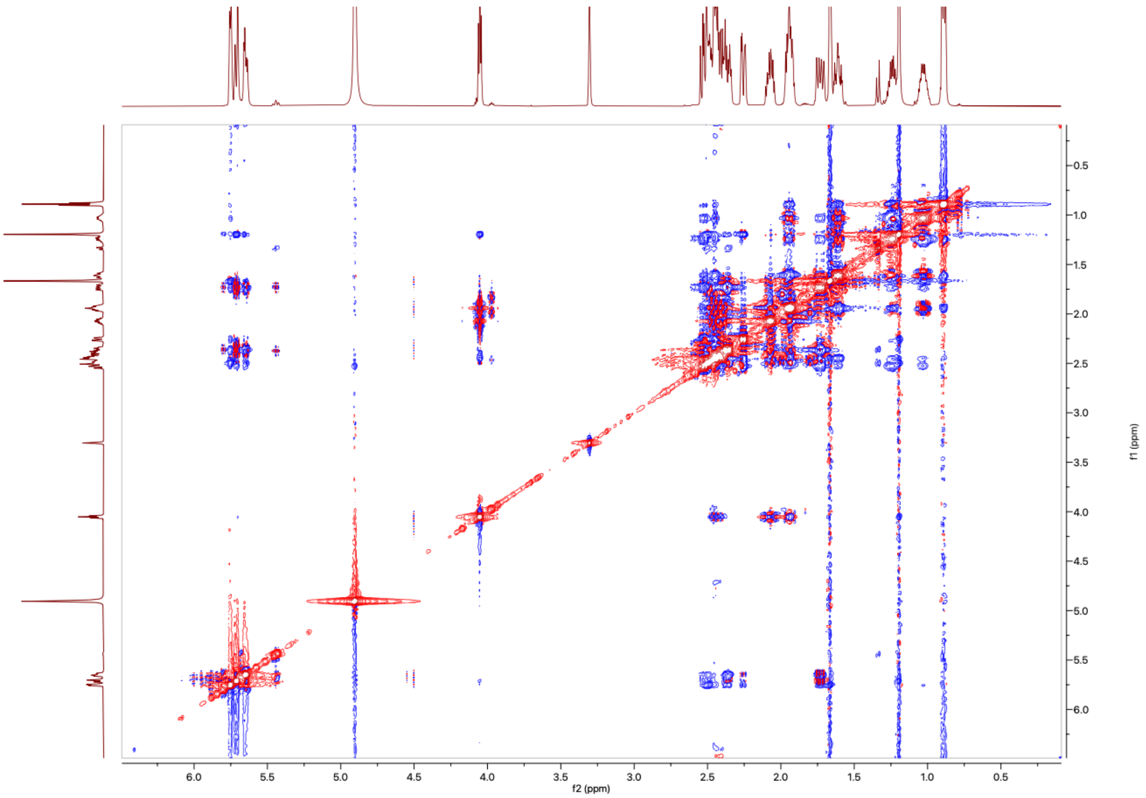

**Figure S46:** The $^{1}H$-$^{1}H$ NOESY NMR Spectrum of **14** in MeOD (600 MHz)

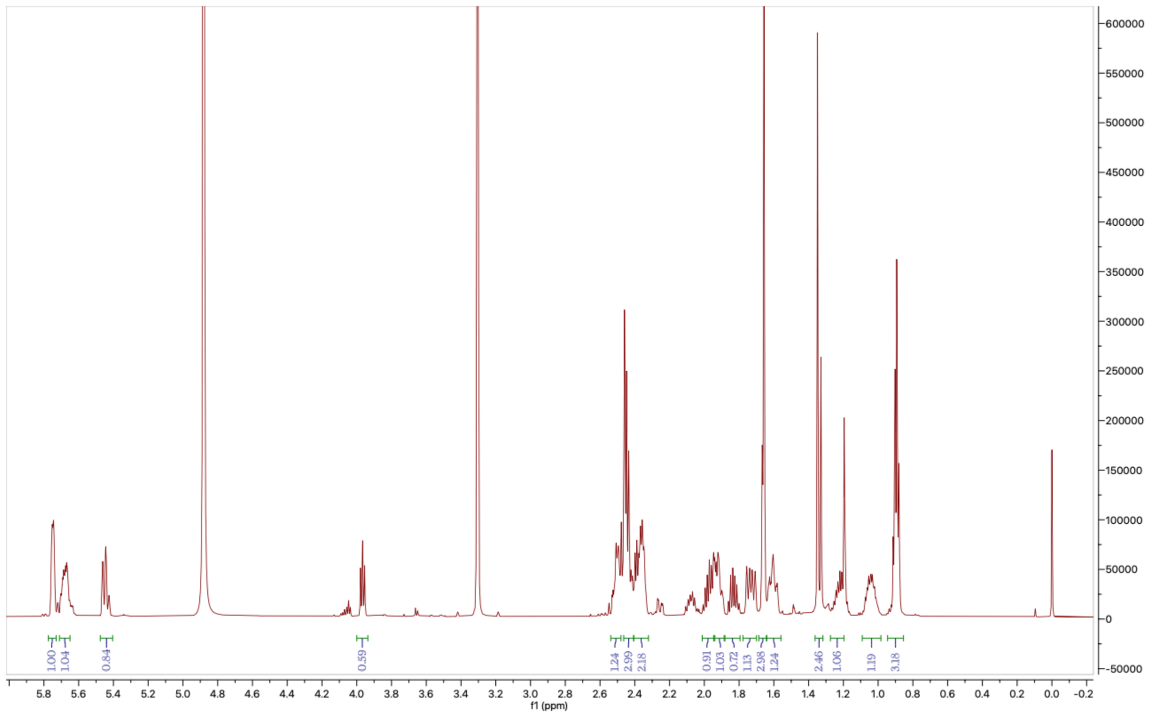

**Figure S47:** The $^{1}$H NMR Spectrum of **15** in MeOD (600MHz)

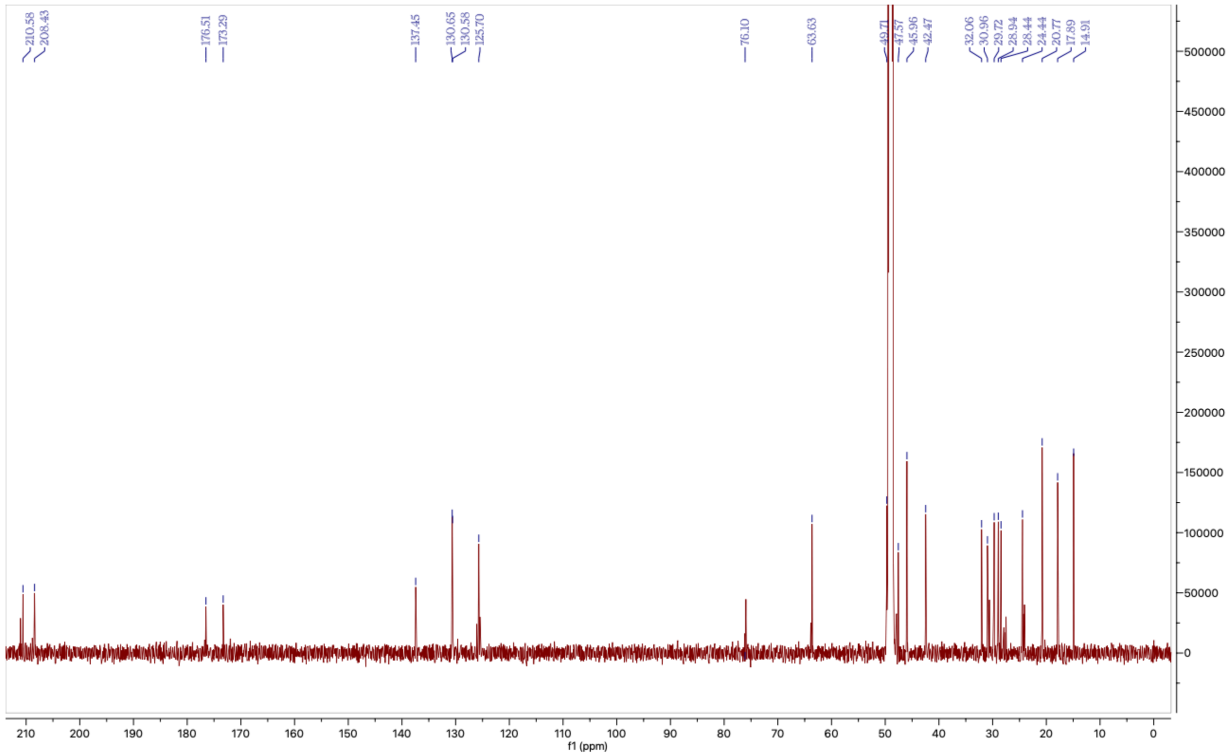

**Figure S48:** The $^{13}$C NMR Spectrum of **15** in MeOD (600MHz)

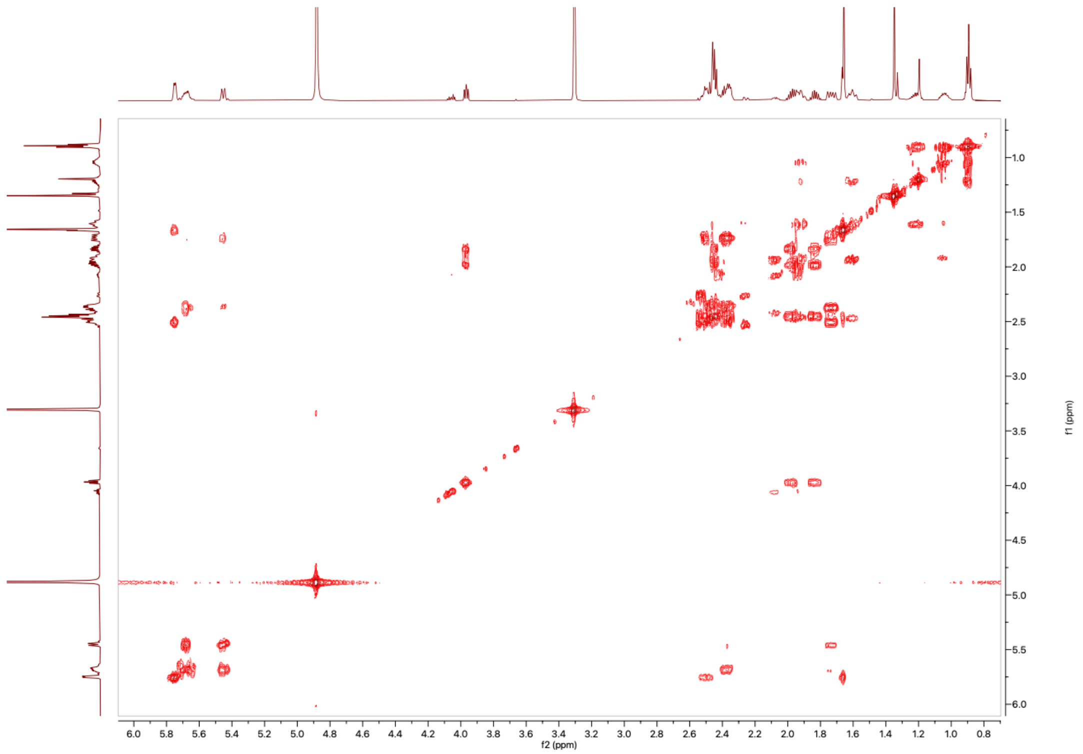

**Figure S49:** The ¹H-¹H COSY NMR Spectrum of **15** in MeOD (600 MHz)

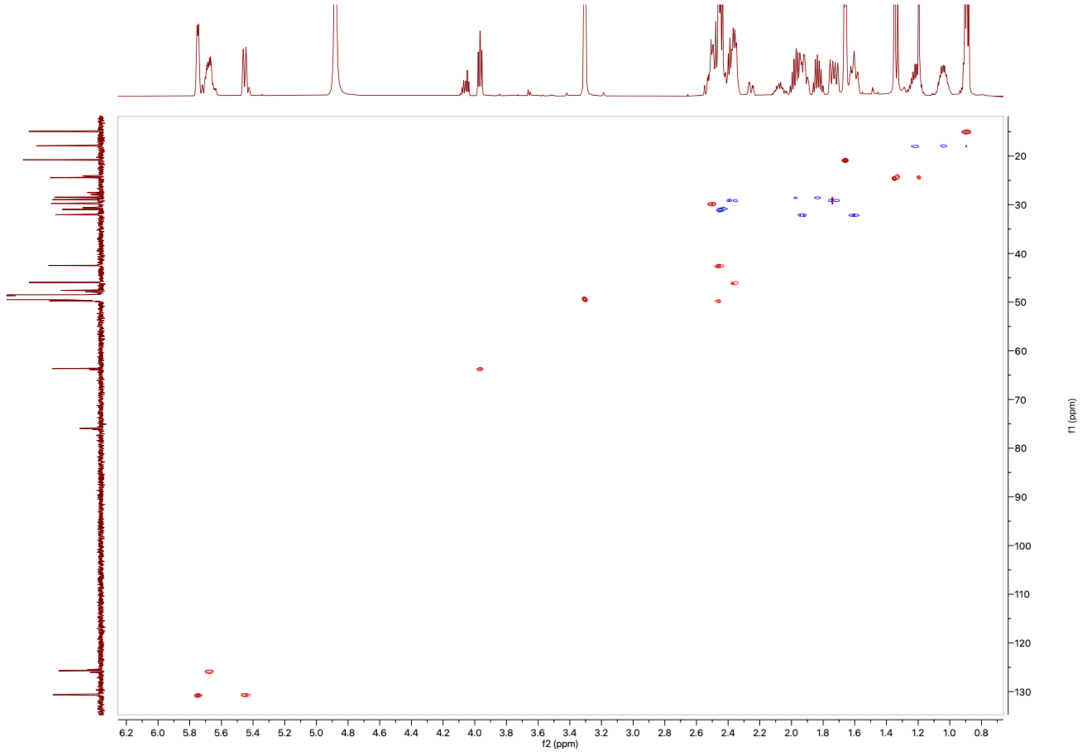

**Figure S50:** The ¹H-¹³C HSQC NMR Spectrum of **15** in MeOD (600 MHz)

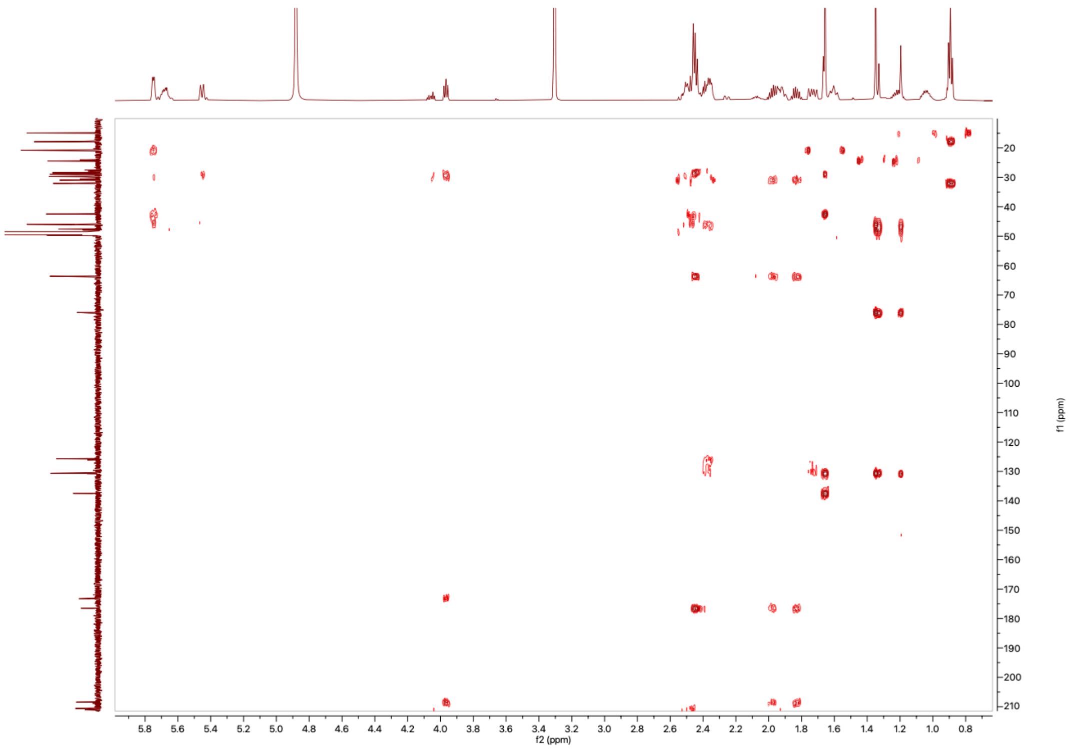

**Figure S51:** The $^{1}H$-$^{13}C$ HMBC NMR Spectrum of **15** in MeOD (600 MHz)

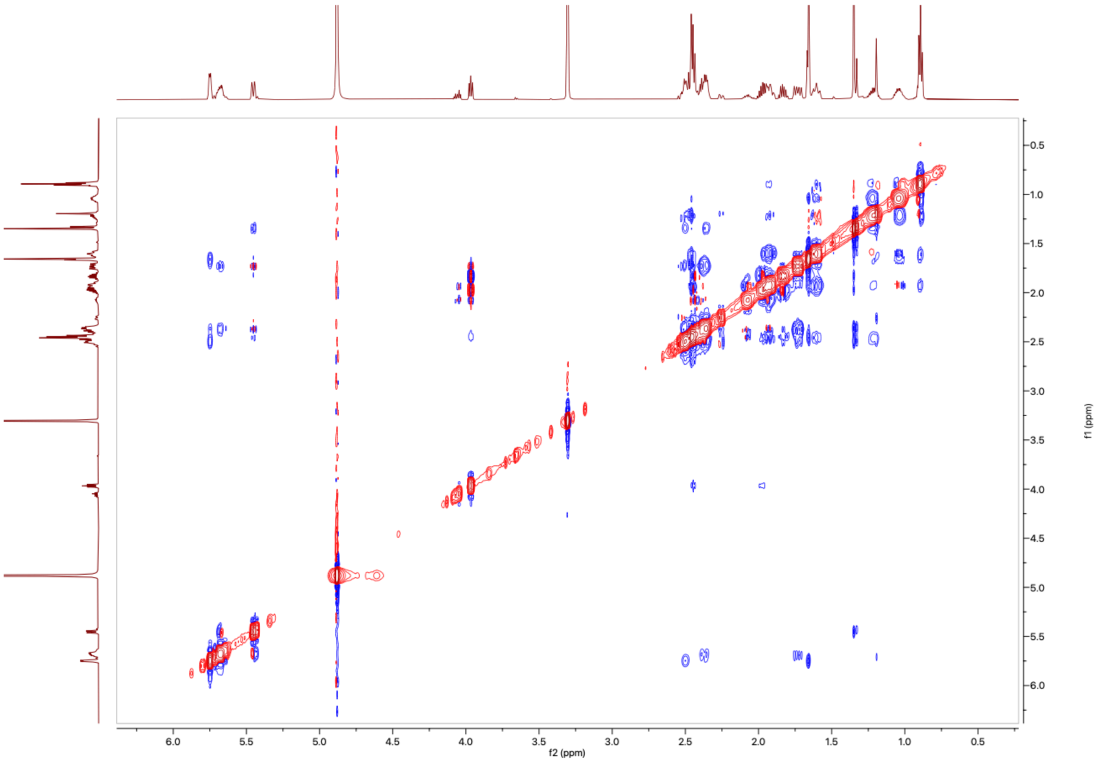

**Figure S52:** The $^{1}H$-$^{1}H$ NOESY NMR Spectrum of **15** in MeOD (600 MHz)

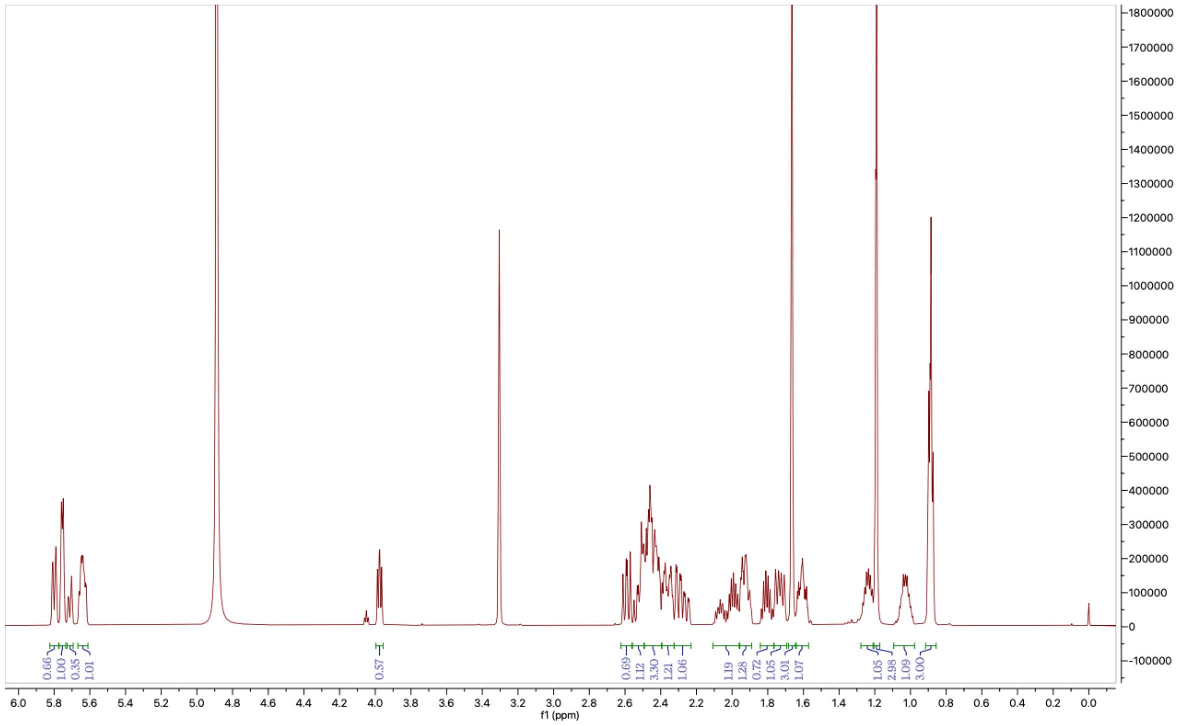

**Figure S53:** The ¹H NMR Spectrum of **16** in MeOD (600MHz)

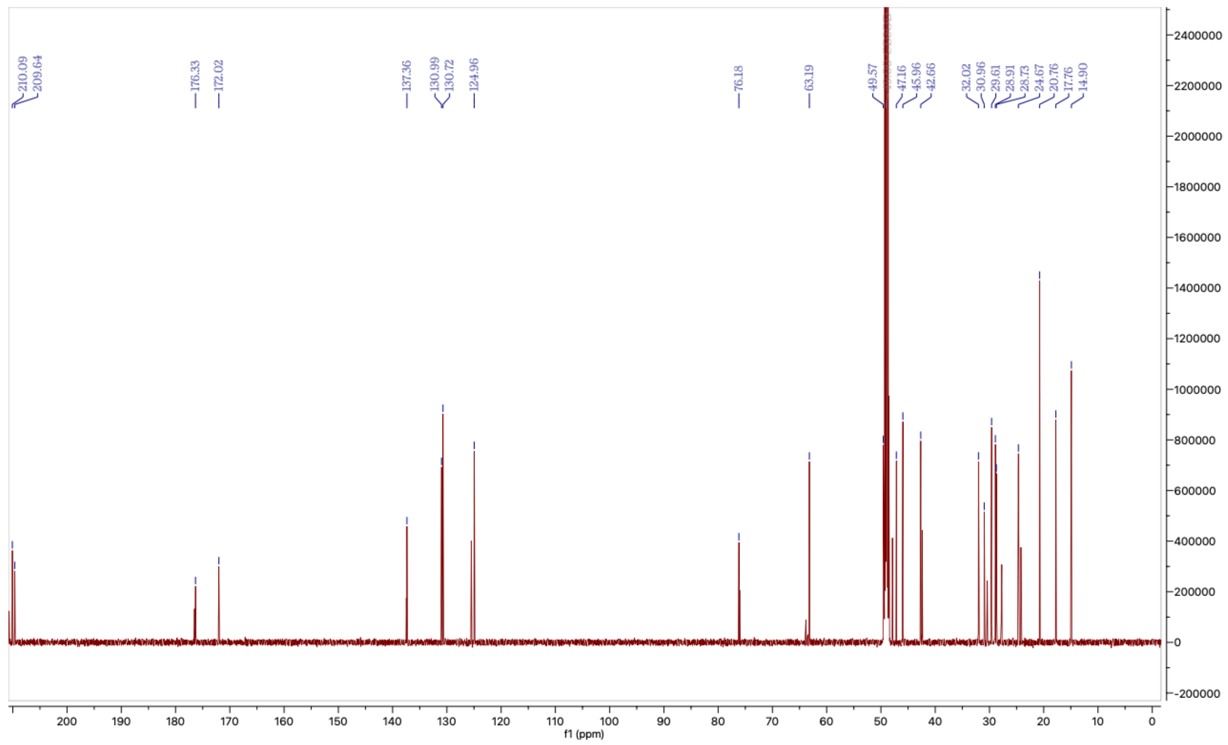

**Figure S54:** The ¹³C NMR Spectrum of **16** in MeOD (600MHz)

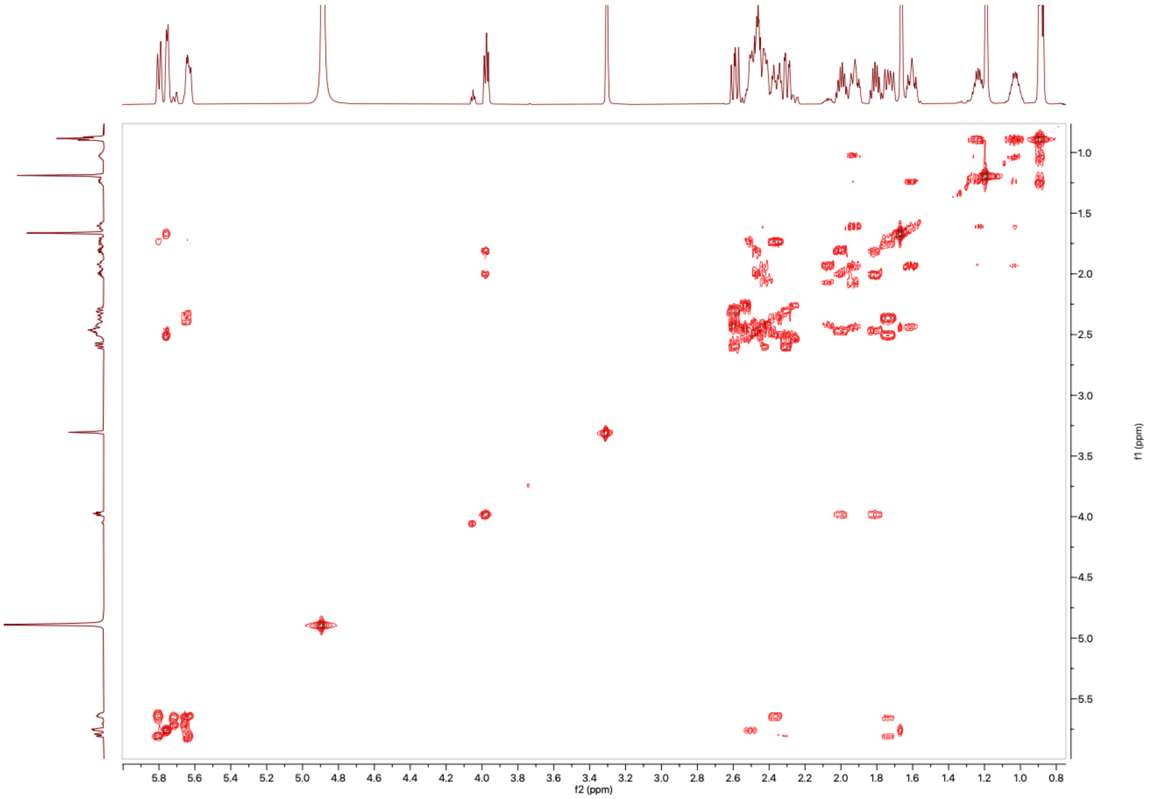

**Figure S55:** The $^1H$-$^1H$ COSY NMR Spectrum of **16** in MeOD (600 MHz)

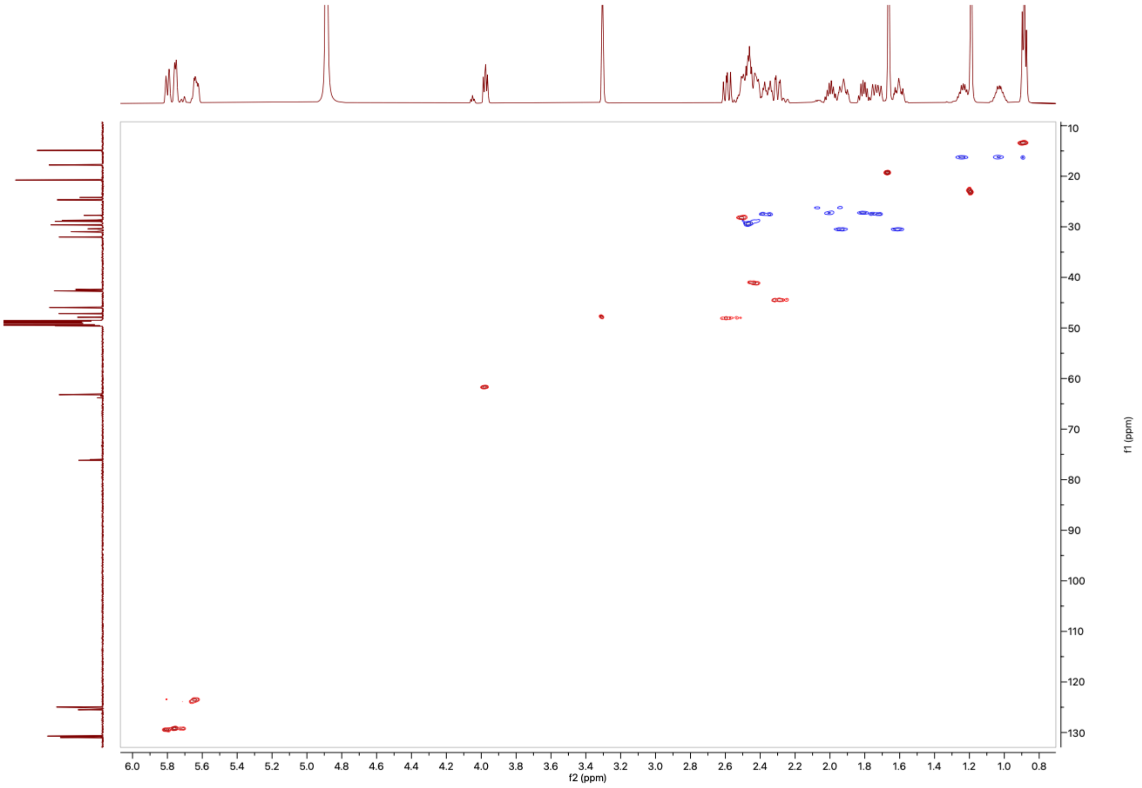

**Figure S56:** The $^1H$-$^{13}C$ HSQC NMR Spectrum of **16** in MeOD (600 MHz)

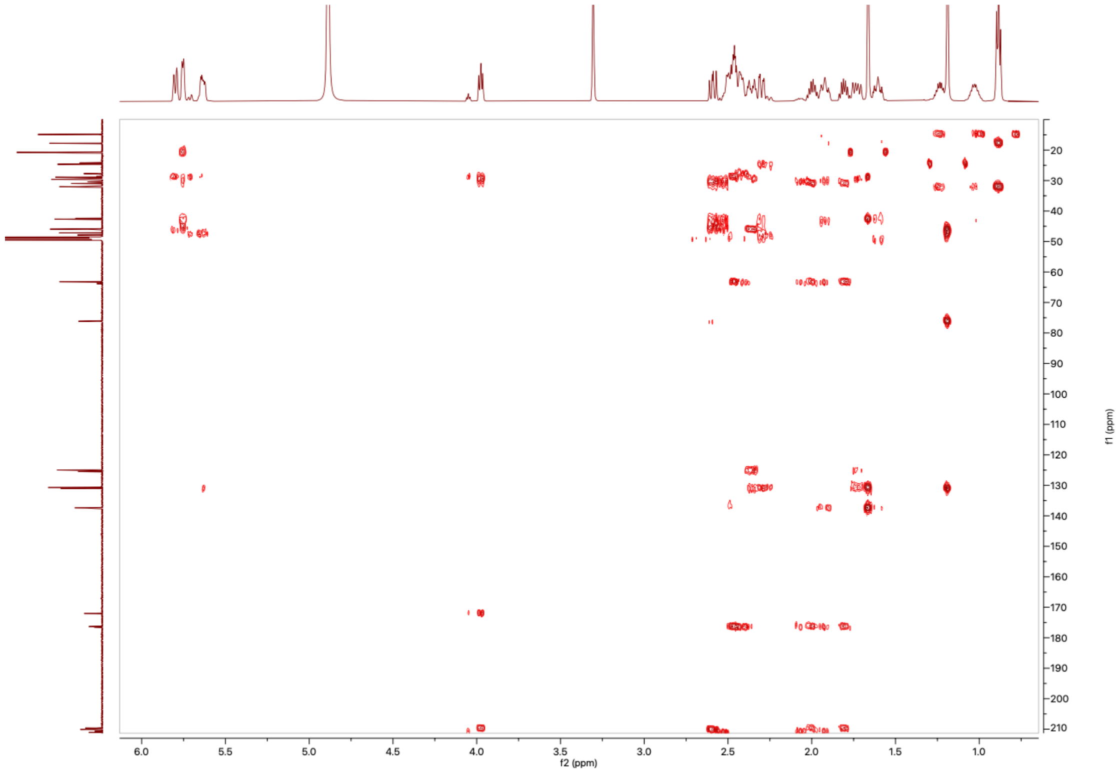

**Figure S57:** The ¹H-¹³C HMBC NMR Spectrum of **16** in MeOD (600 MHz)

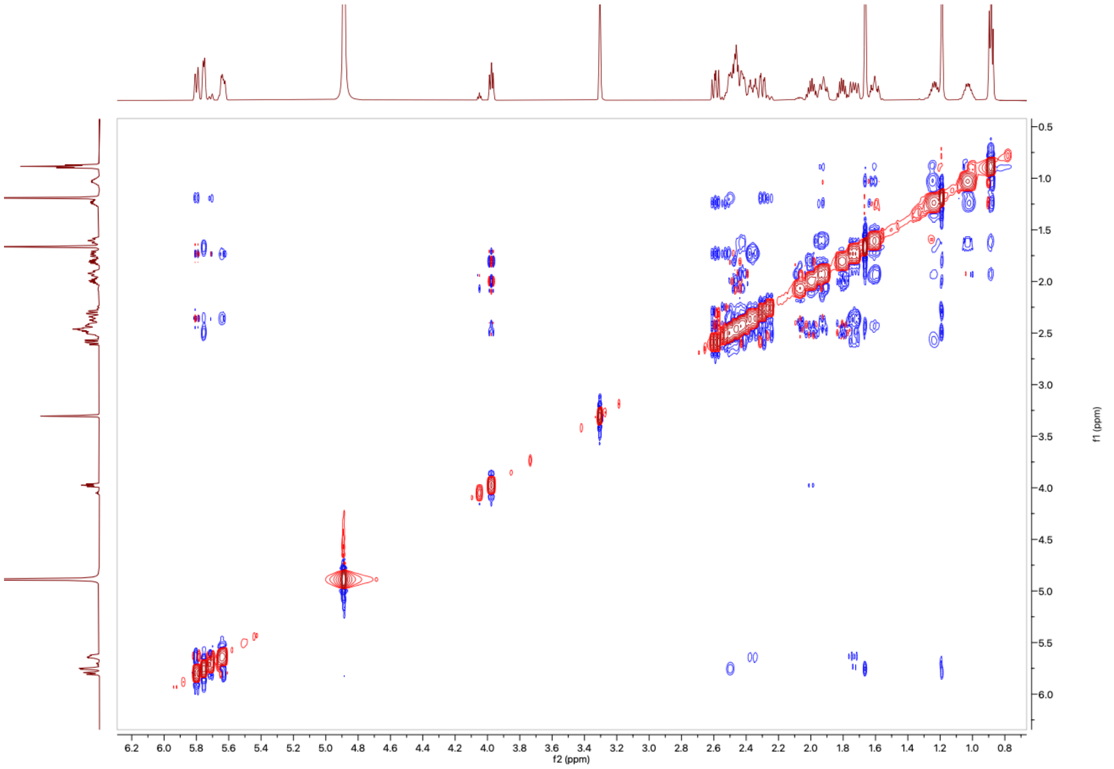

**Figure S58:** The ¹H-¹H NOESY NMR Spectrum of **16** in MeOD (600 MHz)

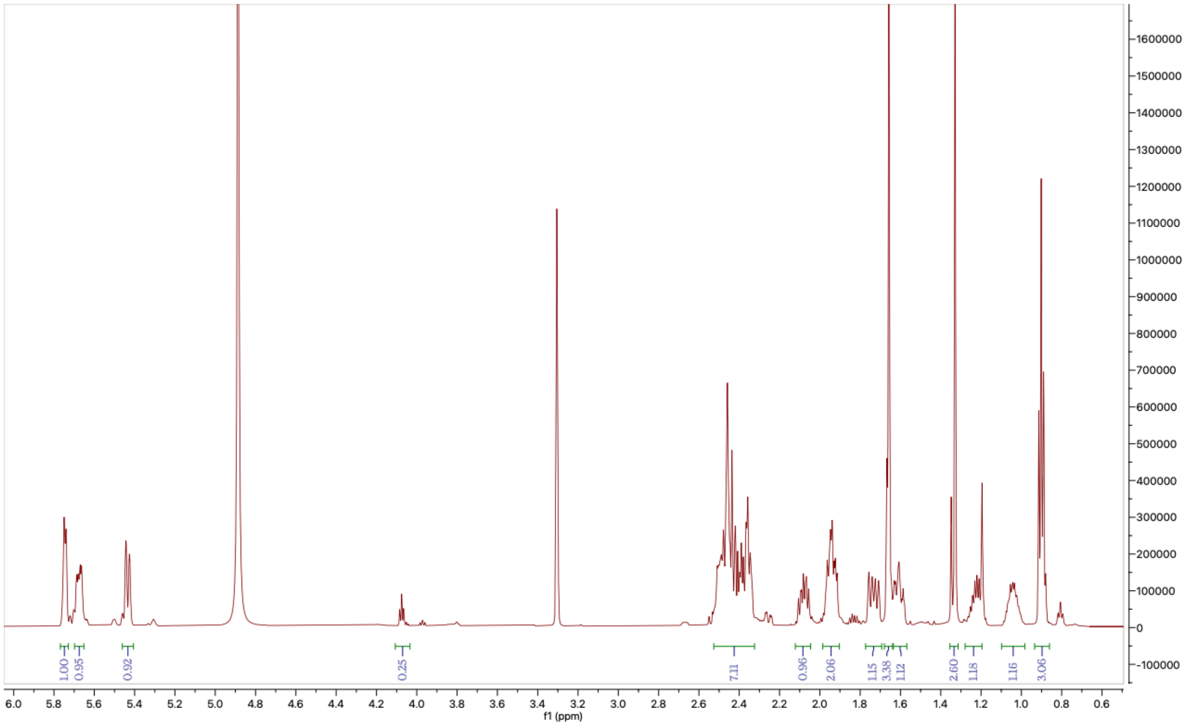

**Figure S59:** The $^1$H NMR Spectrum of **17** in MeOD (600MHz)

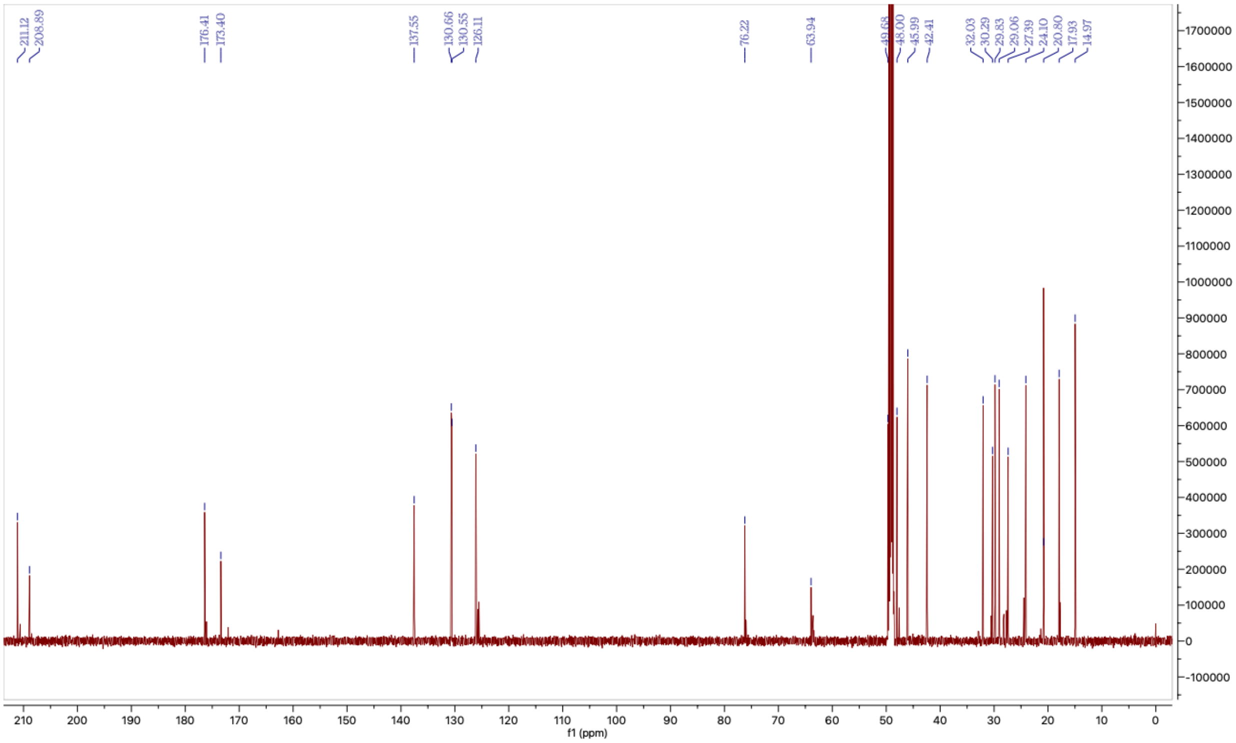

**Figure S60:** The $^{13}$C NMR Spectrum of **17** in MeOD (600MHz)

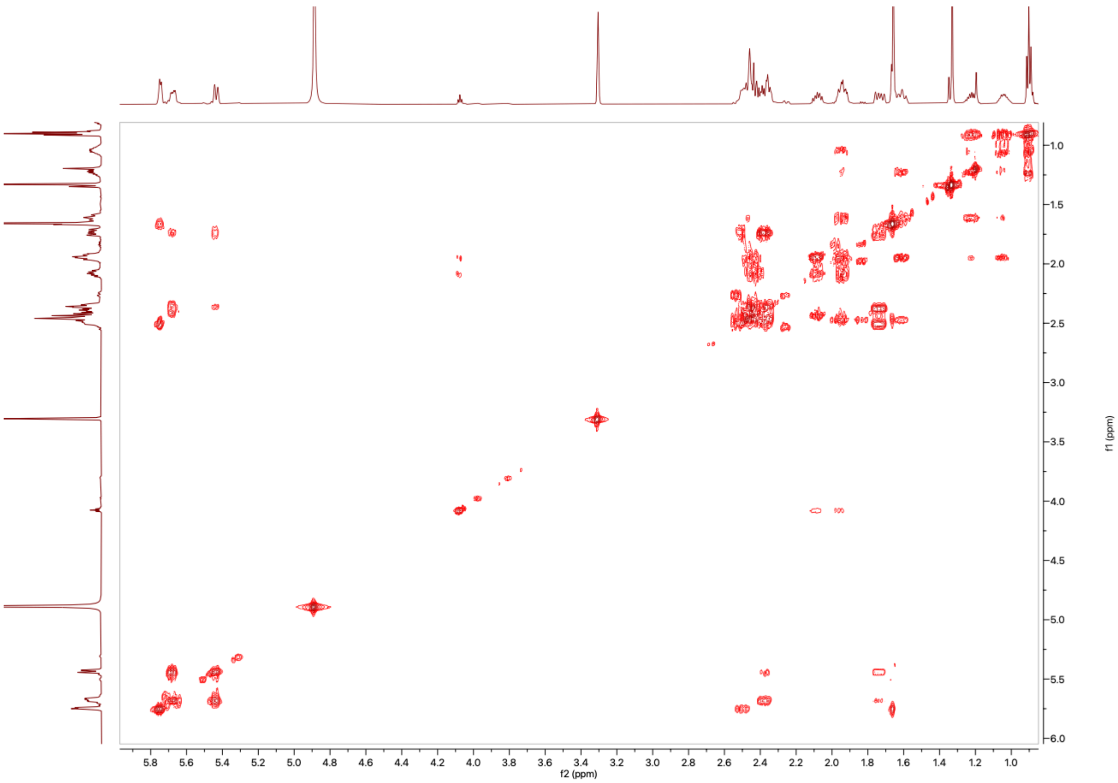

**Figure S61:** The $^1$H-$^1$H COSY NMR Spectrum of **17** in MeOD (600 MHz)

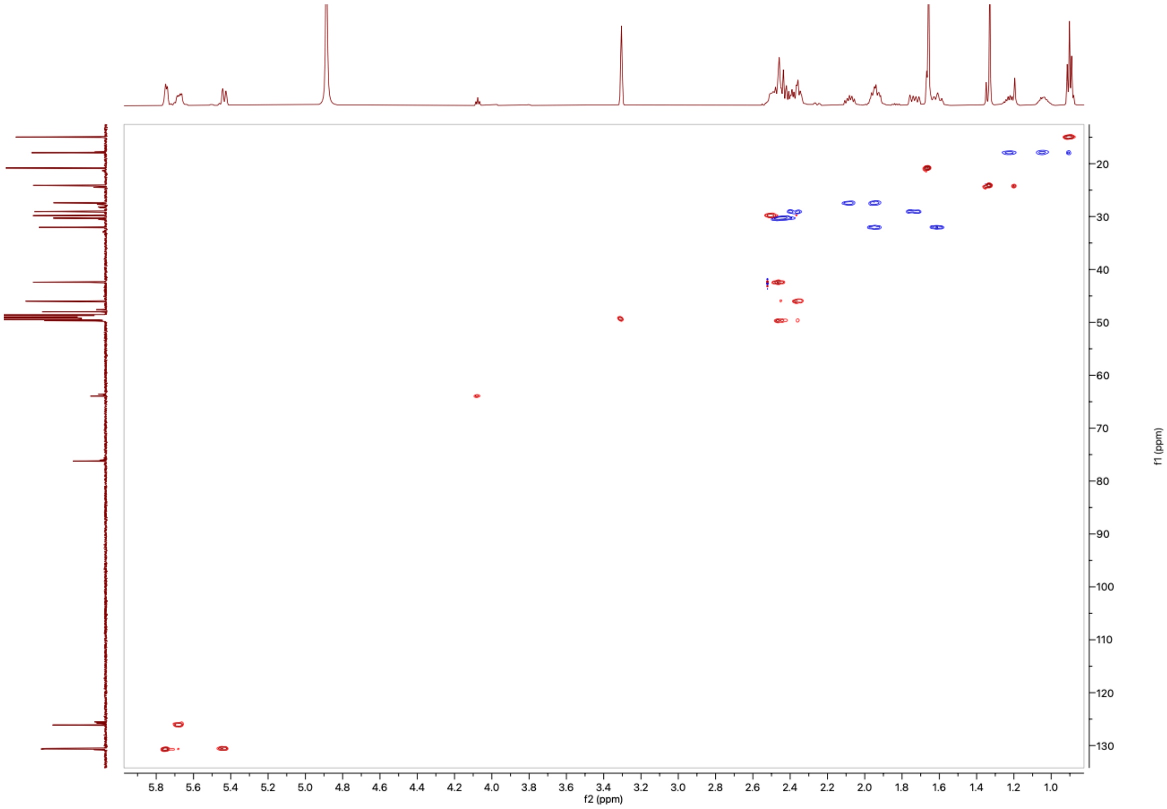

**Figure S62:** The $^1$H-$^{13}$C HSQC NMR Spectrum of **17** in MeOD (600 MHz)

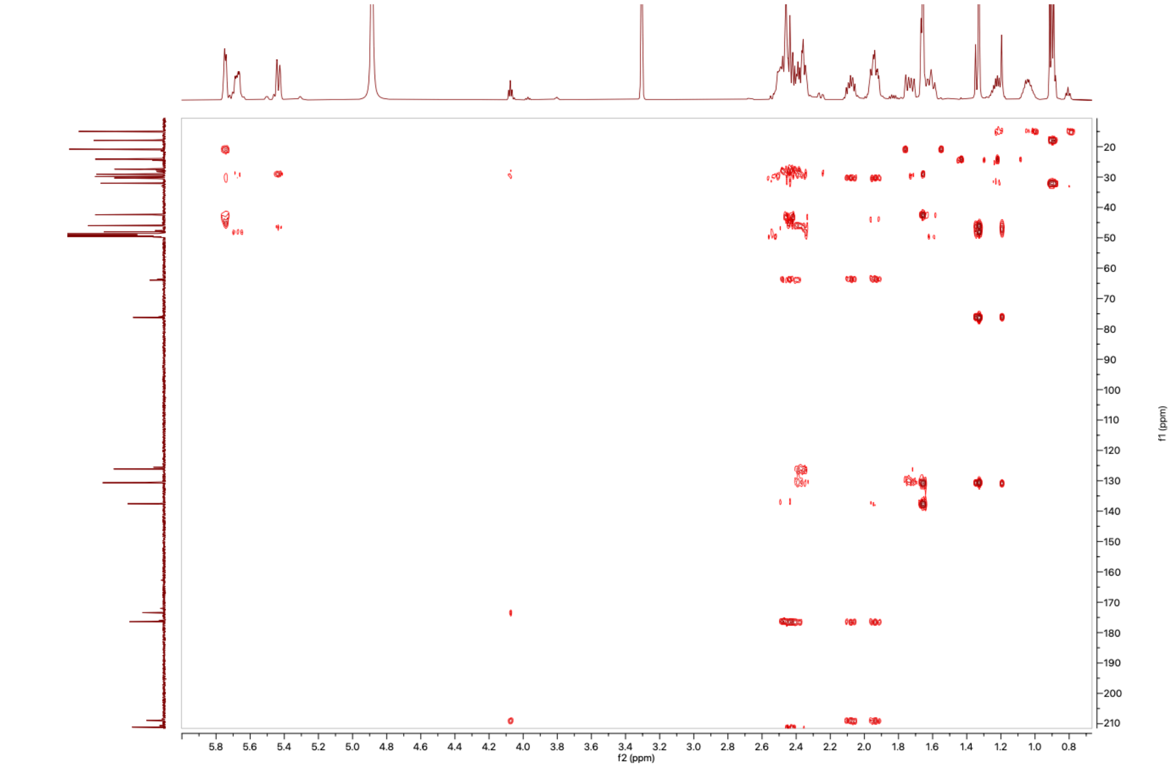

**Figure S63:** The $^{1}H$-$^{13}C$ HMBC NMR Spectrum of **17** in MeOD (600 MHz)

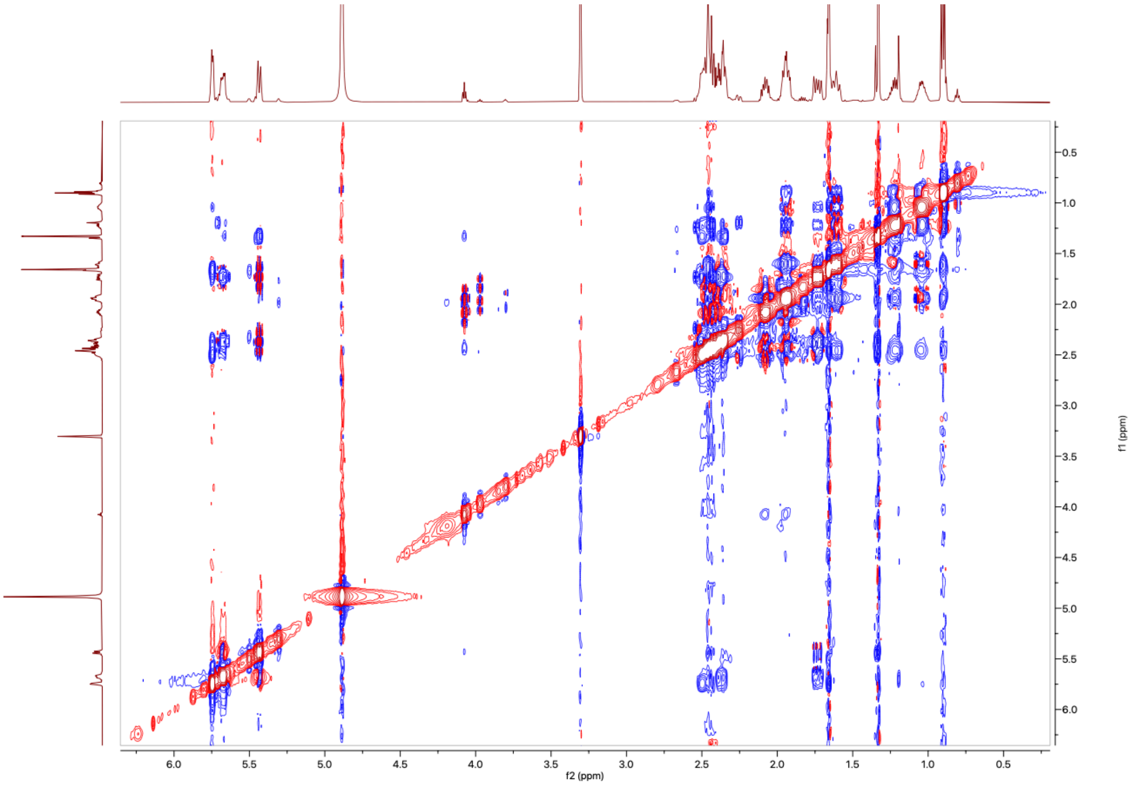

**Figure S64:** The $^{1}H$-$^{1}H$ NOESY NMR Spectrum of **17** in MeOD (600 MHz)

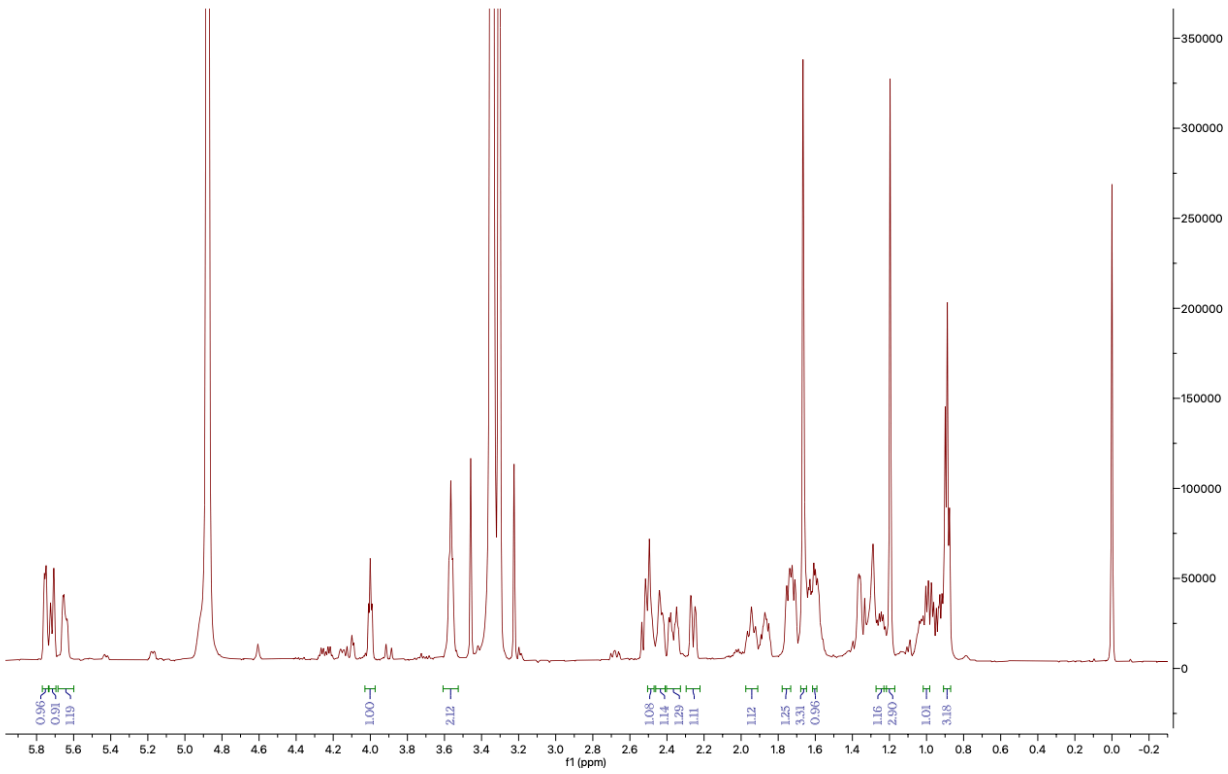

**Figure S65:** The $^{1}H$ NMR Spectrum of **19** in MeOD (600MHz)

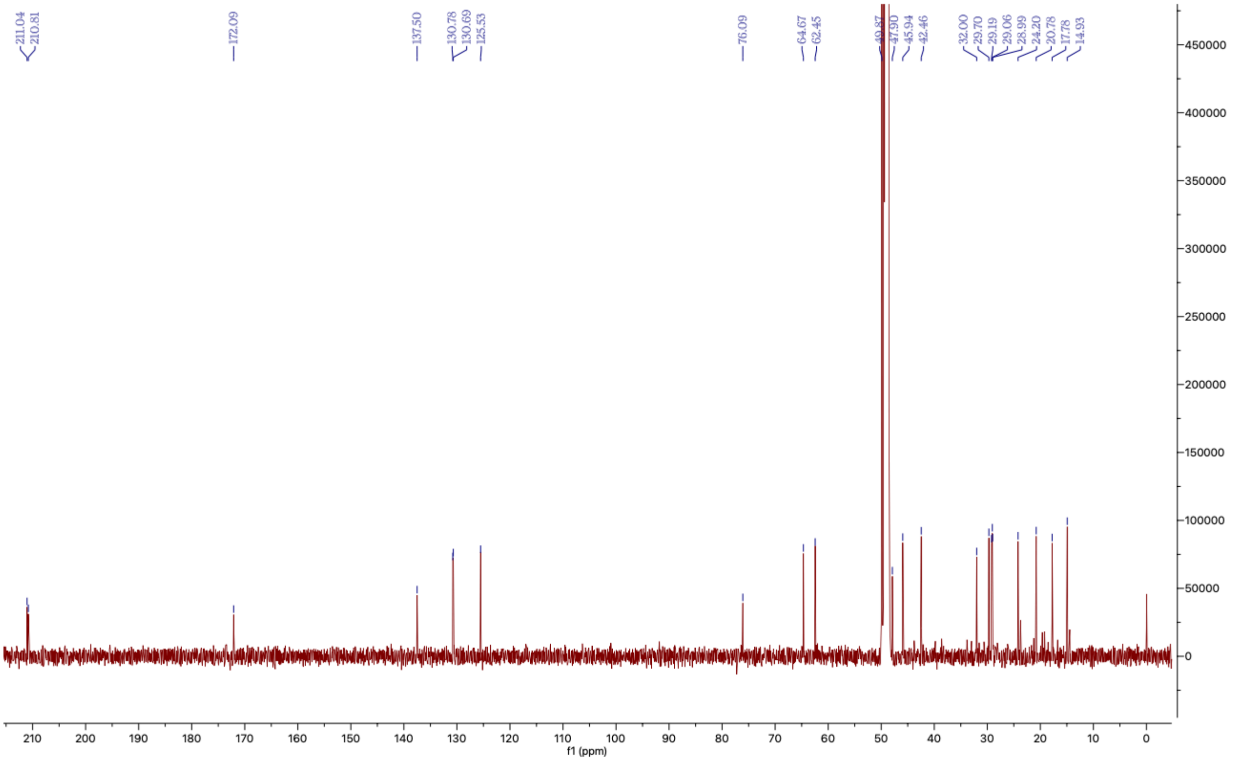

**Figure S66:** The $^{13}C$ NMR Spectrum of **19** in MeOD (150MHz)

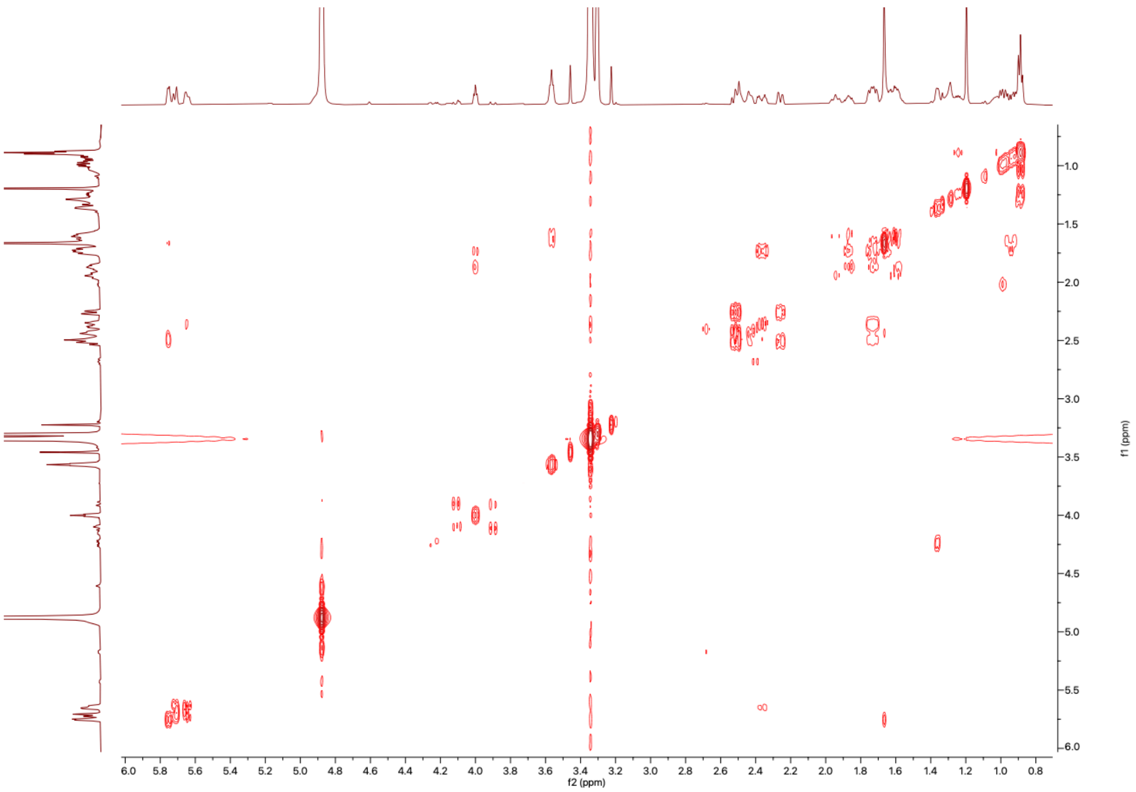

**Figure S67:** The $^1H$-$^1H$ COSY NMR Spectrum of **19** in MeOD (600 MHz)

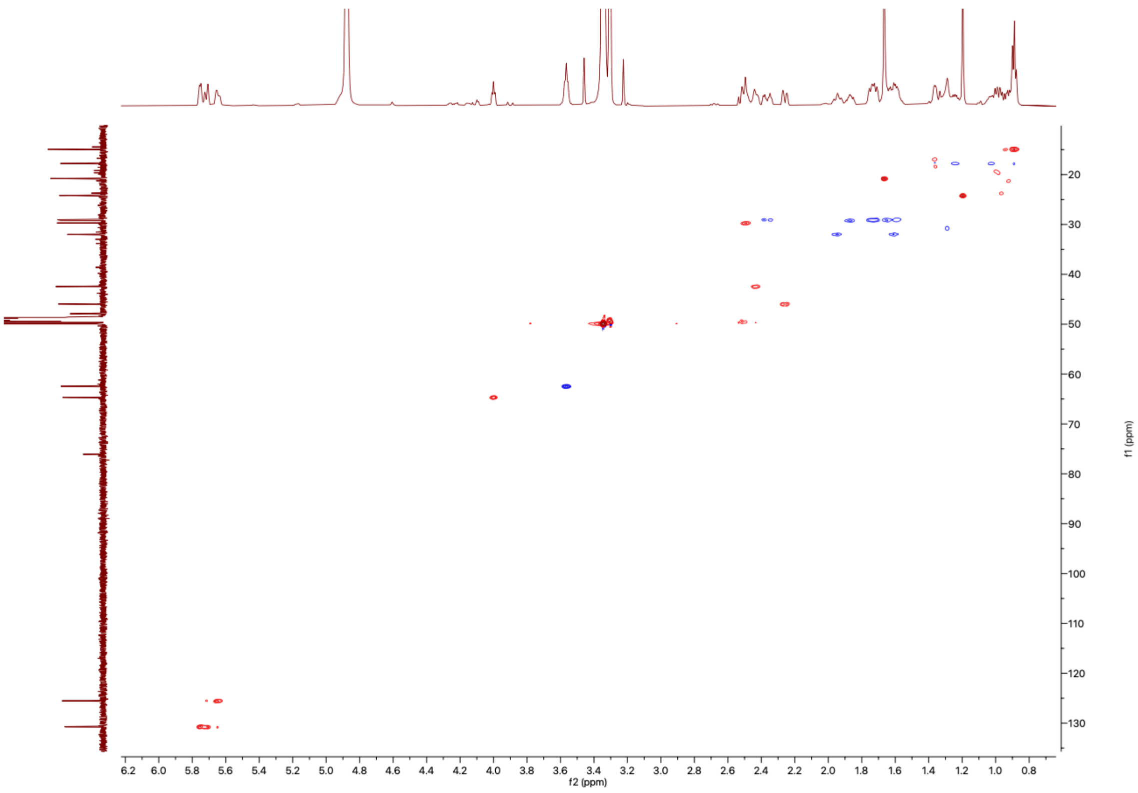

**Figure S68:** The $^1H$-$^{13}C$ HSQC NMR Spectrum of **19** in MeOD (600 MHz)

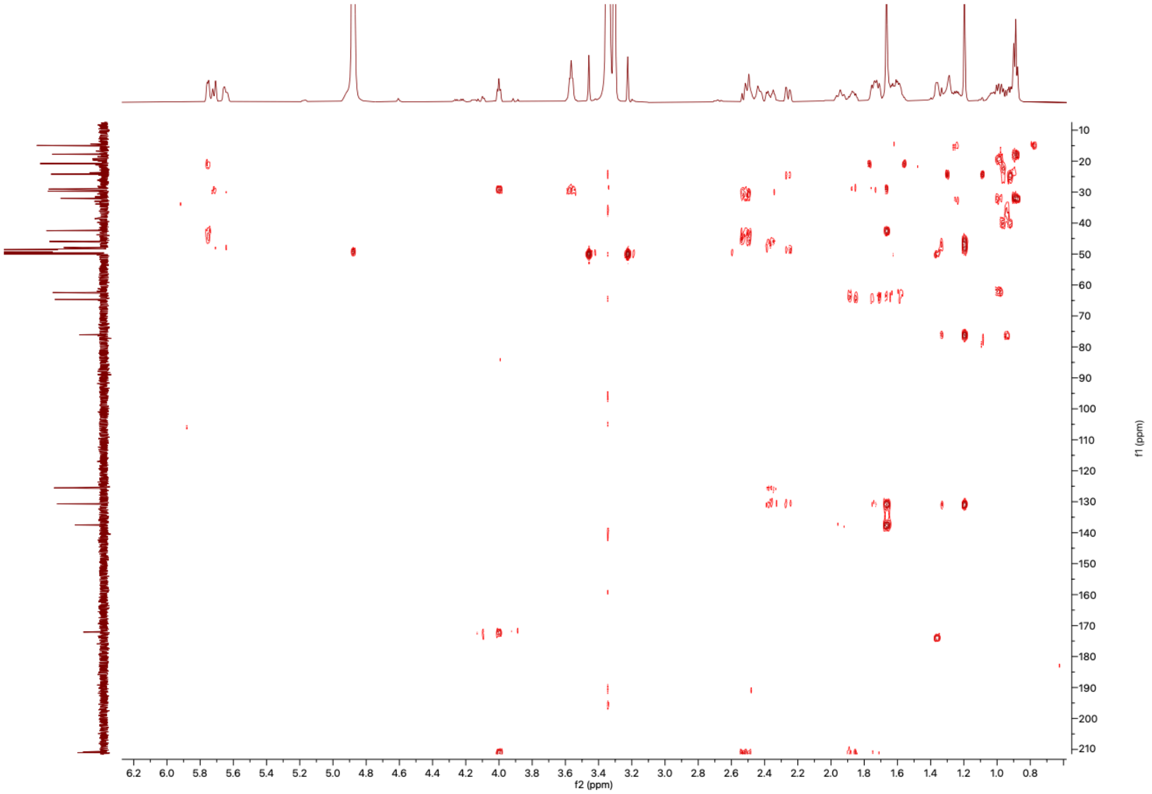

**Figure S69:** The $^{1}H$-$^{13}C$ HMBC NMR Spectrum of **19** in MeOD (600 MHz)

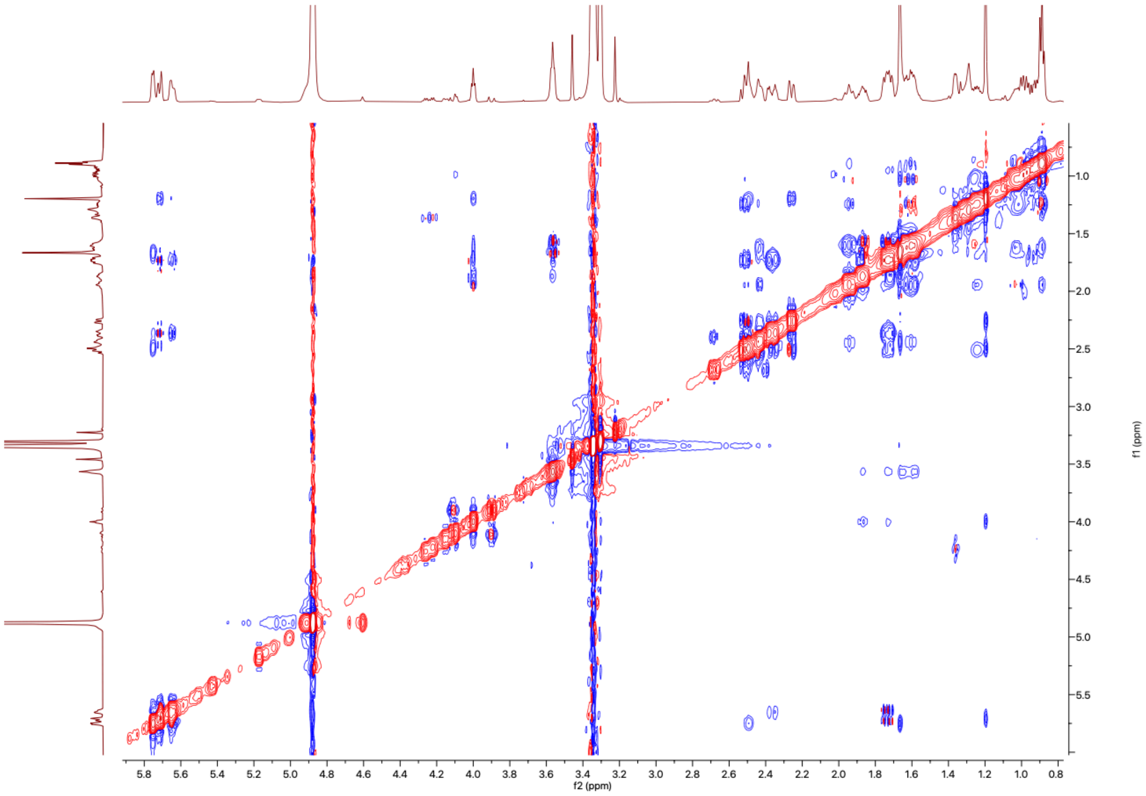

**Figure S70:** The $^{1}H$-$^{1}H$ NOESY NMR Spectrum of **19** in MeOD (600 MHz)

24. National Research Council, Division on Earth and Life Studies, Institute for Laboratory Animal Research & Committee for the Update of the Guide for the Care and Use of Laboratory Animals. *Guide for the Care and Use of Laboratory Animals: Eighth Edition*. (National Academies Press, 2011).
